# The Arabidopsis GyraseB3 contributes to transposon silencing by promoting histone deacetylation

**DOI:** 10.1101/2025.08.11.669681

**Authors:** Isabelle Gy, Sébastien Beaubiat, Nicolas Bouché

## Abstract

DNA methylation and histone modifications are key epigenetic marks controlling chromatin structure, gene expression and transposable element (TE) activity. In plants, the histone demethylase INCREASE IN BONSAI METHYLATION1 (IBM1) prevents heterochromatic silencing marks from accumulating on actively transcribed genes. Through a genetic screen of mutants defective in *IBM1* production, we identified suppressor mutations in genes essential for maintaining balanced genome-wide epigenetic states. The *gyrb3* mutation partly reversed DNA hypermethylation in *IBM1*-deficient plants, revealing a novel role for GyrB3, a nuclear protein combining domains from cyanobacterial gyrases and ELM2/SANT proteins involved in histone acetylation. In *gyrb3* mutants, TEs exhibit transcriptional activity, showing reduced DNA methylation and increased histone H3 acetylation, both of which are epigenetic marks associated with expression activation. GyrB3 physically interacts with histone deacetylases like HISTONE DEACETYLASE6 (HDA6), likely mediating their activities at TEs. The functional overlap between HDA6 and GyrB3 is further supported by the observation that, similar to *gyrb3*, a mutation in *hda6* suppresses the *Ibm2* phenotype. Our findings reinforce that histone deacetylation is essential for TE silencing and that loss of IBM1 in plants abolished the frontiers between genes and TEs, emphasizing its importance in maintaining epigenomic stability.

**Key Points:**

- IBM1 is a histone demethylase that safeguards actively transcribed genes from inappropriate heterochromatic silencing, thereby preserving their function and epigenomic integrity. A *gyrb3* mutant suppresses the developmental abnormalities observed in *ibm1* or *ibm2* mutants.
- GyrB3, though misannotated as a plant gyrase, appears to have evolved away from its conventional role and instead functions in epigenetic regulation.
- GyrB3 is a nuclear protein that interacts with histone deacetylases such as HDA6 to repress transposon activity; mutating *GYRB3* results in elevated transposon expression, driven by reduced DNA methylation and increased histone acetylation.

## INTRODUCTION

DNA methylation and histone post-translational modifications are epigenetic marks found in both plants and animals influencing chromatin structure, regulating gene expression and the activity of transposable elements (TEs). DNA methylation involves the addition of a methyl group to cytosine bases, occurring in plants at the CG, CHG, and CHH sites (where H represents A, T, or C). Modification of lysine residues on histone H3 tail can either activate or repress gene transcription, depending on which lysine residue is modified and the type of modification, such as for instance methylation (including the number of methyl groups added) or acetylation. Methylation on lysine 4 (H3K4me) or acetylation of lysine 3 (H3Kac) of histone H3 are generally associated with active transcription, while methylation of H3K9 or H3K27 are linked to gene repression and heterochromatin formation (1). Enzymes like histone methyltransferases, acetylases, demethylases, and deacetylases regulate the deposition and removal of these epigenetic marks, enabling cells to respond to various stimuli and to control developmental processes.

Histone deacetylases (HDACs) are crucial regulators of gene expression and chromatin dynamic. In yeast, animals and plants, HDACs function within multiprotein complexes to influence transcription during diverse biological processes. In Arabidopsis, 12 of the 18 HDACs are homologous to the yeast RPD3 HDAC and its animal homologs. The Arabidopsis RPD3-like HDACs are classified into four distinct classes, with HISTONE DEACETYLASE6 (HDA6) being a member of Class 1. In addition to playing a key role at genes, HDA6 also controls heterochromatin silencing by removing acetylation from histones associated with TEs (2–9). HDA6 interacts and acts synergistically with other enzymes that regulate epigenetic marks such as the CG-methylating enzyme METHYLTRANSFERASE1 (MET1) (3, 4) or the histone methyltransferases SU(VAR)3-9 HOMOLOGUE SUVH4/5/6 which catalyse the methylation of H3K9 (6). HDA6 is involved in regulating developmental stages like flowering by interacting with FLOWERING LOCUS D (FLD) and MULTICOPY SUPPRESSOR OF IRA1 4 (MSI4)/FVE to repress the expression of FLOWERING LOCUS C (FLC) (10).

The Jumonji C (JmjC) domain-containing protein INCREASE IN BONSAI METHYLATION1 (IBM1) belongs to a different class of histone modifying enzymes that remove methylation from H3K9. IBM1 prevents the deposition of heterochromatic silencing marks on actively transcribed genes (11). H3K9me2 and CHG DNA methylation are closely associated. Indeed, CHG methylation is regulated by the DNA methyltransferase CHROMOMETHYLASE3 (CMT3), which directly binds regions enriched in methylated H3K9 (12, 13). Reciprocally, H3K9me2 is catalyzed by histone methyltransferases such as SUVH4 that binds CHG-methylated cytosines (14). Thus, CMT3 and SUVH4 engage in a self-reinforcing loop between DNA and histone methylation, essential for silencing TEs and repetitive sequences but detrimental to genes in the absence of IBM1. *ibm1* mutants accumulate both H3K9me2 and CHG in coding regions, leading to severe developmental defects (11, 15). The precise mechanism by which IBM1 operates, particularly its interaction with the CMT3/KYP complex in regulating DNA methylation homeostasis at genes, remains an active area of research.

The *IBM1* gene produces two distinct transcript isoforms, with only the longest encoding a functional protein (16). *IBM1* expression is regulated by *INCREASE IN BONSAI METHYLATION2* / *ANTI-SILENCING1* / *SHOOT GROWTH1* (17–19). IBM2 controls the transcription of genes carrying intronic TEs like *IBM1* (20–24), thereby supporting critical developmental processes such as plant immunity (22, 25, 26). We previously identified *FLOWERING TIME CONTROL PROTEIN* (*FPA*) in a genetic screen as a suppressor of the *ibm2* mutation. FPA is an RNA-binding protein, which promotes the use of proximal polyadenylation sites in IBM2-targeted genes, including *IBM1*, and disease resistance gene like *RECOGNITION OF PERONOSPORA PARASITICA7* (*RPP7*). In an *fpa* mutant, only the longest functional *IBM1* transcript is generated, thereby restoring IBM1 function when *fpa* is introgressed into an *ibm2* background (22).

Here, we identified the *gyrb3* mutation as a new suppressor of both the *ibm2* and *ibm1* developmental phenotypes. *GYRB3* encodes a protein of unknown function that was misannotated as a DNA topoisomerase (27). By sequencing the methylomes of siblings obtained from crosses between *ibm2* and *gyrb3*, we revealed that the *gyrb3* mutation partially suppresses the accumulation of CHG and CHH methylation occurring in genes when *IBM2* is mutated. This correlated with transcriptional changes in an *ibm2 gyrb3* double mutant compared to *ibm2*. In a single *gyrb3* mutant, we observed a similar decrease in methylation levels more specifically in pericentromeric TEs, associated with the activation of their expression. GyrB3 is a nuclear protein that contains ELM2/SANT domains, which are characteristic of enzymes involved in regulating histone acetylation. Indeed, GyrB3 interacts with HDA6, HDA9, and HDA19. Chromatin immunoprecipitation sequencing (ChIP-seq) analysis showed that some *gyrb3* TEs accumulate H3Kac, especially those that are transcriptionally reactivated. TEs that accumulate H3Kac in *gyrb3* also accumulate H3Kac in *hda6* and are transcriptionally reactivated in both the individual *gyrb3* and *hda6* mutants. Mutation of *HDA6* in the *ibm2* background also results in suppression of the phenotype. Thus, GyrB3 contributes to the silencing of TEs by facilitating the deacetylation of histone H3, an action mediated by Class I HDAs, such as HDA6.

## MATERIALS AND METHODS

### Plant materials and growth conditions

Arabidopsis plants are in the Col-0 background. The following mutants were used: *ibm1-1* (11)*, ibm2-1* (17), *ibm2-4/sg1-1* (19) and the *axe1-5 hda6* allele, (28). The *gyrb3-2* T-DNA mutant was obtained from the Arabidopsis stock center (SAIL_390_D05). Seeds of plants grown *in vitro* were surface-sterilized and sown on Gamborg B5 medium containing 1% Sucrose. Plants were cultivated in growth chamber at 21°C in long day conditions. Leaf areas were measured with the *Easy Leaf Area* application (29).

For ethylmethane sulfonate (EMS) mutagenesis, ∼ 7,000 *ibm2-4* seeds were incubated in water containing 0.1% EMS for 15 hours at room temperature as described (22). Briefly, plants were grown in pools of 16 (440 M1 pools in total) in greenhouses in long-day conditions at 20°C. The next generation was obtained by growing 200 M2 plants per pool that were selfed to obtain the M3 generation. Whole-genome sequencing was performed on M3 plants exhibiting a wild-type phenotype, using the untreated *ibm2-4* mutant as a reference. Variant analysis was conducted using the *MutDetect* pipeline available on GitHub (https://github.com/ijpb-bioinformatics/mutdetect-pipeline).

### DNA methylation analyses

DNA was extracted with a genomic DNA extraction kit (*Macherey-Nagel*) from the aerial parts of seedlings collected in bulks (n=30) after 15 days of *in vitro* growth under long day conditions. Bisulfite treatment, library preparation and whole-genome sequencing (depth of 38X) were performed at *BGI* using the DNBSEQ technology producing 100 bp paired-end reads (Supplementary Table S1). Raw reads were cleaned and trimmed with *Trim_Galore* v0.6.5 (*Babraham Bioinformatics*). Reads were aligned to the Col-0 *Arabidopsis thaliana* TAIR10.57 reference genome with *Bismark* version v0.24.0 (*Babraham Bioinformatics*) and standard options (*Bowtie2*; 1 mismatch allowed). Identical pairs were collapsed. Differently Methylated Regions (DMRs) were identified using *DMRcaller* (30) by setting the *minProportionDifference* option to 0.4 for CG, 0.2 for CHG and 0.1 for CHH. Metaprofiles were drawn using *deepTools* v3.5.2 (31) and circular graphs using *Circlize* v0.4.16 (32).

To define regions enriched for TEs or genes, the repeats and CDS densities were determined in 500 kb windows covering the whole genome, then the windows were clustered using the k-means method in R (*kmeans*), allowing the distinction of three types of genomic regions (repeat-rich, repeat-poor, and a third intermediate category; Supplementary Table S2) as described before (33). Repeats or genes that span across two regions were excluded.

To examine the overlaps between DMRs of *gyrb3* and *hda6*, raw read BS-seq *hda6* files were retrieved from the GEO repository, accession number GSE216787 (8). To monitor the methylation levels in other epigenetic mutants BS-seq raw read files were retrieved from the GEO repository, accession number GSE169497 for *met1-9* and the *drm1 drm2 cmt3 cmt2* (*ddcc*) quadruple mutant (34) and PRJNA694533 for *ros1-1* (35). The overlaps between DMRs were obtained with the “intersect” command from the *BEDTools* suite.

### RNA-seq analyses

Total RNA was isolated from the aerial parts of seedlings collected in bulks (n=30) after 15 days of *in vitro* growth under long day conditions using the *RNeasy Plant Mini* kit (*Qiagen*) followed by a DNAse I treatment (*Fermentas*). Three biological replicates were sequenced per genotype. Strand specific library preparation and sequencing (DNBSEQ technology) were performed by the *BGI*. On average, 24 million paired-end 100 bp clean reads were obtained. The nf-core *Rnaseq* v3.17.0 pipeline and Nextflow v24.04.2 were used for trimming (*Trim_Galore*) and aligning (*STAR*) the reads to the TAIR10.57 version of the Arabidopsis genome and then to quantify the transcripts (*Salmon*).

The differential analysis was conducted using the obtained count matrix and the transcript lengths, using the nf-core *Differential Abundance* pipeline v1.5.0 and Nextflow v24.04.2. This pipeline is built on the R package *DESeq2* (36). Z-score were calculated on normalized counts (*DESeq2*) using the *scale* function of R with the list of genes that were significantly up- or downregulated (log2FC ≤ -1 or ≥ 1; p-value < 0.05; based on *DESeq2*) in at least one genotype. Z-score hierarchical clustering was obtained with the *hclust* function of R. Clustered heatmaps were drawn with the R package *Pheatmap* v1.0.12. Co-expression analyses were performed using Gaussian mixture models available in the *coseq* R package v1.32.1 (37) and the enrichment analysis was performed using the *clusterProfiler* R package v4.16.0 (38) with the Bioconductor *RFLOMICS* package.

### Chromatin immunoprecipitation (ChIP) sequencing

ChIP experiments were performed on 1 g of aerial parts of 15-days old seedlings grown *in vitro* under long day conditions (two biological replicates per genotype). Plants were crosslinked with 1% formaldehyde solution supplemented with 1 mM PMSF to avoid protein degradation. Crosslinked chromatin was sonicated for 22 cycles at medium intensity (30s ON/30s OFF) followed by 15 cycles at high intensity (30s ON/30s OFF) on a Diagenode Bioruptor. ChIP was performed overnight using Protein A Dynabeads (*ThermoFisher*) and anti-acetyl-histone H3 antibody (Sigma-Aldrich; Ref 06-599). Precipitated DNA was eluted and purified using phenol:chloroform (Sigma-Aldrich; Ref 77617). Chromatin untreated (without IP) and sonicated was processed in parallel and used as the input. Libraries and sequencings were performed at *BGI* using the DNBSEQ technology producing 50 bp single-end reads, with an average of 25 million per sample. Sequencing of both input and IP samples was carried out by the *BGI* company (China).

### ChIP-seq data analysis

Analyses were done using the nf-core *Chipseq* pipeline v2.1.0 and Nextflow v24.04.2. Reads were trimmed (*Trim_Galore*) and clean reads were aligned (*BWA*) to the TAIR10.57 version of the Arabidopsis genome. Only uniquely mapped reads were retained. ChIP-seq peaks were called using MACS v3.0.1 (39). Differential H3Kac sites were identified using *MAnorm2* (40). Sites with adjusted *p*-value below 0.05 and fold change greater than 2 were defined as differential ones. To monitor the H3Kac signal in *hda6* ChIP experiments, raw read files were retrieved from the GEO NCBI data repository. The data described in (41) consist of five biological replicates for the wild-type, and two replicates each for *hda6* and *hda19* (accession number GSE166090). The data described in (42) consist of three biological replicates for the wild-type, three for *hda6* plus their inputs (accession number GSE167288). The data described in (43) consist of two biological replicates for the wild-type and two for *hda6* (accession number GSE132636). Metaplots were drawn with *deepTools* v3.5.2 (31). ChIP signals were visualized with *JBrowse2* v3.6.4 using normalised *BigWig* files scaled to 1 million mapped reads. Peak overlaps were obtained using the *bed_intersect* function of the *valr* R package v0.8.4 and intersections were visualized using the *UpSetR* R package v1.4.0.

### Protein structure predictions

AlphaFold2-Multimer predictions were obtained with *ColabFold* v1.5.5 (44) using standard options. Five distinct models were created, each undergoing three rounds of refinement. The ipTM values were calculated based on the averages of those obtained in the *ColabFold* log file. The predicted HDA6/GyrB3 protein complex was visualized with *Chimera* v1.9.

### Yeast two-hybrid assay

Full-length *GYRB3*, *HDA6*, *HDA7*, *HDA9* and *HDA19* cDNAs were synthetized (*TWIST Bioscience*) and cloned in a *pTwist ENTR* vector. The entry vectors were used to generate the appropriate *pGAD* and *pGBK* yeast two-hybrid expression vectors, using the Gateway technology (*ThermoFisher*). The integrity of the sequences was confirmed by sequencing the vectors carrying the constructs. Yeast two hybrid assays were carried out using the GAL4-based system (*Clontech*) by introducing plasmids harboring the gene of interest in yeast strains AH109 and Y187 by lithium acetate transformation. After mating of haploid yeasts on YPD plates, diploid cells expressing Gal4-BD and Gal4-AD fusion proteins were selected in SD/- LW, a dropout medium without leucine and tryptophan. Protein interactions were assayed by growing diploid cells in serial dilutions for 5 days at 30°C on media lacking leucine, tryptophan and histidine (SD/-LWH). Two proteins were deemed to interact when the spots grew on SD/- LWH. Assays were repeated twice.

### GyrB3 nuclear localization

To examine the subcellular localization of the GyrB3 protein, two constructs were generated: *pGWB5-GYRB3* (a GyrB3-GFP fusion driven by the *35S* promoter) and *pGWB6-GYRB3* (a GFP-GyrB3 fusion under the same promoter). Constructs were infiltrated into transgenic *Nicotiana benthamiana* leaf epidermal cells expressing a nuclear CFP marker fused to histone 2B as nuclear labeling control (45). Transient expression assays were performed as described (46). To enhance transient expression, the P19 viral suppressor of gene silencing was co-expressed (47) by infiltrating a *p35S::P19* construct. Confocal imaging was conducted using a Zeiss LSM710 laser-scanning confocal microscope, with GFP detected using 488 nm excitation and a 501-536 nm emission range, and CFP detected using 458 nm excitation and a 467-497 nm emission range. To confirm the authenticity of the GFP signal, the fluorescence emission spectrum of the sample was recorded from 494 to 728 nm in 9.7 nm intervals under 488 nm excitation.

## RESULTS

### Mutating *GYRB3* suppresses the developmental phenotypes of *ibm2* or *ibm1*

The developmental phenotype of *ibm2* mutants is similar to that of *ibm1* mutants, resulting from a deficiency in producing a long functional *IBM1* mRNA (17–19). A mutant screen was conducted on *ibm2-4* to identify individuals exhibiting a wild-type phenotype, with the goal of selecting genetic suppressors of *ibm2* or *ibm1* (22). Approximately 7,000 *ibm2-4* seeds were treated with EMS and the genetic screen was performed on M3 seedlings (22). We sequenced the whole-genome of M3 plants displaying a wild-type phenotype to compare them to the untreated *ibm2-4* mutant background. Among the various homozygous mutations identified, several were found in coding regions of genes involved in DNA and H3K9 methylation pathways. The *cmt3* or *kyp* mutations suppress *ibm1* by preventing the accumulation of heterochromatic marks on a large range of IBM1 target genes (11). Homozygous mutations corresponding to yet undescribed alleles of both *cmt3* (n=7) and *kyp* (n=3) were identified (Supplementary Table S3). In addition, a *histone demethylase lsd-like* 2 (*ldl2*) allele was detected within our suppressors (Supplementary Table S3). *LDL2* encodes a homolog of H3K4me2 demethylases and its mutation in an *ibm1* background restores the expression patterns of numerous genes and suppress the phenotypes of *ibm1* (48). Interestingly, the *ldl2* allele identified in our analysis results in an identical amino acid alteration to that which was reported for *ldl2-12* (48). In addition to the *fpa* mutant previously described (22), the finding of *cmt3*, *kyp* and *ldl2* alleles reinforced our strategy in identifying *ibm1* suppressors.

Five additional suppressors, harbouring homozygous mutations in the coding regions of genes unrelated to the CMT3/KYP/IBM1 pathway, were identified. One of these suppressors exhibited only two of these mutations, including a deletion of a C (Chr5:1,111,465) in the last exon of *GYRB3* (*AT5G04110*), creating a premature stop codon at that position. The corresponding allele was designated *gyrb3-1* (Fig. 1A). *GYRB3* encodes GyrB3, a protein of unknown function that was likely misannotated as a type II DNA topoisomerase (27). Based on data from the *InterPro* database (49), GyrB3 contains three distinct domains involved in DNA/chromatin binding: Topoisomerase II, ELM2, and SANT (Fig. 1A), making it a good candidate for further investigations.

**Fig. 1.**
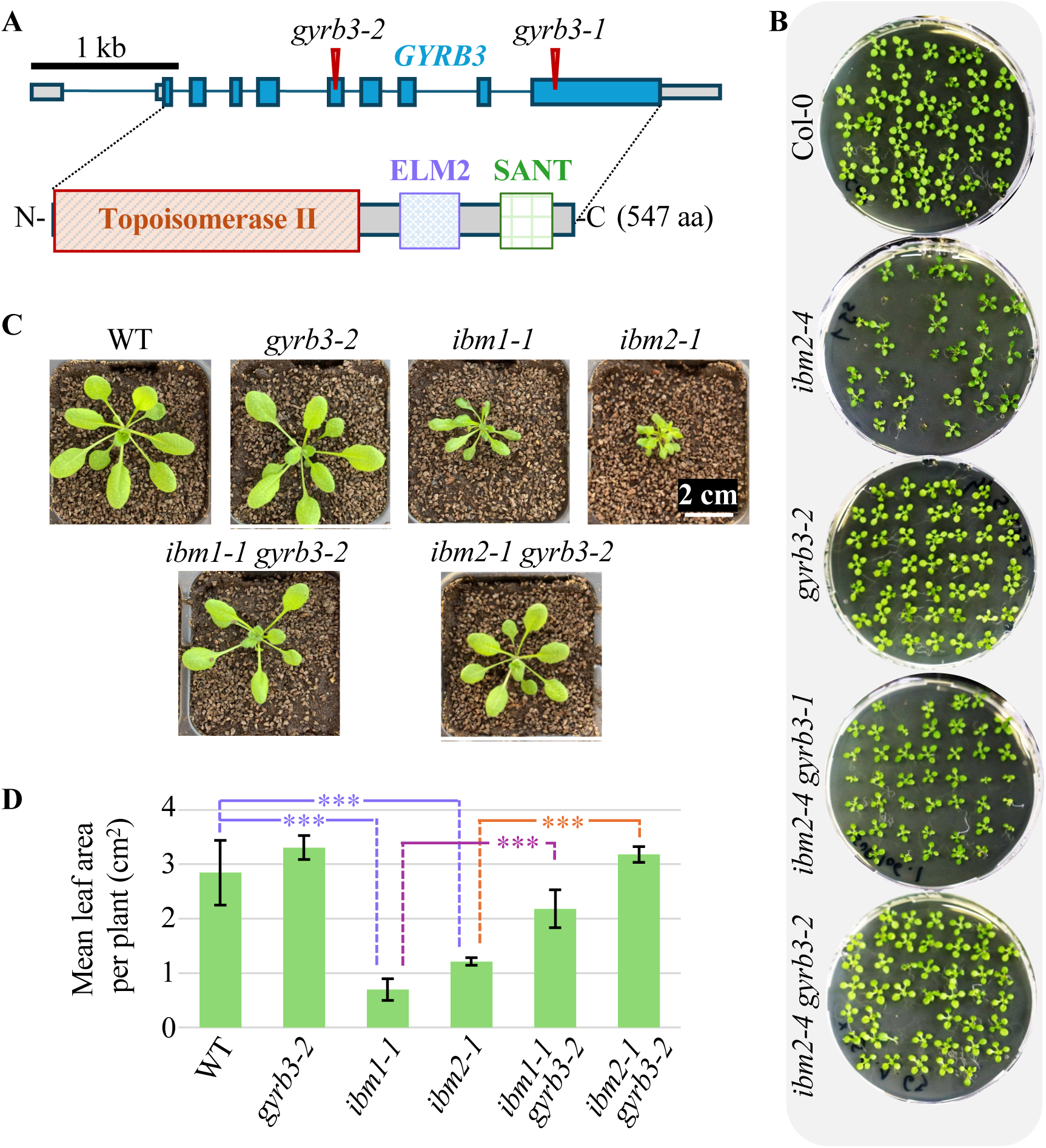
The developmental phenotypes of *ibm2* are suppressed when *GYRB3* is mutated. (A) Structure of the *GYRB3* gene (At5g04110) showing untranslated regions (UTRs, grey boxes), exons (blue boxes), and introns (black lines). The positions of two mutations are indicated by red triangles: *gyrb3-1* a cytosine deletion at Chr5:1,111,465, and *gyrb3-2*, a T-DNA insertion (SAIL_390_D05). The GyrB3 protein (UniProtKB: Q8VY11) is shown below, with annotated domains: Topoisomerase II (amino acids 2-320), ELM2 (361-422) and SANT (469-517). (B) The *gyrb3-1* and *gyrb3-2* mutations suppress the developmental abnormalities observed in *ibm2* mutants such as small, distorted leaves and reduced germination rate. Plants were pictured 15 days after sowing. The experiment was repeated five times; only one repeat is presented, the others are shown in Supplementary Fig. S1. (C) Pictures of *gyrb3-2, ibm1-1*, *ibm2-1*, *ibm1-1 gyrb3-2* and *ibm2-1 gyrb3-2* plants taken 21 days after sowing. The *gyrb3-2, ibm1-1*, *ibm1-1 gyrb3-2* mutants and wild-type (WT) control lines were obtained by self-pollinating F2 progenies from a cross between *ibm1-1* and *gyrb3-2*. The *ibm2-1*, and *ibm2-1 gyrb3-2* mutants were obtained by self-pollinating F2 progenies from a cross between *ibm2-1* and *gyrb3-2*. Five plants per genotype were grown together and only one repeat is presented, the others are shown in Supplementary Fig. S2. (D) Mean leaf areas per plants shown in (C) and Supplementary Fig. S2, measured with the *Easy Leaf Area* application. Error bars indicate standard deviation. A t-test was employed as the statistical method to compare leaf area. ***, p-value < 0.01; the test for all other combinations is non-significant.

To confirm that mutating *GYRB3* suppresses the phenotypes of *ibm2*, a *gyrb3* T-DNA homozygous mutant (*SAIL_*390_D05, referred to as *gyrb3-2*) was crossed with *ibm2-4*. Like the original *ibm2-4 gyrb3-1* suppressor (Fig. 1B and Supplementary Fig. S1), the phenotypes of *ibm2* plants were rescued in the *ibm2-4 gyrb3-2* double mutant (Fig. 1B and Supplementary Fig. S1). *gyrb3-2* was also crossed with a different *ibm2* allele, *ibm2-1*, and their F3 segregating progenies were analysed. Again, the phenotypes of *ibm2* plants were rescued in the double *ibm2-1 gyrb3-2* mutant (Fig. 1C and Supplementary Fig. S2), as confirmed by measuring leaf area (Fig. 1D).

To test if a mutation in *GYRB3* also suppresses the phenotypes of *ibm1*, *gyrb3-2* was crossed with an *ibm1-1* homozygous mutant. In the segregating F3 progeny, plants that carry only the *gyrb3-2* mutation developed like plants carrying both the *IBM1* and the *GYRB3* wild-type alleles (Fig. 1C and Supplementary Fig. S2). Plants homozygous for *ibm1-1* presented severe developmental anomalies, while the double *gyrb3-2 ibm1-1* mutant developed better (Fig. 1C and Supplementary Fig. S2), as confirmed by measuring leaf area (Fig. 1D). Thus, a mutation in *GYRB3* is likely epistatic over an *ibm2* or an *ibm1* mutation.

### The *gyrb3* mutation partially reverses the DNA hypermethylation that occurs in genes when IBM1 is absent

Mutations in either the *IBM1* or *IBM2* gene result in the DNA hypermethylation of many genes in both the CHG and CHH contexts (17–19), resulting from the CMT3/KYP feedback loop acting on genes instead of TEs as observed in the wild-type. To determine whether mutating *GYRB3* modifies the way genes are hypermethylated in *ibm2* or *ibm1*, we sequenced the methylomes of F3 siblings obtained by crossing *ibm2-1* and *gyrb3-2* (Fig. 1C). Genomic DNA was extracted from bulks of seedlings carrying solely the *ibm2-1* allele or both *ibm2-1* and *gyrb3-2*. To compare their methylomes, whole genomes were sequenced after bisulfite treatment (WGBS; Supplementary Table S1) and the patterns of methylation were determined for both genes and TEs. CG methylation metaprofiles for genes or TEs were unchanged between the Col-0 control, *ibm2-1* and *ibm2-1 gyrb3-2* (Fig. 2), however, the CHG hypermethylation monitored in genes of *ibm2-1* seedlings was reduced by 15% (in repeat-poor regions) to 24% (in repeat-rich regions) and the CHH methylation was reduced by 7% (in repeat-poor regions) to 29 % (in repeat-rich regions) in their *ibm2-1 gyrb3-2* siblings (Fig. 2, left panel). The results show that the *gyrb3* mutation partially suppresses the CHG and CHH methylation accumulating in genes of *ibm2*. Surprisingly, a reduction in both CHG and CHH methylation (21% and 19%, respectively) was also observed in TEs located in repeat-rich regions of *ibm2-1 gyrb3-2*, compared to *ibm2* or Col-0 (Fig. 2, right panel).

**Fig. 2.**
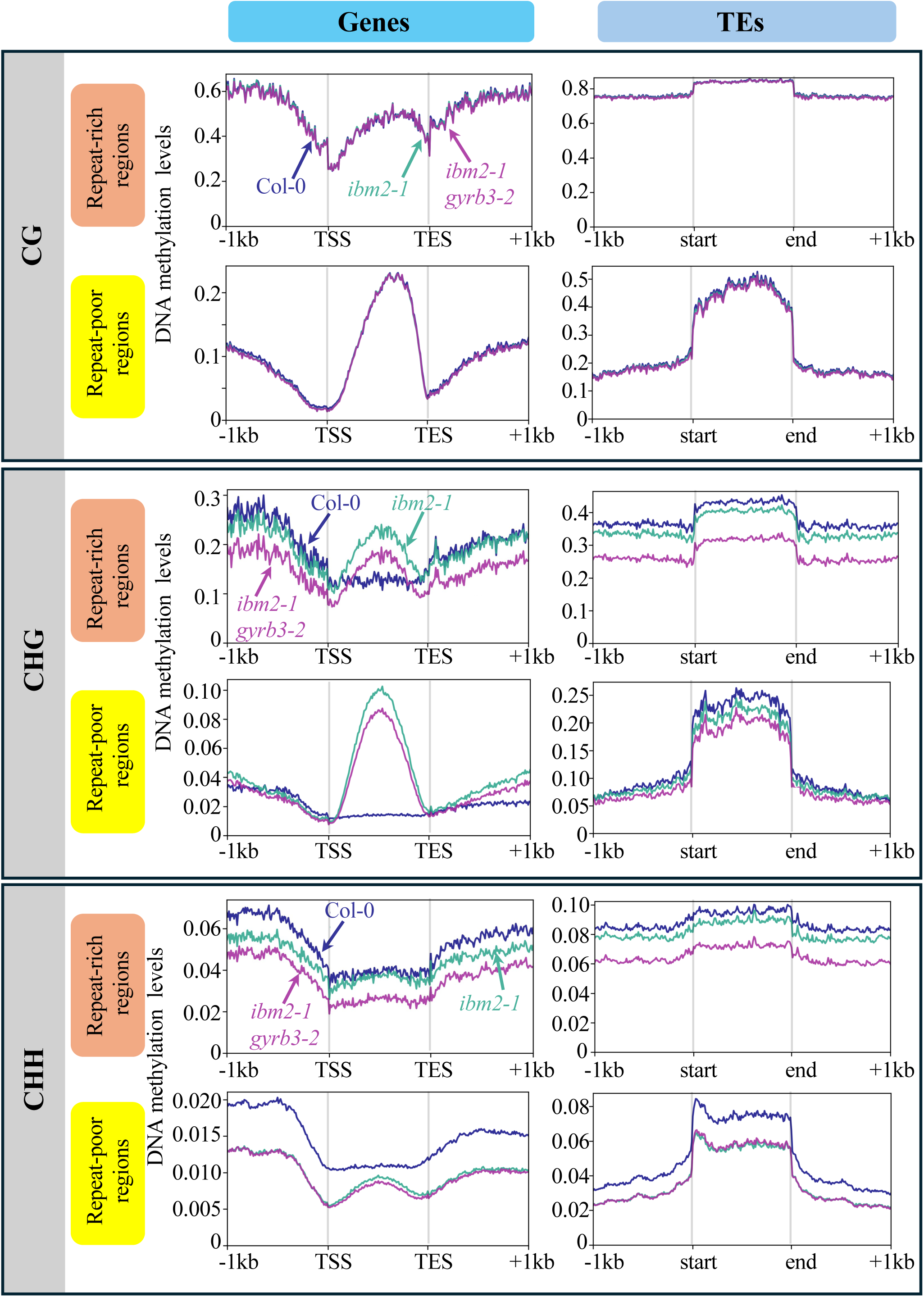
Methylation patterns of genes and TEs in the *ibm2-1* and *ibm2-1 gyrb3-2* mutants. The average methylation levels of genes (left panels) and TEs (right panels) were calculated by segmenting the corresponding annotated regions (TAIR10.57 release) into 100-bp bins. The metaplot analyses also include regions located 1 kb upstream and 1 kb downstream of the gene bodies and TEs. Repeat/TE-rich and repeat-poor (i.e. gene-rich) regions were defined as described in the *Materials and Methods* section. The methylation patterns in Col-0 are shown as controls.

### TEs are hypomethylated and transcriptionally reactivated in a *gyrb3* mutant

To understand why TEs were hypomethylated in the double *ibm2-1 gyrb3* mutant compared to *ibm2*, we sequenced the methylome of the single *gyrb3* mutant (Supplementary Table S1). Average levels of methylation were calculated in 100 bp-tiles partitioning the genomes. The CG methylation was unchanged between the Col-0 control and the *gyrb3-2* mutant (Fig. 3A). However, we observed a significant decrease (t-test; p<0.001) in the general levels of methylation in both the CHG and the CHH contexts that was more pronounced in repeat-rich regions compared to gene-rich regions (Fig. 3A). The methylation metaprofiles of TEs confirmed that changes occurred in both the CHG and CHH contexts (Fig. 3B) but not in the CG context (Supplementary Fig. S3). Notably, the levels of methylation for TEs localized in repeat-rich regions were reduced by 19% for CHGs and 16.5% for CHHs between Col-0 and the *gyrb3-2* mutant (Fig. 3B and Supplementary Fig. S3). We confirmed the results by identifying the Differentially Methylated Regions (DMRs) in *gyrb3-2* compared to Col-0 (Fig. 3C). Indeed, the highest number of DMRs was observed for CHG hypoDMRs (n= 5 557), most of them being localized in regions enriched for TEs (Fig. 3C). These DMRs are localised in pericentromeric regions (Fig. 3D) and mainly overlap with TEs (Fig. 3E). Altogether, the results show that some TEs are hypomethylated in a *gyrb3* background compared to the Col-0 control. Therefore, the non-CG hypomethylation of TEs detected in the double *ibm2 gyrb3* mutant (Fig. 2) is likely a consequence of the non-functionality of *GYRB3*.

**Fig. 3.**
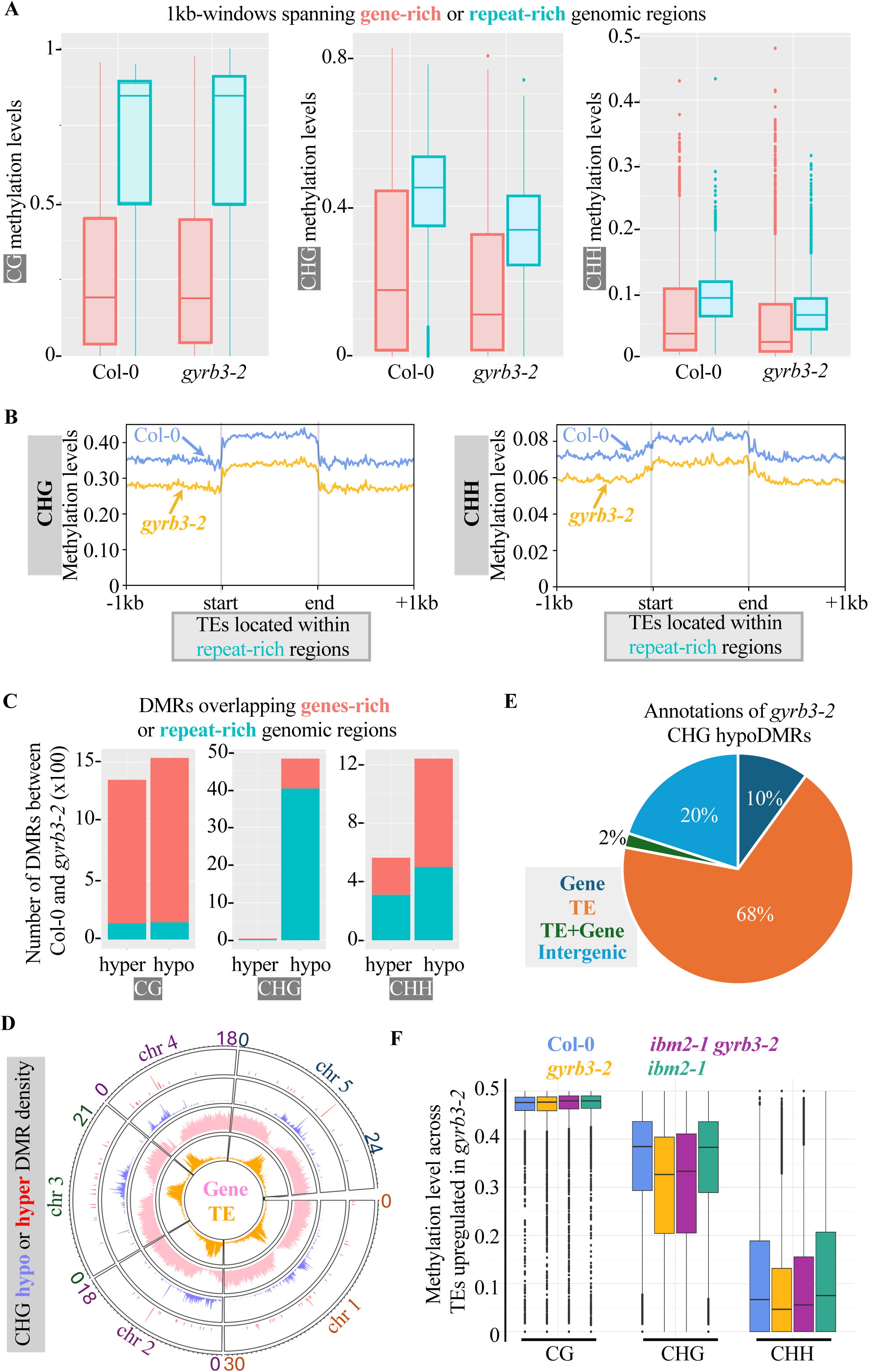
The *gyrb3* methylome shows alterations compared to the Col-0 control. (A) Boxplots showing mean methylation content of the *gyrb3* mutant and the Col-0 control in genomic regions enriched in TEs (*repeat-rich*) or genes (*gene-rich*). The Arabidopsis genome (TAIR10.57 release) was partitioned in 1 kb-tiles and methylation levels correspond to the ratios of methylated cytosines over the total number of cytosines. Only cytosines covered by at least five reads were considered. The average methylation levels were calculated by combining the two biological replicates for each genotype. (B) Metaprofiles of TEs localized in repeat-rich regions of the Col-0 control and the *gyrb3-2* mutant. Patterns of CHG and CHH methylation are presented. The complete list of metaprofiles for TEs and genes is shown in Supplementary Fig. S3. The average methylation levels of TEs were determined by dividing the corresponding annotated regions (TAIR10.57 release) into 100-bp bins. The metaplot analyses also include regions located 1 kb upstream and 1 kb downstream of the gene bodies and TEs. Repeat/TE-rich and repeat-poor (i.e. gene-rich) regions were defined a described in the *Materials and Methods* section. (C) Number of DMRs found between Col-0 and *gyrb3-2* in regions enriched for genes (red) or TEs (blue). Regions (Supplementary Table S2) were defined as described in the *Materials and Methods* section. DMRs overlapping regions with intermediate levels of TEs/genes (Supplementary Table S2) are excluded from the graph. (D) Circular plot showing the distribution of CHG hypo- (blue) or hypermethylated (red) DMRs found in *gyrb3-2* across the five chromosomes of Arabidopsis. The numbers of DMRs, TEs (i.e. Gypsy elements) and genes contained within bins of 200-bp are plotted in the center. The chromosome scale is given in Mb. Regions were considered differentially methylated when the absolute differences in methylation were at least 20% for mCHG, between *gyrb3-2* and the Col-0 control. (E) Annotation of CHG hypoDMRs upregulated peaks in *gyrb3-2*. “gene+TE” corresponds to DMRs overlapping with both genes and TEs, “gene” to DMRs overlapping with genes, and “TE” to DMRs overlapping with TEs. All other DMRs were classified as “Intergenic”. (F) DNA methylation levels in TEs significantly (log2FC ≥ 1, p-value < 0.05) upregulated in *gyrb3-2* (n=36).

To investigate whether the hypomethylation of TEs in *gyrb3* impacts their expression, an RNA-seq analysis was performed on seedlings of the *gyrb3-2*, *ibm2-4* and *ibm2-4 gyrb3-2* mutants, compared to Col-0, using three biological replicates for each genotype. 36 TEs were upregulated in *gyrb3* (Table 1 and Supplementary Table S4), and none were downregulated, indicating that the expression of a subset of TEs is released in the *gyrb3* mutant. 78% (n=28) of these 36 TEs overlap with regions hypomethylated in the CHG context in *gyrb3-2* compared to Col-0, and 44% (n=16) with regions hypomethylated in CHH (Supplementary Table S4). Thus, the transcriptional reactivation of TEs in *gyrb3* is associated with a decrease of methylation in both the CHG and CHH contexts. This was confirmed by comparing the methylation levels of all 36 *gyrb3-*upregulated TEs in the *gyrb3-2*, *ibm2-4* and *ibm2-4 gyrb3-2* backgrounds (Fig. 3F). The TEs reactivated in *gyrb3* were overall less methylated in the CHG and CHH contexts in *gyrb3* and *ibm2-4 gyrb3-2*, compared to Col-0 or *ibm2-4* (Fig. 3F). Accordingly, in the wild-type, GyrB3 is involved in silencing TEs, and mutating *GYRB3* leads to the transcriptional reactivation of some of them.

**Table 1.** Number of genes and TEs significantly (p-value < 0.05) upregulated (log2FC ≥ 1) or downregulated (log2FC ≤ -1) between the genotypes indicated. RNA-seq analysis was conducted with three biological replicates.

|  | Genes |  | TEs |  |
| --- | --- | --- | --- | --- |
|  | up | down | up | down |
| *gyrb3-2* versus Col-0 | 68 | 48 | 36 | 0 |
| *ibm2-4 gyrb3-2* versus Col-0 | 294 | 341 | 62 | 25 |
| *ibm2-4* versus Col-0 | 298 | 119 | 32 | 33 |
| *gyrb3-2* versus *ibm2-4* | 119 | 224 | 53 | 28 |
| *ibm2-4 gyrb3-2* versus *ibm2-4* | 70 | 184 | 25 | 5 |

### A *gyrb3* mutation partially compensates *ibm2*-mediated transcriptomic alterations

In the *ibm* mutants, many genes exhibit similarities with TEs by accumulating both H3K9me2 and CHG DNA methylation in their coding regions leading to transcriptome changes. We hypothesized that mutating *GYRB3* in either an *ibm2* or *ibm1* background could prevent this accumulation of methylation, thereby restoring transcription. Genes that were significantly down- or upregulated (log2FC ≤ -1 or ≥ 1; p-value < 0.05) relative to the Col-0 control were identified through RNA-seq analysis (Table 1 and Supplementary Table S5). Following log-normalized Z-score transformation of gene expression counts, hierarchical clustering of genes differentially expressed in at least one genotype was performed. The resulting heatmap (Supplementary Fig. S4) showed that biological replicates clustered consistently within each genotype. None of the genes involved in regulating non-CG methylation like *ROS1*, *DEMETER*, *CMT3*, *SUVH4*, *SUVH5*, *SUVH6*, and *IBM1* are differentially expressed in *gyrb3*. We performed a co-expression analysis by clustering differentially expressed genes according to their average expression profile in all samples. We used the *Coexpression_coseq* function with its default settings to cluster the genes from the combined lists of all differentially expressed genes. Cluster 1 comprises genes (n=168) that show similar expression in Col-0, *gyrb3-2* and *ibm2-4 gyrb3-2* but exhibit higher expression in *ibm2-4* (Fig. 4A). Cluster 2 includes genes (n=368) that show higher expression in all mutants compared to Col-0, whereas cluster 3 comprises genes (n=373) with lower expression levels in all mutants relative to Col-0 (Fig. 4A). Among all gene clusters, only cluster 1 shows unique expression changes in *ibm2* compared to other genotypes. Gene ontology analysis indicated that cluster 1 is significantly enriched in terms associated with plant pathogen defense (Supplementary Fig. S5), consistent with previous findings that *IBM2* targets resistance genes such as *RPP7* or *RPP4* (22).

**Fig. 4.**
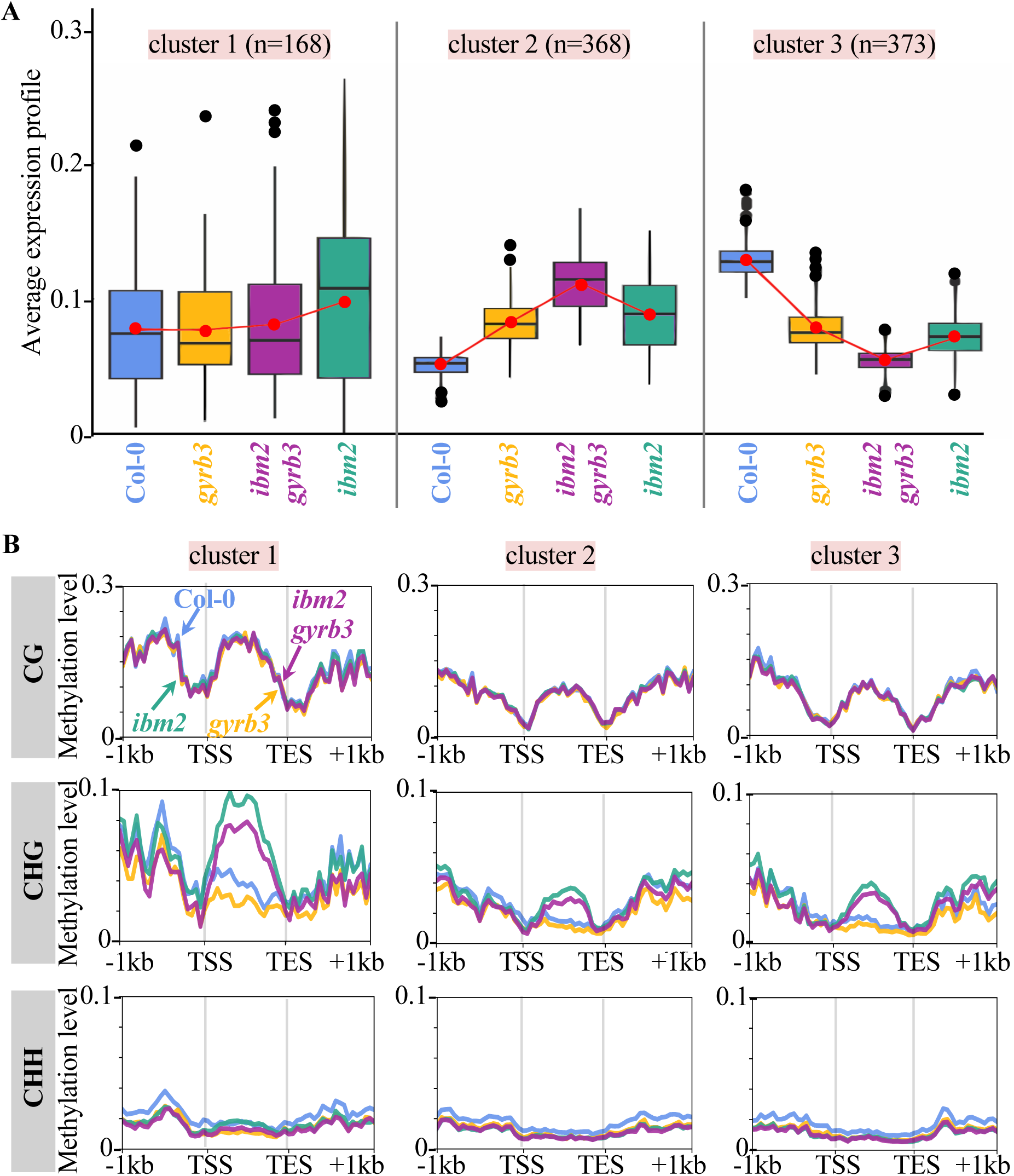
Transcriptional changes occurring when *GYRB3* is mutated in *ibm2*. (A) Gene expression profiles were calculated as the normalized expression values divided by the mean normalized expression across all conditions. Replicates were collapsed by averaging within each condition. The optimal number of clusters, determined using the Integrated Completed Likelihood (ICL) criterion, was three. (B) Metaprofiles of DNA methylation levels across genes belonging to the three clusters identified in (A).

The methylation patterns of the genes contained within each cluster were monitored across the different mutant backgrounds by generating metaprofiles (Fig. 4B). No differences in CG or CHH methylation were detected among the genotypes within the same cluster. In the *ibm2* mutant, genes with high CG gene body methylation tend to have increased ectopic CHG methylation. Indeed, genes of cluster 1 that exhibit the highest levels of CG methylation also show the highest levels of CHG methylation in *ibm2*. Consistent with the genome-wide reduction of CHG methylation observed in *gyrb3* (Fig. 2 and 3), CHG methylation within gene bodies is reduced in the *ibm2 gyrb3* double mutant compared to *ibm2* alone. This effect is particularly pronounced in cluster 1, which contains genes that are the most affected by the *ibm2* mutation among all clusters. Altogether, the results indicate that the transcriptional alterations caused by the *ibm2* mutation can be partially suppressed by mutating *GYRB3* and that this suppression is associated with methylation changes occurring in *ibm* mutant backgrounds.

### GyrB3 contributes to the silencing of transposons through histone deacetylation

GyrB3 is annotated as a type II DNA topoisomerase, still it misses crucial motifs usually found in these proteins, and contrarily to other GyrB Arabidopsis proteins, GyrB3 cannot complement a bacterial *gyrB* strain. These observations led other groups to conclude that GyrB3 is likely not a gyrase subunit (27). BLASTP comparisons with all Arabidopsis proteins revealed that GyrB3 shares domains with enzymes involved in the control of histone acetylation, like ARID1 (50) or ELM2/SANT-containing proteins (41) (Supplementary Fig. S6). In Arabidopsis, proteins containing ELM2/SANT domains play crucial roles in gene expression regulation and plant stress responses, particularly through histone acetylation (41, 42, 51). We therefore investigated whether GyrB3 might contribute to the deposition of H3Kac marks. To this end, chromatin immunoprecipitation sequencing (ChIP-seq) analysis was employed to investigate the distribution of H3Kac in aerial parts of *gyrb3-2* seedlings after 15 days of *in vitro* growth. No significant differences were found in the number and length of H3Kac ChIP peaks at the genome-wide level (Supplementary Fig. S7). We used the MAnorm2 model (40) to normalize and quantify the ChIP signals across the Col-0 and *gyrb3-2* samples and to detect differences of H3Kac accumulation in a locus-specific manner (Fig. 5A). 74 loci showed significant enrichment (log2FC ≥ 1; p-value < 0.05) for H3Kac in *gyrb3-2* compared to Col-0, mainly overlapping with TEs (n=43; Fig. 5B). Only one peak showed significantly reduced H3Kac levels (log2FC ≤ -1; p-value < 0.05) in *gyrb3-2* compared to Col-0 (Fig. 5A). Of the 36 transcriptionally upregulated TEs in *gyrb3-2*, 20 overlapped with regions gaining H3Kac in *gyrb3-2* compared to Col-0. Additionally, in all regions where H3Kac increased in *gyrb3-2*, DNA methylation levels were lower in the mutant compared to Col-0 (Fig. 5C). Collectively, these results suggest that the absence of GyrB3 leads to both the reduction of DNA methylation levels and the accumulation of histone acetylation at specific TEs, releasing their expression in the *gyrb3* mutants.

**Fig. 5.**
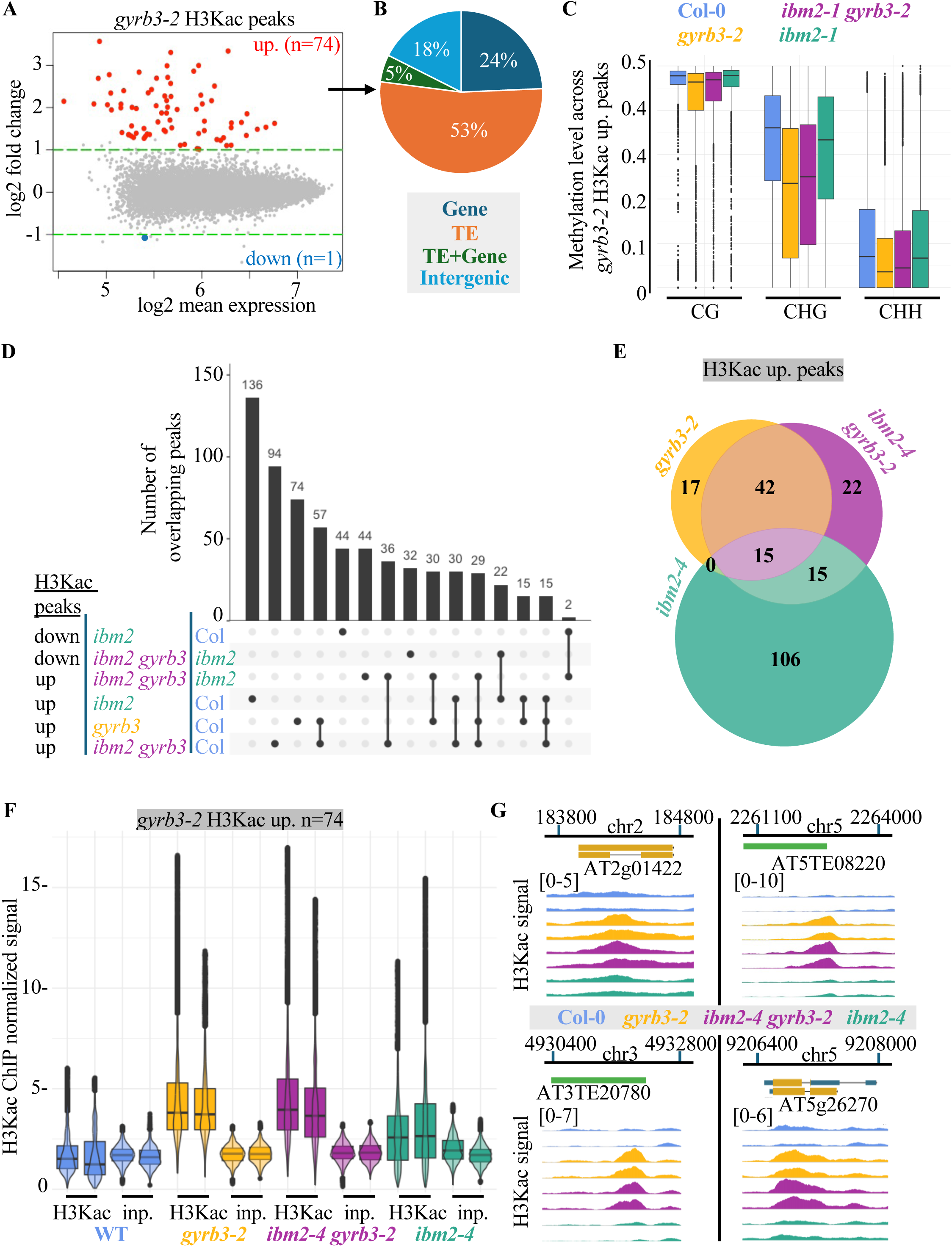
Levels of H3Kac are impaired in a *gyrb3* mutant. (A) MA plots for H3Kac peaks in Col-0 and *gyrb3-2*. Significantly differentially accumulated peaks (log2FC ≤ -1 or ≥ 1; p-value < 0.05) are indicated by red dots and distinguished from non-significant peaks by green dashed lines. (B) Annotation of H3Kac upregulated peaks in *gyrb3-2*. “gene+TE” corresponds to peaks overlapping with both genes and transposons, “gene” to peaks overlapping with genes, and “TE” to peaks overlapping with transposons. All other peaks were classified as “Intergenic.” (C) DNA methylation levels in H3Kac peaks significantly upregulated (log2FC ≥ 1, p-value < 0.05) in *gyrb3-2* versus Col-0. (D) UpSetR plot showing all possible combinations of overlaps between H3Kac peaks differently accumulating in *gyrb3-2*, *ibm2-4*, *ibm2-4 gyrb3-2* versus Col-0 or *ibm2-4*. The numbers represent the count of overlapping peaks for the category shown in the bottom left panel. (E) Overlaps between H3Kac peaks significantly upregulated (log2FC ≥ 1, p-value < 0.05) in *gyrb3-2, ibm2-4 and ibm2-4* gyrb3-2 compared to the Col-0 control. (F) H3Kac levels in the Col-0 control, *gyrb3-2*, *ibm2-4* and *ibm2-4 gyrb3-2* for regions accumulating H3Kac in *gyrb3-2* compared to Col-0 (n=74). ChIP signals were quantified using normalized *BigWig* files scaled to 1 million mapped reads. The detected signals in the inputs (*inp.*) served as controls. (G) *JBrowse2* browser views of normalized H3Kac ChIP signals (two repeats are shown per genotype) in Col-0, *gyrb3-2*, *ibm2-4 gyrb3-2* and *ibm2-4* for four regions containing H3Kac peaks significantly upregulated (log2FC ≥ 1, p-value < 0.05) in *gyrb3-2* and taken as examples. The genomic positions relative to the chromosome start are indicated. *BigWig* files used for visualization were scaled to 1 million mapped reads.

To examine changes occurring for histone acetylation profiles when combining the *gyrb3* and *ibm2* mutations, H3Kac ChIP-seq analyses were performed for *ibm2-4* and *ibm2-4 gyrb3-2* mutants. Using the methods applied to *gyrb3-2*, peaks that were significantly up- or downregulated (log2FC or ≥ 1 or ≤ -1; p-value < 0.05) were identified in all mutant backgrounds (Table 2 and Fig. 5D). Out of the 74 regions enriched for H3Kac in *gyrb3-2* compared to Col-0, 77% (n=57) remained in the same state in *ibm2-4 gyrb3-2* (Fig. 5D and 5E). This was confirmed by plotting and comparing the normalized ChIP H3Kac signal of these regions for all genotypes (Fig. 5F and 5G). Thus, the accumulation of H3Kac observed in *gyrb3* at TEs also occurs in the double *ibm2 gyrb3* mutant. A total of 136 regions were enriched for H3Kac in the *ibm2* mutant, while 44 regions exhibited reduced H3Kac levels. Most of these regions were located within genes (44% of the H3Kac-enriched regions and 93% of the depleted regions), consistent with the drastic epigenetic changes observed in genes in both *ibm1* and *ibm2*. We identified regions (n=22) with altered H3Kac levels in the *ibm2* mutant which were restored in the *ibm2 gyrb3* mutant (Fig. 5D). This implies that acetylation changes occur between *ibm2 gyrb3*, *gyrb3* and the Col-0 background.

**Table 2.** Number of H3Kac peaks up- or downregulated (log2FC ≥ 1 or ≤ -1; p-value < 0.05) in the genotypes indicated.

| Genotypes compared | H3Kac peaks |  |
| --- | --- | --- |
|  | up | down |
| *gyrb3-2* versus Col-0 | 74 | 1 |
| *ibm2-4 gyrb3-2* versus Col-0 | 94 | 9 |
| *ibm2-4* versus Col-0 | 136 | 44 |
| *ibm2-4 gyrb3-2* versus *ibm2-4* | 44 | 32 |

### GyrB3 is a nuclear protein interacting with HDA6

Given its role in histone acetylation and the lack of catalytic domain, we investigated whether GyrB3 directly interacts with histone deacetylases. First, we employed the AlphaFold-Multimer (AFM) software (52) to predict the structure of GyrB3 in association with all HDA proteins of Arabidopsis. Using the interface-predicted template modelling (ipTM) score derived from AFM as a measure of the structural accuracy of a protein complex, GyrB3 was predicted to interact with all RPD3-like HDACs of Class I, namely HDA6, HDA7, HDA9, and HDA19, but not with other HDAC family members (Fig. 6A). Restricting the analysis to the three predicted domains of GyrB3 in association with HDA6, HDA9, or HDA19 suggested that the ELM2 domain is a strong candidate for interaction between GyrB3 and HDA proteins, contrarily to the topoisomerase or the SANT domains (Fig. 6B). This was further confirmed by visualizing the structure of the protein complex formed between HDA6 and GyrB3 (Fig. 6C) and by visualising the Predicted Aligned Error (PAE) data provided in AFM outputs (Supplementary Fig. S8). Results from a yeast two-hybrid (Y2H) assay further demonstrated that GyrB3 physically interacts *in vivo* with HDA6, HDA9 and HDA19, and weakly with HDA7 (Fig. 6D and Supplementary Fig. S9). Although GyrB3 was found to interact with itself (Fig. 6D), this interaction could not be confirmed using AFM. Based on both *in silico* and yeast analyses, we concluded that GyrB3 likely interacts with the Class I HDA proteins.

**Fig. 6.**
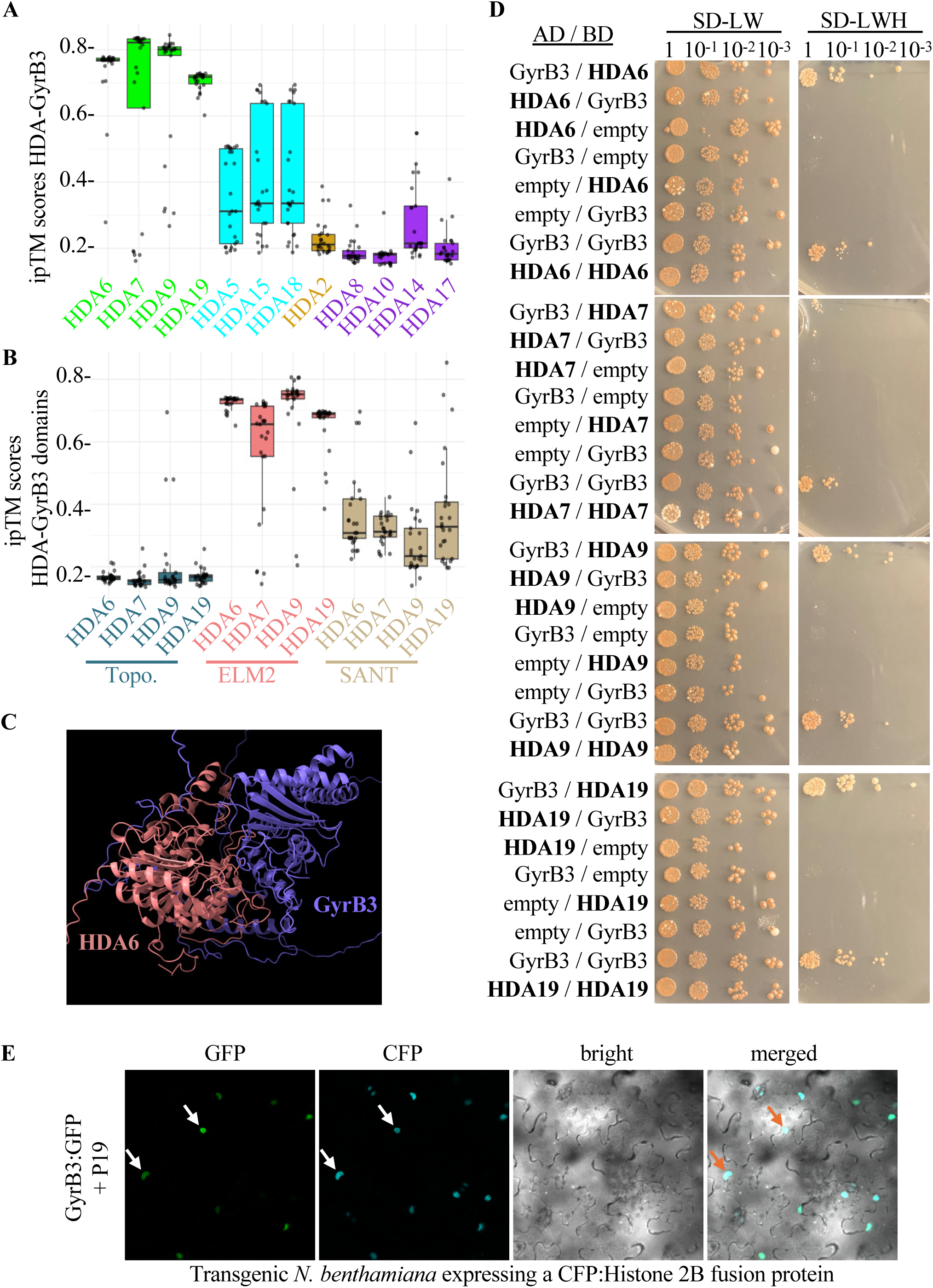
GyrB3 is a nuclar protein interacting with histone deacetylases (HDAs) (A) Plot of the interface-predicted template modelling (ipTM) scores derived from the AlphaFold-Multimer (AFM) software to measure the structural accuracy of the protein complexes predicted between GyrB3 and the HDA proteins of Arabidopsis. The plots were built using the ipTM scores derived from five different models. Arabidopsis RPD3-like HDACs are divided into four classes: Class I in green (HDA6, HDA7, HDA9, HDA19), Class II in cyan (HDA5, HDA15, HDA18), Class III in brown (HDA2), and unclassified in purple (HDA8, HDA10, HDA14, HDA17). (B) Plot of the interface-predicted template modelling (ipTM) scores derived from the AlphaFold-Multimer (AFM) software to measure the structural accuracy of the protein complexes predicted between the three GyrB3 predicted protein domains and HDA6, HDA7, HDA9 or HDA19. The plots were built using the ipTM scores derived from five different models. The predicted domains of GyrB3 are described in Fig. 1A. (C) Predicted HDA6/GyrB3 protein complex visualized with *Chimera* v1.9. (D) Yeast-two-hybrid assays testing interactions between HDA6, HDA7, HDA9, HDA19 and GyrB3. Full length proteins were fused with Gal4 DNA binding domain (BD) and Gal4 activation domain (AD), respectively, and co-expressed in yeast cells. For each combination, serial dilutions of yeast cells were spotted on non-selective medium (-LW), or selective media (-LWH). An additional repeat of this experiment is shown in Supplementary Fig. S9. (E) Nuclear localization of GyrB3:GFP fusion protein in tobacco cells. Subcellular localization patterns were examined in the epidermal cells of *N. benthamiana* leaves from transgenic plants stably expressing a nuclear CFP:Histone 2B fusion protein as a nuclear marker (45), following agroinfiltration with *a p35S::GyrB3:GFP* (GyrB3:GFP) construct and a *p35S::P19* (P19) construct, which enhances transient expression (47). Fluorescence was analyzed 48 hours after infiltration. Supplementary Fig. S10 provides additional data for the GFP:GyrB3 fusion and the P19 control. Arrows indicate representative nuclei.

Since GyrB3 may be involved in chromatin regulation, we tested whether the protein localizes to the nucleus. GFP:GYRB3 and GYRB3:GFP fusion constructs were generated and transiently expressed in tobacco leaves to determine the subcellular localization of GyrB3. Confocal microscopy revealed that the GFP signal was specifically localized to the nucleus (Fig. 6E and supplementary Fig. S10), indicating that GyrB3 is a nuclear-targeted protein, consistent with the proposed role of GyrB3 in regulating chromatin-associated processes with HDA proteins.

### GyrB3 and HDA6 share a common set of genomic targets

To further elucidate the relationships between GyrB3 and HDA6 or HDA19 proteins, we retrieved raw data from three independent H3Kac ChIP-seq experiments involving *hda* mutants (41–43) that were collected under conditions comparable to ours (bulks of seedlings grown *in vitro* for 15 days). The H3Kac normalized ChIP signals were monitored in regions accumulating H3Kac in *gyrb3-2*. For the first dataset (41), the H3Kac signal was increased in the two *hda6* biological replicates, but not in *hda19*, neither in the five wild-type control repeats nor when the signal was examined in regions randomly chosen (Fig. 7A). For the second dataset (42), we observed a comparable increase for the two *hda6* H3Kac-ChIP biological repeats, but not for the inputs, or any of the controls (Fig. 7B). A third dataset (43), gave identical results (Supplementary Fig. S11). Therefore, the regions that accumulate H3Kac in *gyrb3-2* also accumulate H3Kac in *hda6*. To characterize how these regions are epigenetically controlled, the analysis was extended to DNA methylation, including additional mutants, such as *met1-9* and *ros1-1*, using publicly available datasets (34, 35). HDA6 and MET1 physically interacts and cooperate to promote DNA methylation; thus many regions deregulated in *hda6* mutants show impaired CG and non-CG methylation (3, 4). ROS1 is a DNA demethylase, and mutations in the *ROS1* gene lead to genome-wide DNA hypermethylation, except for certain pericentromeric regions, likely due to compensatory activity by other DNA demethylases (35). Differences of methylation occurring in *ros1-1* are reversed by mutating *HDA6* (35, 53, 54). Plotting methylation levels revealed that regions hyperacetylated in both *gyrb3-2* and *hda6* are also hypomethylated in the CHG context in *met1-9* and display reduced CG methylation in *hda6* and *met1* mutants (Fig. 7C). This aligns with prior findings that CG and non-CG methylation are interconnected at HDA6-targeted regions (3, 4). However, in a *gyrb3* background this coupling is lost since *gyrb3* retain CG methylation in regions targeted by both enzymes (Fig. 7C). No changes were observed in *ros1* (Fig. 7C).

**Fig. 7.**
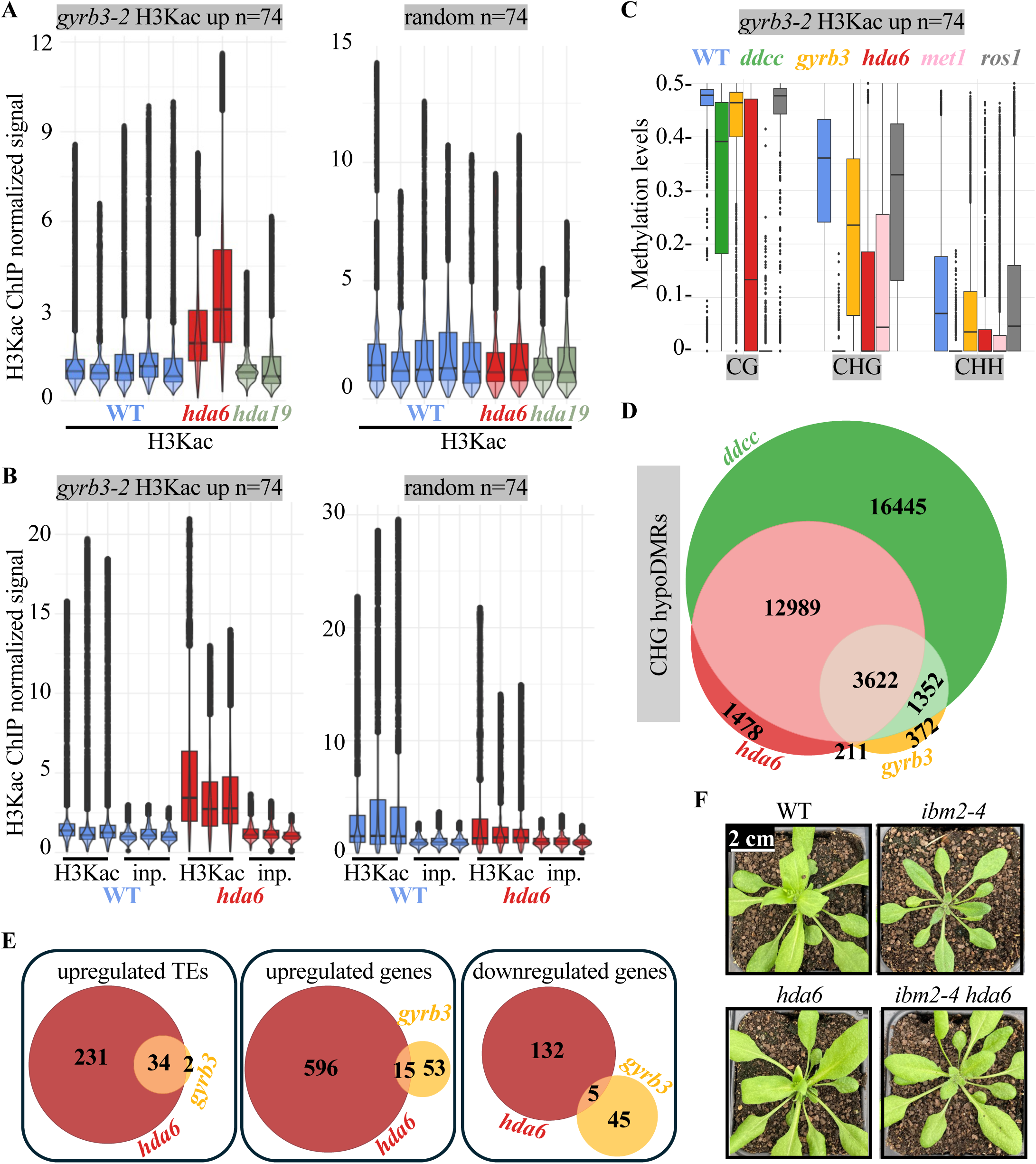
HDA6 and GyrB3 target common genomic regions for H3Kac and methylation. (A) H3Kac levels in five wild-types (WT), two *hda6* (*axe1-5*) and two *hda19* (SALK_139445) biological repeats in regions accumulating H3Kac in *gyrb3-2* (n=74). ChIP signals were quantified using normalized *BigWig* files scaled to 1 million mapped reads. The data (41) were retrieved from the GEO repository (accession number GSE166090). As a control, ChIP signals were also measured in randomly selected regions of similar length. (B) H3Kac levels in three wild-type (WT) and three *hda6* (*axe1-5*) biological repeats, plus their corresponding input DNAs (*inp.*), in regions accumulating H3Kac in *gyrb3-2* (n=74). ChIP signals were quantified using normalized *BigWig* files scaled to 1 million mapped reads. The data (42) were retrieved from the GEO repository (accession number GSE167288). As a control, ChIP signals were also measured in randomly selected regions of similar length. (C) DNA methylation levels in *gyrb3-2* H3Kac peaks significantly upregulated (log2FC ≥ 1, p-value < 0.05) in epigenetic mutants such as the *drm1 drm2 cmt3 cmt2* quadruple mutant (*ddcc*), *hda6* (*axe1-5*), *met1-9*, and *ros1-1*. (D) Venn diagram showing overlaps between CHG hypoDMRs identified in *gyrb3-2* with those found in *hda6* (*axe1-5*) and the *drm1 drm2 cmt3 cmt2* quadruple mutant (*ddcc*) in which all non-CG methylation is abolished. (E) Overlaps between genes or TEs differentially expressed in *hda6* (*axe1-5*) and *gyrb3-2* mutants. The *hda6* RNA-seq data (42) were retrieved from the GEO repository (accession number GSE167288). (F) Pictures of the wild-type (WT), *ibm2-4*, *hda6* (*axe1-5*) and the double *ibm2-4 hda6* F3 plants taken 30 days after sowing. Lines were obtained by self-pollinating F2 plants from a cross between *ibm2-4* and *hda6*. All plants were individually genotyped for the mutations in both F2s and F3s. Supplementary Fig. S13 shows the same plants after an additional week of growth, pictured in the greenhouse.

Because *gyrb3* contains far more DMRs than the number of H3Kac peaks identified, we investigated whether the genomic regions hypomethylated in *gyrb3* overlapped with those in *hda6*, using publicly available BS-seq datasets (8) and focusing on CHG methylation, where the most pronounced differences were observed (Fig. 2 and 3). Out of the 5557 CHG hypoDMRs identified in *gyrb3* (Fig. 3C), 69% overlapped with CHG hypoDMRs identified in *hda6* using our analysis pipeline (Fig. 7D), suggesting that GyrB3 and HDA6 likely function in the same pathway for regulating CHG methylation in these regions. Almost all CHG hypoDMRs identified in both *hda6* and *gyrb3* were present in a *drm1 drm2 cmt3 cmt2* (*ddcc*) quadruple mutant (Fig. 7D) in which all non-CG methylation is abolished (34). All TEs exhibiting hyperacetylation in *gyrb3* are covered by these CHG hypoDMRs (Supplementary Table S4). Comparable trends were also observed for CHH hypoDMRs (Supplementary Fig. S12A). Regions that are CHG hypomethylated in both *gyrb3* and *hda6* retained CG methylation similarly to the wild-type across all mutant backgrounds, to the exception of *met1-9* which serves as a control (Supplementary Fig. S12B-C). Notably, the regions within *gyrb3* that accumulate H3Kac (Fig. 7C) constitute only a small fraction of the *gyrb3/hda6* CHG hypoDMRs, which explains the overall differences observed in CG methylation trends for *hda6* (Supplementary Fig7C and Fig. S12C). In contrast, the CG hypoDMRs identified in *hda6* (n=5997) exhibit major changes in levels of non-CG methylation, as previously reported (3, 4), including in the *ros1-1* background (Supplementary Fig. S12D). This suggests that GyrB3 may regulate non-CG methylation in conjunction with HDA6, independently of pathways that rely on ROS1 and MET1.

To identify whether the expression of genes and TEs could be similarly affected in both *hda6* and *gyrb3*, we analysed previously published RNA-seq data (42) generated from the same tissue types used in our study (i.e. 12-day-old seedlings grown under long day conditions). We performed RNA-seq analyses using the same pipelines as those applied to our own RNA-seq data. Almost all TEs upregulated (log2FC ≥ 1; p-value < 0.05; based on *DESeq2*) in *gyrb3-2* were also upregulated in *hda6* (Fig. 7E). In contrast, the overlaps between genes that were differentially expressed was much more limited (Fig. 7E). Therefore, the common targets of GyrB3 and HDA6 are mainly TEs, particularly those belonging to the LTR-retrotransposon family (Supplementary Table S4). Only 15% of all TEs upregulated in *hda6* are controlled by GyrB3, suggesting that HDA6 also regulates TEs through pathways independent of GyrB3. Alternatively, other GyrB3-related enzymes may cooperate with HDA6 in *gyrb3* mutants to regulate TEs, suggesting functional redundancy.

Finally, since GyrB3 suppressed the developmental phenotype of both *ibm1* and *ibm2* mutants (Fig. 1B-C, supplementary Fig. S1 and 2), and given the functional overlap between HDA6 and GyrB3, we tested whether an *hda6* mutation, like *gyrb3*, suppresses the phenotype of *ibm2*. The *hda6* (*axe1-5* allele) mutant (28) was crossed with *ibm2-4*. In a segregating F3 progeny, only plants that carry the *ibm2-4* homozygous mutation fixed in the F2s presented developmental anomalies. All other genotypes, including the double *ibm2-4 hda6*, mutant developed normally (Fig. 7F and Supplementary Fig. S13). Therefore, similar to *GYRB3*, a mutation in *HDA6* suppresses the *Ibm2* phenotype, confirming the functional overlap between HDA6 and GyrB3.

## DISCUSSION

Histone deacetylation regulates TEs through several interconnected mechanisms (2–9). Our study reveals for the first time the role played by GyrB3, a nuclear protein that interacts with histone deacetylases such as HDA6 and helps maintain epigenetic balance by silencing TEs.

### Suppressor mutagenesis of *ibm2* highlights new components of epigenetic regulations

A genetic screen to identify suppressors of *ibm2-4* revealed that a mutation in *GYRB3* rescues the developmental phenotypes of *ibm1* and *ibm2* (Fig. 1B-D and Supplementary Fig. S1 and S2). Other mutations that suppress the phenotypes of *ibm1* have been identified in previous studies. Mutating *FPA* in *ibm2* restores *IBM1* transcription, which is otherwise impaired. *fpa* is a suppressor of *ibm2* rescuing the expression of the long functional *IBM1* transcript, as well as other transcripts targeted by IBM2 (22). Nevertheless, most of the suppressor mutations identified from the screening of *ibm2*-mutagenised plants fall into a category that predominantly compensates for the genome-wide loss of *IBM1* function, rather than restoring the expression of *IBM1*. For instance, mutations in *KYP* and *CMT3* disrupt the reinforcing loop responsible for DNA/H3K9 methylation invasion occurring in genes of *ibm1* mutants. Most suppressors (56%) found in our screen are *kyp* or *cmt3* mutants (Supplementary Table S3), indicating that removing the proteins responsible for inappropriate spreading of heterochromatin into genes is the most effective strategy for rescuing *ibm* mutants. LDL2 reduces H3K4 methylation at genes enriched for H3K9me2 in *ibm1*, resulting in reduced gene expression. This effect can be reversed in an *ldl2* mutant (48) which was also identified in our genetic suppressor screen (Supplementary Table S3). In the present study, we show that mutations in *GYRB3* partially reduce the CHG and CHH hypermethylation occurring in genes in the absence of *IBM1/2* (Fig. 2). Changes in H3Kac levels were observed between *ibm2* and *ibm2 gyrb3* lines (Fig. 5F-G). Together, this suggests that mutating *GYRB3* may act at multiple loci to compensate for IBM1/2 deficiency, although the underlying mechanism remains to be clarified.

### GyrB3 illustrates plant adaptation for TE epigenetic control

In plants, prokaryotic-origin DNA gyrase coexists with eukaryotic DNA topoisomerase II to relax supercoiled DNA. The Arabidopsis genome encodes one *GYRA* and three *GYRB* genes, with GyrA, GyrB1, and GyrB2 targeted to organelles (chloroplasts and mitochondria) while GyrB3 is a nuclear protein (Fig. 6E and supplementary Fig. S10). GyrB3 is smaller than other GyrBs; sequence analysis reveals that GyrB3 lacks key conserved motifs found in GyrB proteins and a significant portion of the N-terminal domain essential for ATP hydrolysis and for controlling DNA topology (27). Phylogenetic analysis indicates that GyrB3 aligns with eukaryotic type II topoisomerases, suggesting a different evolutionary origin from GyrA, GyrB1, and GyrB2, which are closely related to cyanobacterial gyrases (55). Consequently, the function of GyrB3 as a gyrase subunit remained ambiguous (27). Our findings reveal that GyrB3 contributes to the regulation of epigenetic modifications. In a *gyrb3* mutant, a subset of TEs become transcriptionally active (Table 1) exhibiting reduced DNA methylation (Fig. 3F) associated with an increase in histone H3 acetylation (Fig. 5C). Therefore, GyrB3 likely plays a role in suppressing TE activity by promoting histone deacetylation mediated by HDA enzymes. GyrB3 have diverged from the typical gyrase subunit function, adopting roles related to DNA binding and gene regulation. GyrB3 might be another example of how plants utilize the toolbox of protein domains inherited from cyanobacteria, thereby enhancing their own eukaryotic cellular functions (56).

### GyrB3 functions as a new potential component of histone deacetylation complexes

*In vivo* interaction assays suggest that GyrB3 physically interacts with HDA6, HDA9, and HDA19 and weakly with HDA7, in agreement with recent results showing that HDA7 is inactive (57). Structure predictions suggest that the interaction with HDAs occurs through the ELM2 domain of GyrB3 (Fig. 6B). Similarly to *gyrb3*, a mutation in *hda6* also suppresses the *ibm2* mutant phenotype, suggesting that GyrB3 and HDA6 may act in overlapping or convergent pathways (Fig 7E and supplementary Fig. S13).

GyrB3 exhibits homology to the ELM2 and SANT domains, which are frequently associated with complexes involving histone deacetylases in both animals and plants. The Nucleosome Remodelling and Deacetylase (NuRD) is a major mammalian chromatin remodelling complex, that includes proteins containing ELM2-SANT domains (58). In Arabidopsis, ESANT1 and ESANT2 are ELM2-SANT containing domain proteins interacting with HDA6 and HDA19 (41). SANT proteins (SANT1-4) form distinct complexes with HDA6 to regulate gene expression by modulating histone deacetylation and by coordinating H3Kac and lysine 2-hydroxyisobutyrylation (Khib) (41, 42, 51). Mutations in *SANT* upregulate heat-inducible genes and enhance thermotolerance (51). They also cause delayed flowering, associated with increased expression of floral repressors such as FLC, MAF4, and MAF5 (42), a phenotype not observed in *gyrb3* mutants, although further studies are required to confirm this observation. Together with HDA6, SANT proteins are crucial for histone deacetylation, but unlike GyrB3, they do not influence the silencing of TEs (41, 42, 59). Thus, GyrB3 might be part of an HDA6 complex different from the ones already described in Arabidopsis. This hypothesis is supported by the fact that GyrB3 has not been identified as an interactor of HDA proteins in any of the studies aimed at identifying HDA protein partners (6, 6, 41, 42, 60). Given that both *gyrb3* and *hda6* exhibit similar patterns of H3Kac enrichment at specific genomic loci (Fig. 7A-B and supplementary Fig. S11), that most *gyrb3* CHG hypoDMRs are shared with *hda6* (Fig. 7D), and that most of the TEs targeted by GyrB3 are also targets of HDA6 (Fig. 7E), it is plausible that GyrB3 may facilitate the recruitment or retention of HDA6 at these sites. Alternatively, GyrB3 could influence the enzymatic activity or conformational stability of HDA6. Such a role would position GyrB3 as an important regulator of histone deacetylation and chromatin remodeling.

### HDA6-mediated epigenetic pathways converge to repress TEs

Previous studies have shown that HDA6 facilitates chromatin condensation by interacting with key enzymes involved in various epigenetic pathways. The physical interaction between HDA6 and MET1 is critical for the silencing of a specific subset of TEs (3, 4). In *hda6* mutants, CG methylation is reduced (3, 4, 7) (Supplementary Fig. S12D), indicating that HDA6 facilitates MET1-mediated silencing. In contrast to *hda6*, *gyrb3* mutants largely retain CG methylation, including in regions targeted by both HDA6 and GyrB3 (Fig. 7C and supplementary Fig. S12C), but exhibit a specific loss of CHG and CHH methylation (Fig. 3 and supplementary Fig. S3). Together, these findings indicate that GyrB3 likely links HDA6 to non-CG methylation pathways, consistent with the observation that *gyrb3* suppresses *ibm1* and *ibm2*, which hyperaccumulate CHG methylation.

HDA6 also interacts with the H3K9 histone methyltransferases (6). HDA6 and SUVH4/5/6 collaborate by removing acetyl groups and adding methyl groups to histone H3, resulting in chromatin condensation and transcriptional silencing to regulate TEs (6). In addition, HDA6 plays a role in repressing active histone methylation marks such as H3K4me3 and H3K4me2 at TEs by physically interacting with histone demethylases like FLOWERING LOCUS D (FLD) (10). Another HDA6 epigenetic interactor is MULTICOPY SUPPRESSOR OF IRA1 (MSI1), a conserved component of the Polycomb Repressive Complex 2 (PRC2), which facilitates gene silencing through H3K27me3 (61). Using AlphaFold, we investigated whether GyrB3 could be predicted to interact with MSI1, MBD2, MBD4, CMT3, MET1, SUVH4, SUVH6, but found no supporting evidence. Additional research is required to decipher all protein interactors of GyrB3 beyond HDA proteins and to better understand the specific function of the GyrB3 complex, particularly its relationship with other key regulators of DNA methylation.

In summary, our findings demonstrate that GyrB3, despite not being a functional subunit of DNA gyrase, is involved in epigenetic regulation in Arabidopsis. Specifically, GyrB3 is a nuclear protein that interacts with histone deacetylases such as HDA6, contributing to the silencing of TEs, and likely playing a key role in maintaining genome stability by preventing transposition. Future investigations should focus on clarifying the precise role of GyrB3 in pollen cells and its contribution to TEs silencing during male gametogenesis when GyrB3 is highly expressed (Supplementary Fig. S14), and when transposition must be strictly regulated.

## ACKNOWLEDGMENTS

We thank Vincent Coustham for helpful discussions and for critical reading of the manuscript. We thank T. Kakutani and H. Saze for providing the *ibm2-1* and the *ibm1-1* mutants. We are very grateful to A. Chambon and A. Vayssières for their assistance with the confocal microscope.

## REFERENCES

1. Bannister, A.J. and Kouzarides, T. (2011) Regulation of chromatin by histone modifications. Cell Res., 21, 381–395.

2. Probst, A.V., Fagard, M., Proux, F., Mourrain, P., Boutet, S., Earley, K., Lawrence, R.J., Pikaard, C.S., Murfett, J., Furner, I., et al. (2004) Arabidopsis histone deacetylase HDA6 is required for maintenance of transcriptional gene silencing and determines nuclear organization of rDNA repeats. Plant Cell, 16, 1021–1034.

3. To, T.K., Kim, J.-M., Matsui, A., Kurihara, Y., Morosawa, T., Ishida, J., Tanaka, M., Endo, T., Kakutani, T., Toyoda, T., et al. (2011) Arabidopsis HDA6 regulates locus-directed heterochromatin silencing in cooperation with MET1. PLoS Genet., 7, e1002055.

4. Liu, X., Yu, C.-W., Duan, J., Luo, M., Wang, K., Tian, G., Cui, Y. and Wu, K. (2012) HDA6 directly interacts with DNA methyltransferase MET1 and maintains transposable element silencing in Arabidopsis. Plant Physiol., 158, 119–129.

5. Blevins, T., Pontvianne, F., Cocklin, R., Podicheti, R., Chandrasekhara, C., Yerneni, S., Braun, C., Lee, B., Rusch, D., Mockaitis, K., et al. (2014) A two-step process for epigenetic inheritance in Arabidopsis. Mol. Cell, 54, 30–42.

6. Yu, C.-W., Tai, R., Wang, S.-C., Yang, P., Luo, M., Yang, S., Cheng, K., Wang, W.-C., Cheng, Y.-S. and Wu, K. (2017) HISTONE DEACETYLASE6 Acts in Concert with Histone Methyltransferases SUVH4, SUVH5, and SUVH6 to Regulate Transposon Silencing. Plant Cell, 29, 1970–1983.

7. Yang, J., Yuan, L., Yen, M.-R., Zheng, F., Ji, R., Peng, T., Gu, D., Yang, S., Cui, Y., Chen, P.-Y., et al. (2020) SWI3B and HDA6 interact and are required for transposon silencing in Arabidopsis. *Plant J*. Cell Mol. Biol., 102, 809–822.

8. Hsieh, J.-W.A., Yen, M.-R., Hung, F.-Y., Wu, K. and Chen, P.-Y. (2024) Epigenetic factors direct synergistic and antagonistic regulation of transposable elements in Arabidopsis. Plant Physiol., 196, 1939–1952.

9. Li, W., Zhang, X., Zhang, Q., Li, Q., Li, Y., Lv, Y., Liu, Y., Cao, Y., Wang, H., Chen, X., et al. (2024) PICKLE and HISTONE DEACETYLASE6 coordinately regulate genes and transposable elements in Arabidopsis. Plant Physiol., 196, 1080–1094.

10. Yu, C.-W., Liu, X., Luo, M., Chen, C., Lin, X., Tian, G., Lu, Q., Cui, Y. and Wu, K. (2011) HISTONE DEACETYLASE6 interacts with FLOWERING LOCUS D and regulates flowering in Arabidopsis. Plant Physiol., 156, 173–184.

11. Saze, H., Shiraishi, A., Miura, A. and Kakutani, T. (2008) Control of genic DNA methylation by a jmjC domain-containing protein in Arabidopsis thaliana. Science, 319, 462–465.

12. Du, J., Zhong, X., Bernatavichute, Y.V., Stroud, H., Feng, S., Caro, E., Vashisht, A.A., Terragni, J., Chin, H.G., Tu, A., et al. (2012) Dual binding of chromomethylase domains to H3K9me2-containing nucleosomes directs DNA methylation in plants. Cell, 151, 167–180.

13. Du, J., Johnson, L.M., Groth, M., Feng, S., Hale, C.J., Li, S., Vashisht, A.A., Wohlschlegel, J.A., Patel, D.J. and Jacobsen, S.E. (2014) Mechanism of DNA methylation-directed histone methylation by KRYPTONITE. Mol. Cell, 55, 495–504.

14. Johnson, L.M., Bostick, M., Zhang, X., Kraft, E., Henderson, I., Callis, J. and Jacobsen, S.E. (2007) The SRA methyl-cytosine-binding domain links DNA and histone methylation. Curr. Biol. CB, 17, 379–384.

15. Miura, A., Nakamura, M., Inagaki, S., Kobayashi, A., Saze, H. and Kakutani, T. (2009) An Arabidopsis jmjC domain protein protects transcribed genes from DNA methylation at CHG sites. EMBO J., 28, 1078–1086.

16. Rigal, M., Kevei, Z., Pélissier, T. and Mathieu, O. (2012) DNA methylation in an intron of the IBM1 histone demethylase gene stabilizes chromatin modification patterns. EMBO J., 31, 2981–2993.

17. Saze, H., Kitayama, J., Takashima, K., Miura, S., Harukawa, Y., Ito, T. and Kakutani, T. (2013) Mechanism for full-length RNA processing of Arabidopsis genes containing intragenic heterochromatin. Nat. Commun., 4, 2301.

18. Wang, X., Duan, C.-G., Tang, K., Wang, B., Zhang, H., Lei, M., Lu, K., Mangrauthia, S.K., Wang, P., Zhu, G., et al. (2013) RNA-binding protein regulates plant DNA methylation by controlling mRNA processing at the intronic heterochromatin-containing gene IBM1. Proc. Natl. Acad. Sci. U. S. A., 110, 15467–15472.

19. Coustham, V., Vlad, D., Deremetz, A., Gy, I., Cubillos, F.A., Kerdaffrec, E., Loudet, O. and Bouché,N. (2014) SHOOT GROWTH1 maintains Arabidopsis epigenomes by regulating IBM1. PloS One, 9, e84687.

20. Duan, C.-G., Wang, X., Zhang, L., Xiong, X., Zhang, Z., Tang, K., Pan, L., Hsu, C.-C., Xu, H., Tao, W.A., et al. (2017) A protein complex regulates RNA processing of intronic heterochromatin-containing genes in Arabidopsis. Proc. Natl. Acad. Sci. U. S. A., 114, E7377–E7384.

21. Osabe, K., Harukawa, Y., Miura, S. and Saze, H. (2017) Epigenetic Regulation of Intronic Transgenes in Arabidopsis. Sci. Rep., 7, 45166.

22. Deremetz, A., Le Roux, C., Idir, Y., Brousse, C., Agorio, A., Gy, I., Parker, J.E. and Bouché, N. (2019) Antagonistic Actions of FPA and IBM2 Regulate Transcript Processing from Genes Containing Heterochromatin. Plant Physiol., 180, 392–403.

23. Zhang, Y.-Z., Lin, J., Ren, Z., Chen, C.-X., Miki, D., Xie, S.-S., Zhang, J., Chang, Y.-N., Jiang, J., Yan, J., et al. (2021) Genome-wide distribution and functions of the AAE complex in epigenetic regulation in Arabidopsis. J. Integr. Plant Biol., 63, 707–722.

24. Raingeval, M., Leduque, B., Baduel, P., Edera, A., Roux, F., Colot, V. and Quadrana, L. (2024) Retrotransposon-driven environmental regulation of FLC leads to adaptive response to herbicide. Nat. Plants, 10, 1672–1681.

25. Wang, J. and Eulgem, T. (2023) The Arabidopsis RRM domain proteins EDM3 and IBM2 coordinate the floral transition and basal immune responses. *Plant J*. Cell Mol. Biol., 116, 128–143.

26. Wang, J. and Eulgem, T. (2024) Growth deficiency and enhanced basal immunity in Arabidopsis thaliana mutants of EDM2, EDM3 and IBM2 are genetically interlinked. PloS One, 19, e0291705.

27. Evans-Roberts, K.M., Breuer, C., Wall, M.K., Sugimoto-Shirasu, K. and Maxwell, A. (2010) Arabidopsis thaliana GYRB3 Does Not Encode a DNA Gyrase Subunit. PLOS ONE, 5, e9899.

28. Murfett, J., Wang, X.J., Hagen, G. and Guilfoyle, T.J. (2001) Identification of Arabidopsis histone deacetylase HDA6 mutants that affect transgene expression. Plant Cell, 13, 1047–1061.

29. Easlon, H.M. and Bloom, A.J. (2014) Easy Leaf Area: Automated digital image analysis for rapid and accurate measurement of leaf area. Appl. Plant Sci., 2, apps.1400033.

30. Catoni, M., Tsang, J.M., Greco, A.P. and Zabet, N.R. (2018) DMRcaller: a versatile R/Bioconductor package for detection and visualization of differentially methylated regions in CpG and non-CpG contexts. Nucleic Acids Res., 46, e114.

31. Ramírez, F., Ryan, D.P., Grüning, B., Bhardwaj, V., Kilpert, F., Richter, A.S., Heyne, S., Dündar, F. and Manke, T. (2016) deepTools2: a next generation web server for deep-sequencing data analysis. Nucleic Acids Res., 44, W160–165.

32. Gu, Z., Gu, L., Eils, R., Schlesner, M. and Brors, B. (2014) circlize Implements and enhances circular visualization in R. Bioinforma. Oxf. Engl., 30, 2811–2812.

33. Jouffroy, O., Saha, S., Mueller, L., Quesneville, H. and Maumus, F. (2016) Comprehensive repeatome annotation reveals strong potential impact of repetitive elements on tomato ripening. BMC Genomics, 17, 624.

34. Zhao, L., Zhou, Q., He, L., Deng, L., Lozano-Duran, R., Li, G. and Zhu, J.-K. (2022) DNA methylation underpins the epigenomic landscape regulating genome transcription in Arabidopsis. Genome Biol., 23, 197.

35. Wang, Q., Bao, X., Chen, S., Zhong, H., Liu, Y., Zhang, L., Xia, Y., Kragler, F., Luo, M., Li, X.D., et al. (2021) AtHDA6 functions as an H3K18ac eraser to maintain pericentromeric CHG methylation in Arabidopsis thaliana. Nucleic Acids Res., 49, 9755–9767.

36. Love, M.I., Huber, W. and Anders, S. (2014) Moderated estimation of fold change and dispersion for RNA-seq data with DESeq2. Genome Biol., 15, 550.

37. Rau, A. and Maugis-Rabusseau, C. (2018) Transformation and model choice for RNA-seq co-expression analysis. Brief. Bioinform., 19, 425–436.

38. Yu, G., Wang, L.-G., Han, Y. and He, Q.-Y. (2012) clusterProfiler: an R package for comparing biological themes among gene clusters. Omics J. Integr. Biol., 16, 284– 287.

39. Zhang, Y., Liu, T., Meyer, C.A., Eeckhoute, J., Johnson, D.S., Bernstein, B.E., Nusbaum, C., Myers, R.M., Brown, M., Li, W., et al. (2008) Model-based analysis of ChIP-Seq (MACS). Genome Biol., 9, R137.

40. Tu, S., Li, M., Chen, H., Tan, F., Xu, J., Waxman, D.J., Zhang, Y. and Shao, Z. (2021) MAnorm2 for quantitatively comparing groups of ChIP-seq samples. Genome Res., 31, 131–145.

41. Feng, C., Cai, X.-W., Su, Y.-N., Li, L., Chen, S. and He, X.-J. (2021) Arabidopsis RPD3-like histone deacetylases form multiple complexes involved in stress response. J. Genet. Genomics Yi Chuan Xue Bao, 48, 369–383.

42. Zhou, X., He, J., Velanis, C.N., Zhu, Y., He, Y., Tang, K., Zhu, M., Graser, L., de Leau, E., Wang, X., et al. (2021) A domesticated Harbinger transposase forms a complex with HDA6 and promotes histone H3 deacetylation at genes but not TEs in Arabidopsis. J. Integr. Plant Biol., 63, 1462–1474.

43. Hung, F.-Y., Chen, C., Yen, M.-R., Hsieh, J.-W.A., Li, C., Shih, Y.-H., Chen, F.-F., Chen, P.-Y., Cui, Y. and Wu, K. (2020) The expression of long non-coding RNAs is associated with H3Ac and H3K4me2 changes regulated by the HDA6-LDL1/2 histone modification complex in Arabidopsis. NAR Genomics Bioinforma., 2, lqaa066.

44. Mirdita, M., Schütze, K., Moriwaki, Y., Heo, L., Ovchinnikov, S. and Steinegger, M. (2022) ColabFold: Making Protein folding accessible to all. Nat. Methods, 10.1038/s41592-022-01488-1.

45. Martin, K., Kopperud, K., Chakrabarty, R., Banerjee, R., Brooks, R. and Goodin, M.M. (2009) Transient expression in Nicotiana benthamiana fluorescent marker lines provides enhanced definition of protein localization, movement and interactions in planta. *Plant J*. Cell Mol. Biol., 59, 150–162.

46. Azimzadeh, J., Nacry, P., Christodoulidou, A., Drevensek, S., Camilleri, C., Amiour, N., Parcy, F., Pastuglia, M. and Bouchez, D. (2008) Arabidopsis TONNEAU1 proteins are essential for preprophase band formation and interact with centrin. Plant Cell, 20, 2146–2159.

47. Jay, F., Brioudes, F. and Voinnet, O. (2023) A contemporary reassessment of the enhanced transient expression system based on the tombusviral silencing suppressor protein P19. *Plant J*. Cell Mol. Biol., 113, 186–204.

48. Inagaki, S., Takahashi, M., Hosaka, A., Ito, T., Toyoda, A., Fujiyama, A., Tarutani, Y. and Kakutani, T. (2017) Gene-body chromatin modification dynamics mediate epigenome differentiation in Arabidopsis. EMBO J., 36, 970–980.

49. Blum, M., Andreeva, A., Florentino, L.C., Chuguransky, S.R., Grego, T., Hobbs, E., Pinto, B.L., Orr, A., Paysan-Lafosse, T., Ponamareva, I., et al. (2025) InterPro: the protein sequence classification resource in 2025. Nucleic Acids Res., 53, D444–D456.

50. Zheng, B., He, H., Zheng, Y., Wu, W. and McCormick, S. (2014) An ARID domain-containing protein within nuclear bodies is required for sperm cell formation in Arabidopsis thaliana. PLoS Genet., 10, e1004421.

51. Zhou, X., Fan, Y., Zhu, X., Zhao, R., He, J., Li, P., Shang, S., Goodrich, J., Zhu, J.-K. and Zhang, C.-J. (2024) SANT proteins modulate gene expression by coordinating histone H3KAc and Khib levels and regulate plant heat tolerance. Plant Physiol., 196, 902– 915.

52. Jumper, J., Evans, R., Pritzel, A., Green, T., Figurnov, M., Ronneberger, O., Tunyasuvunakool, K., Bates, R., Žídek, A., Potapenko, A., et al. (2021) Highly accurate protein structure prediction with AlphaFold. Nature, 596, 583–589.

53. He, X.-J., Hsu, Y.-F., Pontes, O., Zhu, J., Lu, J., Bressan, R.A., Pikaard, C., Wang, C.-S. and Zhu, J.-K. (2009) NRPD4, a protein related to the RPB4 subunit of RNA polymerase II, is a component of RNA polymerases IV and V and is required for RNA-directed DNA methylation. Genes Dev., 23, 318–330.

54. Zhang, S., Zhan, X., Xu, X., Cui, P., Zhu, J.-K., Xia, Y. and Xiong, L. (2015) Two domain-disrupted hda6 alleles have opposite epigenetic effects on transgenes and some endogenous targets. Sci. Rep., 5, 17832.

55. Wall, M.K., Mitchenall, L.A. and Maxwell, A. (2004) Arabidopsis thaliana DNA gyrase is targeted to chloroplasts and mitochondria. Proc. Natl. Acad. Sci. U. S. A., 101, 7821– 7826.

56. Dhabalia Ashok, A., de Vries, S., Darienko, T., Irisarri, I. and de Vries, J. (2024) Evolutionary assembly of the plant terrestrialization toolkit from protein domains. Proc. Biol. Sci., 291, 20240985.

57. Saharan, K., Baral, S., Shaikh, N.H. and Vasudevan, D. (2024) Structure-function analyses reveal Arabidopsis thaliana HDA7 to be an inactive histone deacetylase. Curr. Res. Struct. Biol., 7, 100136.

58. Low, J.K.K., Silva, A.P.G., Sharifi Tabar, M., Torrado, M., Webb, S.R., Parker, B.L., Sana, M., Smits, C., Schmidberger, J.W., Brillault, L., et al. (2020) The Nucleosome Remodeling and Deacetylase Complex Has an Asymmetric, Dynamic, and Modular Architecture. Cell Rep., 33, 108450.

59. Wang, S., Wang, M., Ichino, L., Boone, B.A., Zhong, Z., Papareddy, R.K., Lin, E.K., Yun, J., Feng, S. and Jacobsen, S.E. (2024) MBD2 couples DNA methylation to transposable element silencing during male gametogenesis. Nat. Plants, 10, 13–24.

60. Ning, Y.-Q., Chen, Q., Lin, R.-N., Li, Y.-Q., Li, L., Chen, S. and He, X.-J. (2019) The HDA19 histone deacetylase complex is involved in the regulation of flowering time in a photoperiod-dependent manner. *Plant J*. Cell Mol. Biol., 98, 448–464.

61. Xu, Y., Li, Q., Yuan, L., Huang, Y., Hung, F.-Y., Wu, K. and Yang, S. (2022) MSI1 and HDA6 function interdependently to control flowering time via chromatin modifications. *Plant J*. Cell Mol. Biol., 109, 831–843.

