## Supplementary Figures for "The Arabidopsis GyraseB3 contributes to transposon silencing by promoting histone deacetylation"

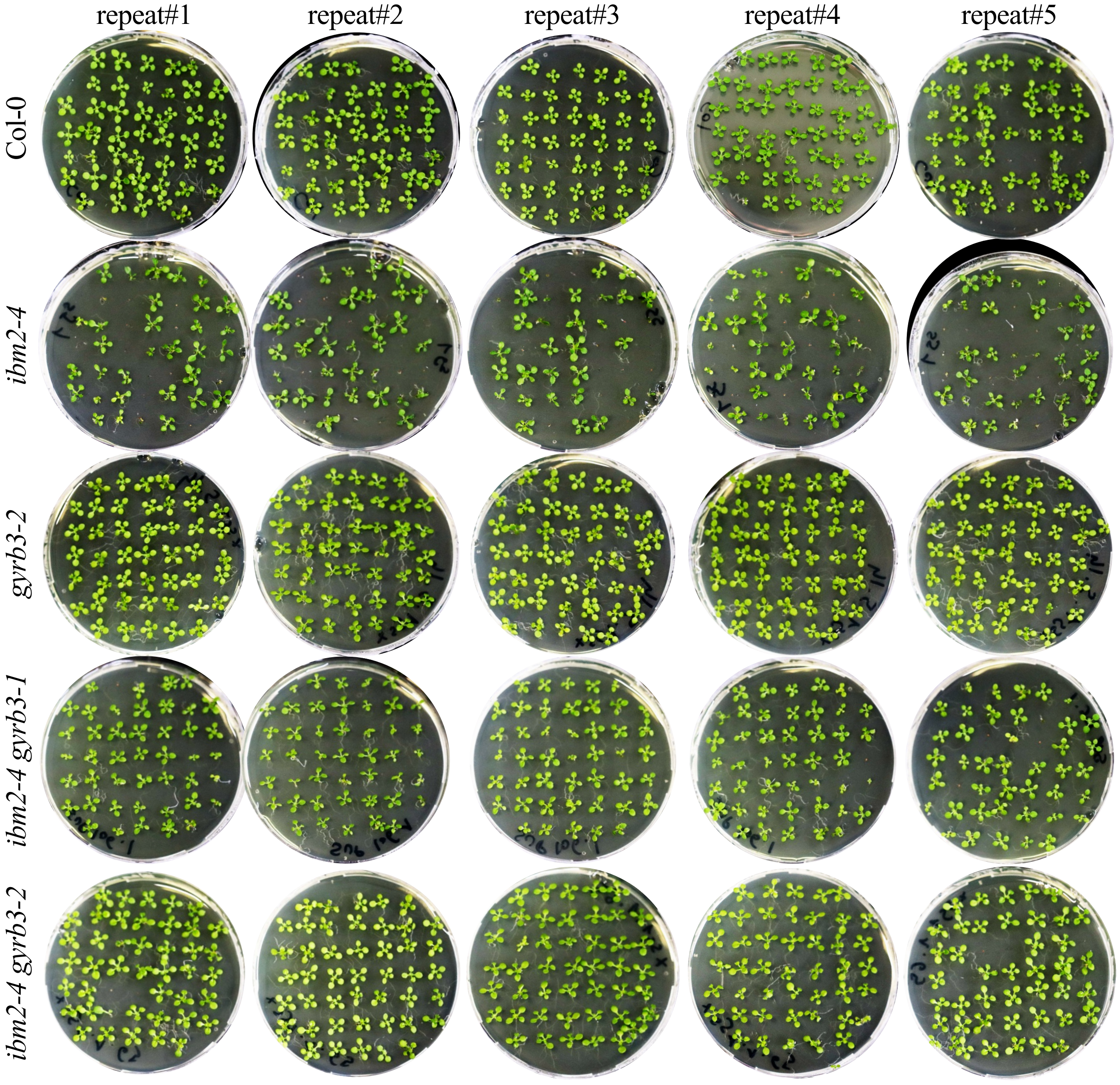


**Supplementary Fig. S1. Disrupting the GyrB3 function suppresses the developmental defects of *ibm2-4***

The *gyrb* mutations suppress the developmental abnormalities observed in *ibm2* mutants such as small, distorted leaves and reduced germination rate. Plants were pictured 15 days after sowing. The experiment was repeated five times; only repeat#1 is presented in Fig. 1, and all repeats are shown here. WT, Col-0.

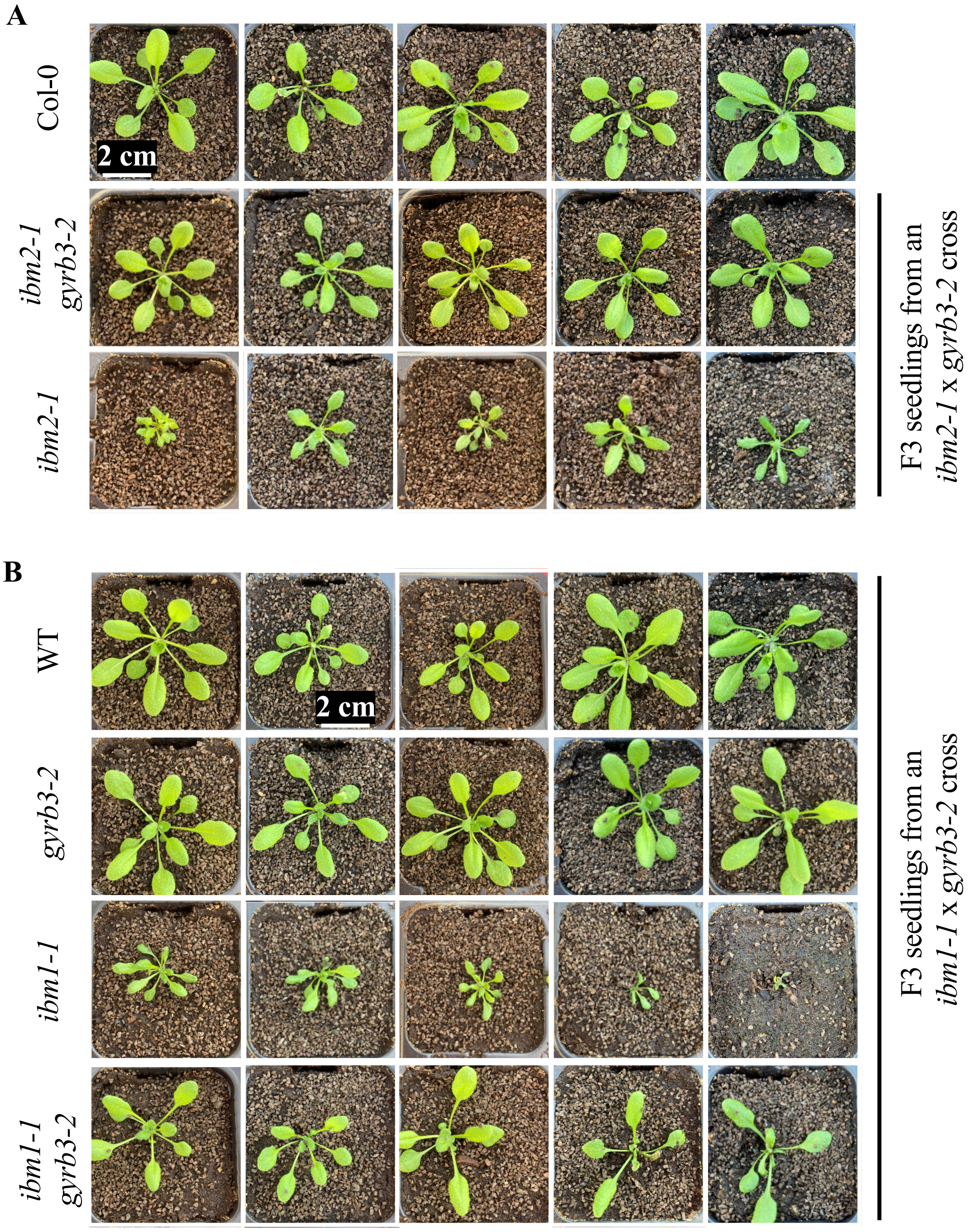


**Supplementary Fig. S2. *gyrb3* suppresses *ibm2-1* and *ibm1-1***

Pictures of F3 plants homozygous for *ibm2-1 gyrb3-2* or *ibm2-1* (A) and *gyrb3-2, ibm1-1* or *ibm1-1 gyrb3-2* (B) taken 21 days after sowing. Controls included Col-0 control plants (A) and F3 plants fixed for the wild-type (WT) *IBM1* and *GYRB3* alleles (B).

Plants were obtained through the self-pollination of F2 plants that were derived from an *ibm2-1* x *gyrb3-2* cross (A) or an *ibm1-1* x *gyrb3-2* cross (B). Five plants were grown together and only one is presented in Fig. 1C while all plants are shown here.

Mean leaf areas per plants are sown in Fig. 1D.

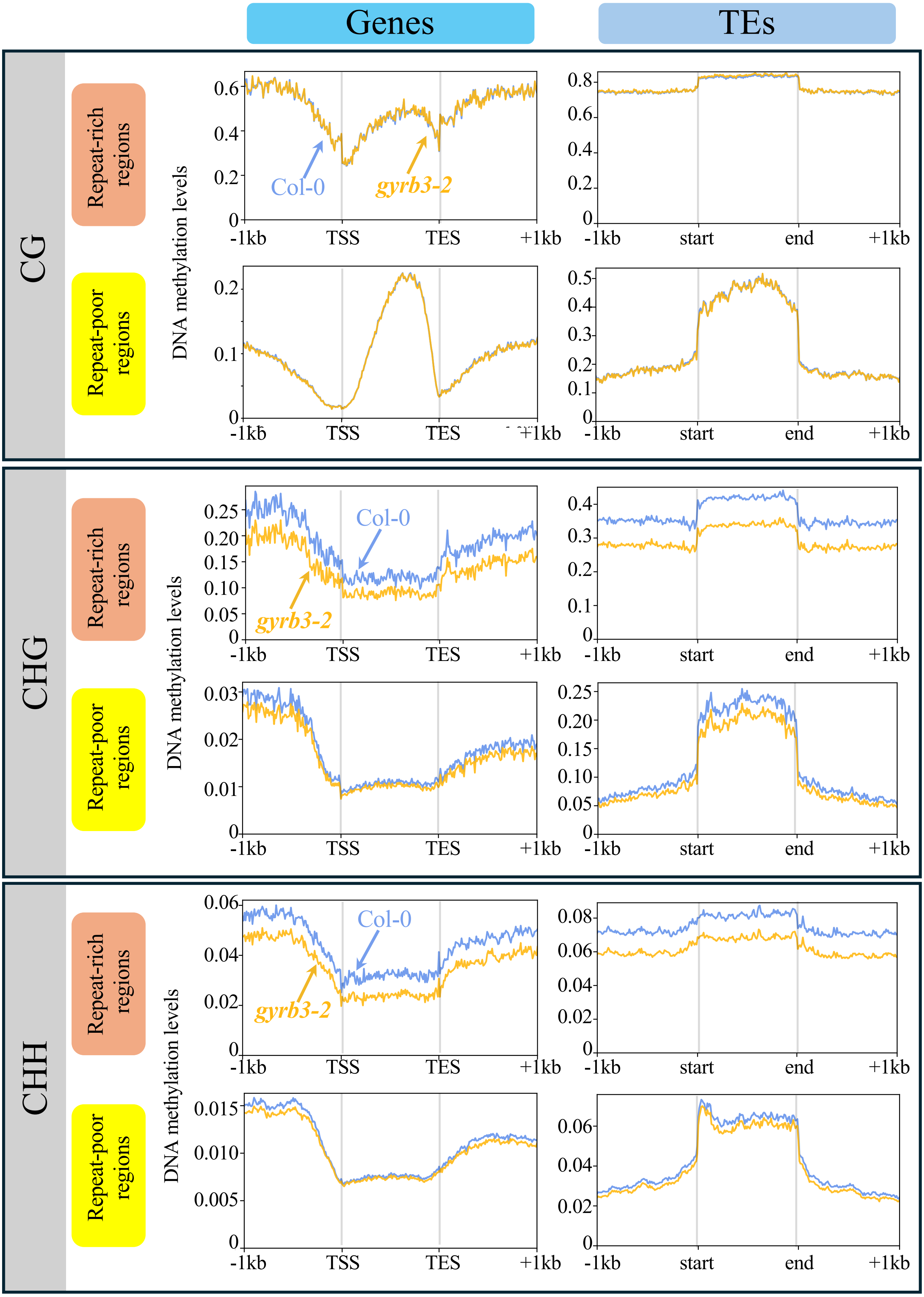


**Supplementary Fig. S3. Patterns of methylation of genes and TEs in Col-0 and *gyrb3-2***

The average methylation levels of genes (left panels) and TEs (right panels) were calculated by segmenting the corresponding annotated regions (TAIR10.57 release) into 100-bp bins. The metaplot analyses also include regions located 1 kb upstream and 1 kb downstream of the gene bodies and TEs. Repeat/TE-rich and repeat-poor (i.e. gene-rich) regions were defined as described in the *Materials and Methods* section.

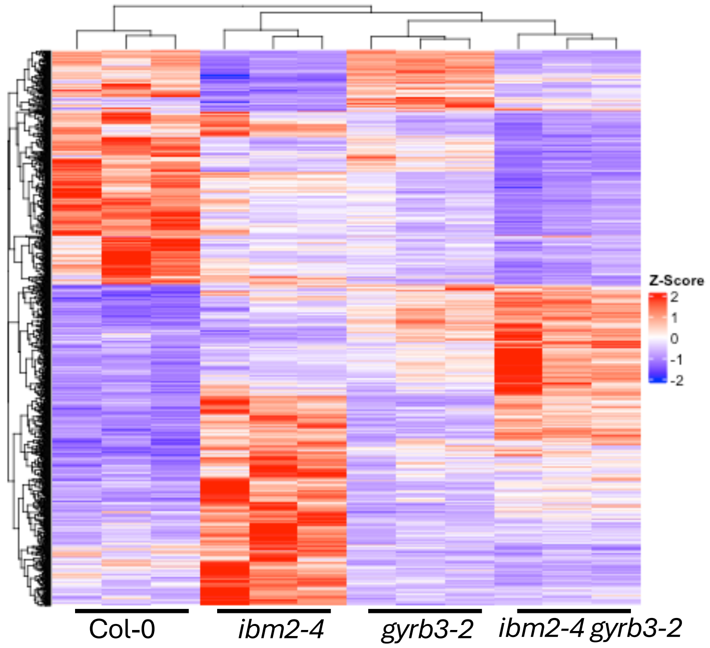


**Supplementary Fig. S4. Hierarchical clustering of genes differentially expressed between Col-0, *gyrb3-2*, *ibm2-4*, *ibm2-4 gyrb3-2***

Heatmap and clustering analysis of genes that are significantly (p-value < 0.05) upregulated (log2FC ≥ 1) or downregulated (log2FC ≤ -1) in at least one of the mutant genotypes compared to Col-0. Hierarchical clustering was performed after z-score transformation of the normalized counts (*DESeq2*) using the *scale* function of R.

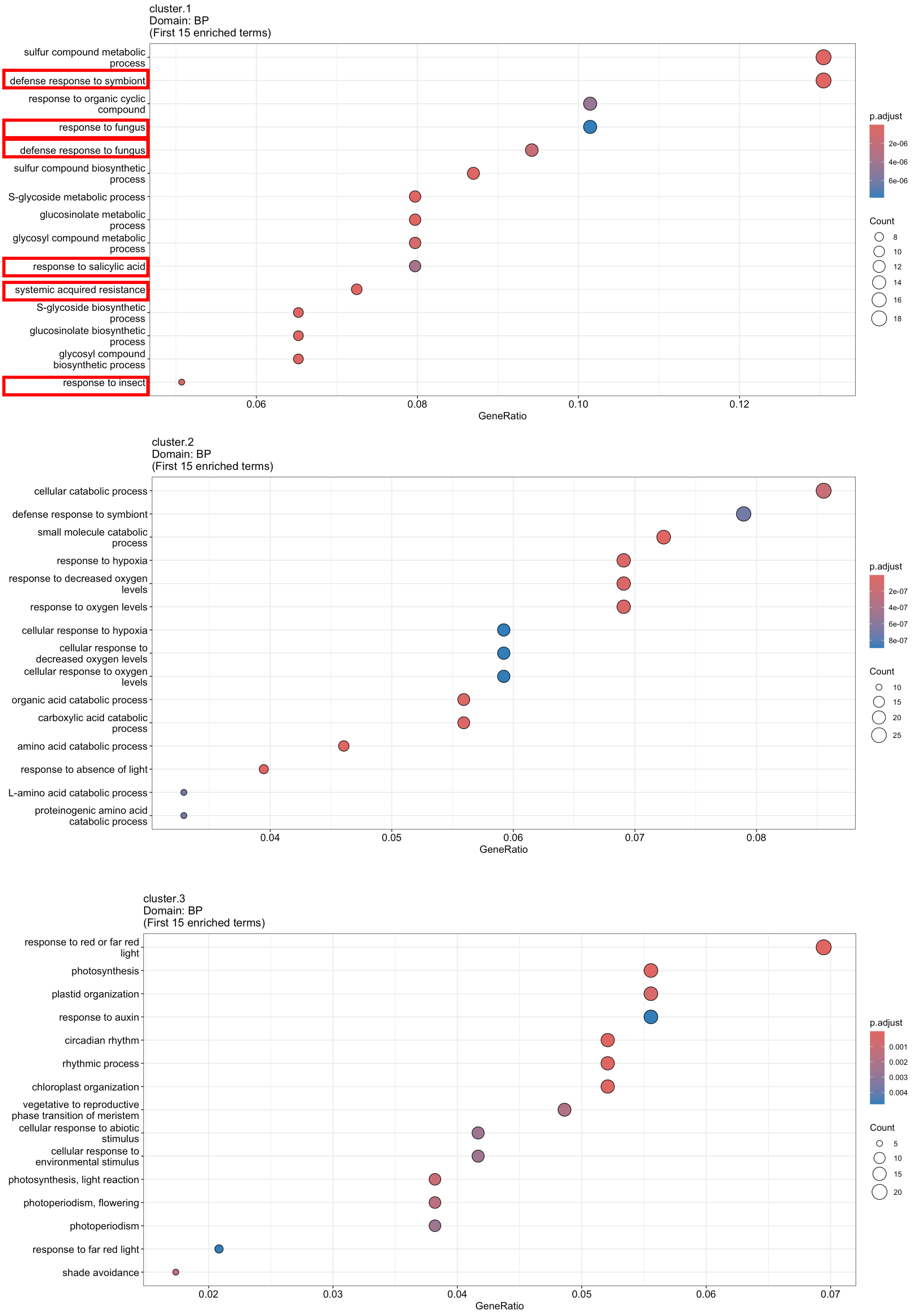


**Supplementary Fig. S5. RNAseq cluster gene ontology analysis focusing on the analysis of biological process (BP) terms**

Dot size represents the number of differentially expressed genes in the BP terms indicated on the left. Red triangles highlight BP terms associated with pathogen defence. Results obtained for the genes in the three clusters are presented.

**A**

| Protein | Function | E-value | % identity | Align len | Query from | Query to | Target from | Target to | Bit-score | # identical | Posi-tives | Gaps | Query len | Target len |
|---|---|---|---|---|---|---|---|---|---|---|---|---|---|---|
| AT5G04110.1 | GYRB3 | 0 | 100 | 546 | 1 | 546 | 1 | 546 | 1035.02 | 546 | 546 | 0 | 546 | 547 |
| AT5G04130.1 | DNA GYRASE B2 | 1.49E-83 | 62 | 252 | 17 | 260 | 288 | 536 | 275.018 | 155 | 192 | 11 | 546 | 733 |
| AT5G04130.2 | DNA GYRASE B2 | 4.05E-80 | 66 | 185 | 17 | 196 | 288 | 472 | 260.381 | 123 | 152 | 5 | 546 | 520 |
| AT3G10270.1 | DNA GYRASE B1 | 5.81E-78 | 55 | 284 | 17 | 260 | 296 | 576 | 261.151 | 156 | 193 | 43 | 546 | 773 |
| AT4G11400.1 | ARID/BRIGHT DNA-binding domain;ELM2 domain protein | 7.6E-29 | 38 | 169 | 358 | 523 | 371 | 524 | 120.553 | 65 | 98 | 18 | 546 | 574 |
| AT1G26580.1 | PTHR22970:SF28 - ELM2 DOMAIN-CONTAINING PROTEIN | 2.13E-19 | 32 | 176 | 356 | 521 | 130 | 302 | 91.2781 | 57 | 89 | 13 | 546 | 494 |
| AT2G46040.1 | ARID/BRIGHT DNA-binding domain;ELM2 domain protein | 7.8E-19 | 33 | 175 | 350 | 520 | 349 | 511 | 89.7373 | 57 | 88 | 16 | 546 | 563 |
| AT1G13880.1 | ELM2 domain-containing protein | 1.63E-16 | 32 | 175 | 358 | 521 | 125 | 284 | 82.0333 | 56 | 89 | 26 | 546 | 425 |
| AT2G03470.1 | ELM2 domain-containing protein | 1.83E-16 | 30 | 196 | 329 | 521 | 89 | 278 | 82.0333 | 59 | 93 | 9 | 546 | 451 |
| AT2G03470.2 | ELM2 domain-containing protein | 2E-16 | 30 | 196 | 329 | 521 | 88 | 277 | 81.6481 | 59 | 93 | 9 | 546 | 450 |

**B**

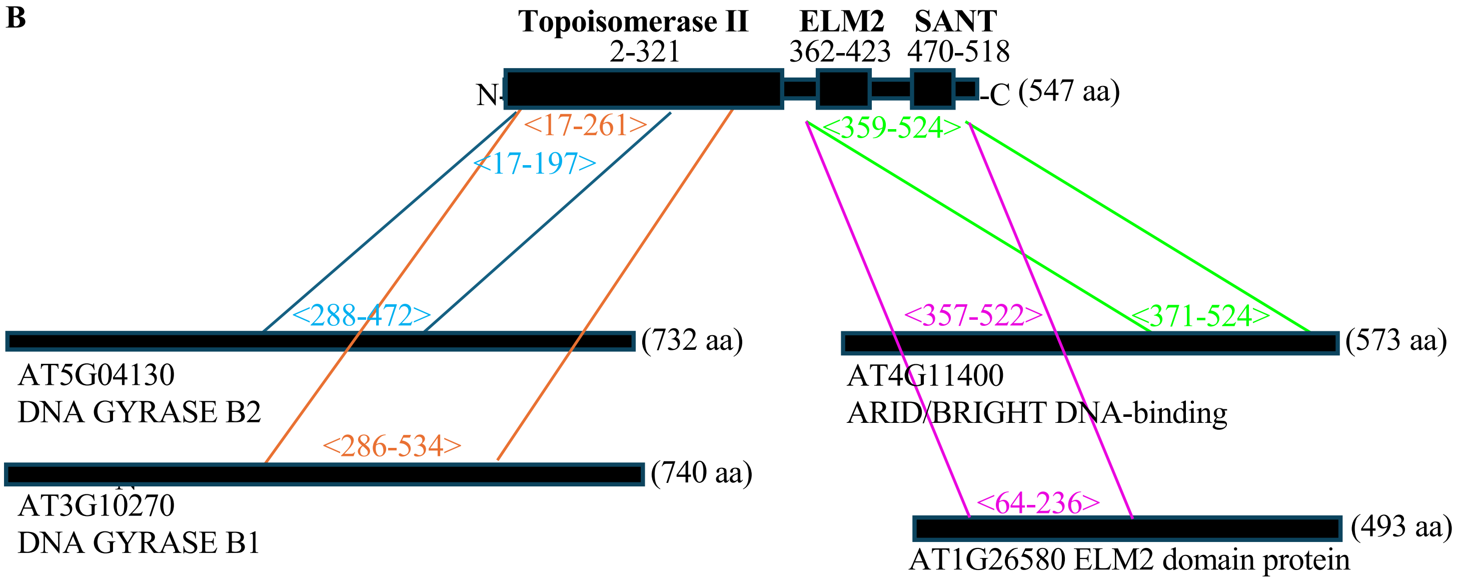


**Supplementary Fig. S6. GyrB3 shares domains with ARID- and ELM2/SANT-containing proteins**
(A) GyrB3 protein sequence alignment against the Arabidopsis proteome. Spreadsheet view of BLASTP results classified by percentage of identity compared with GyrB3. BlastP searches were conducted using the *Phytozome* platform.
(B) Homologies found between the different domains of GyrB3, and proteins listed in (A).

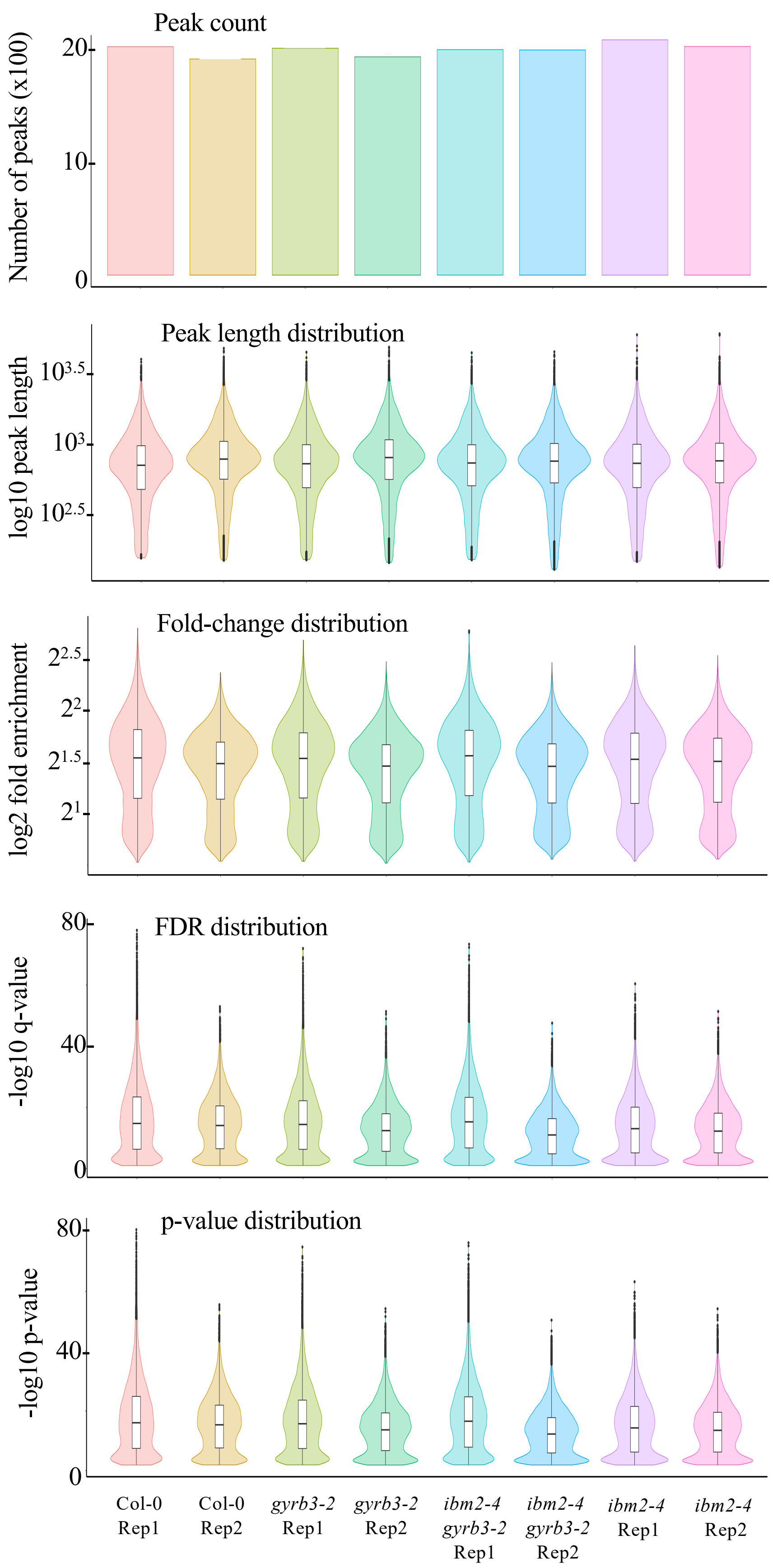


**Supplementary Fig. S7. Description of the H3Kac peaks identified in the different genotypes**

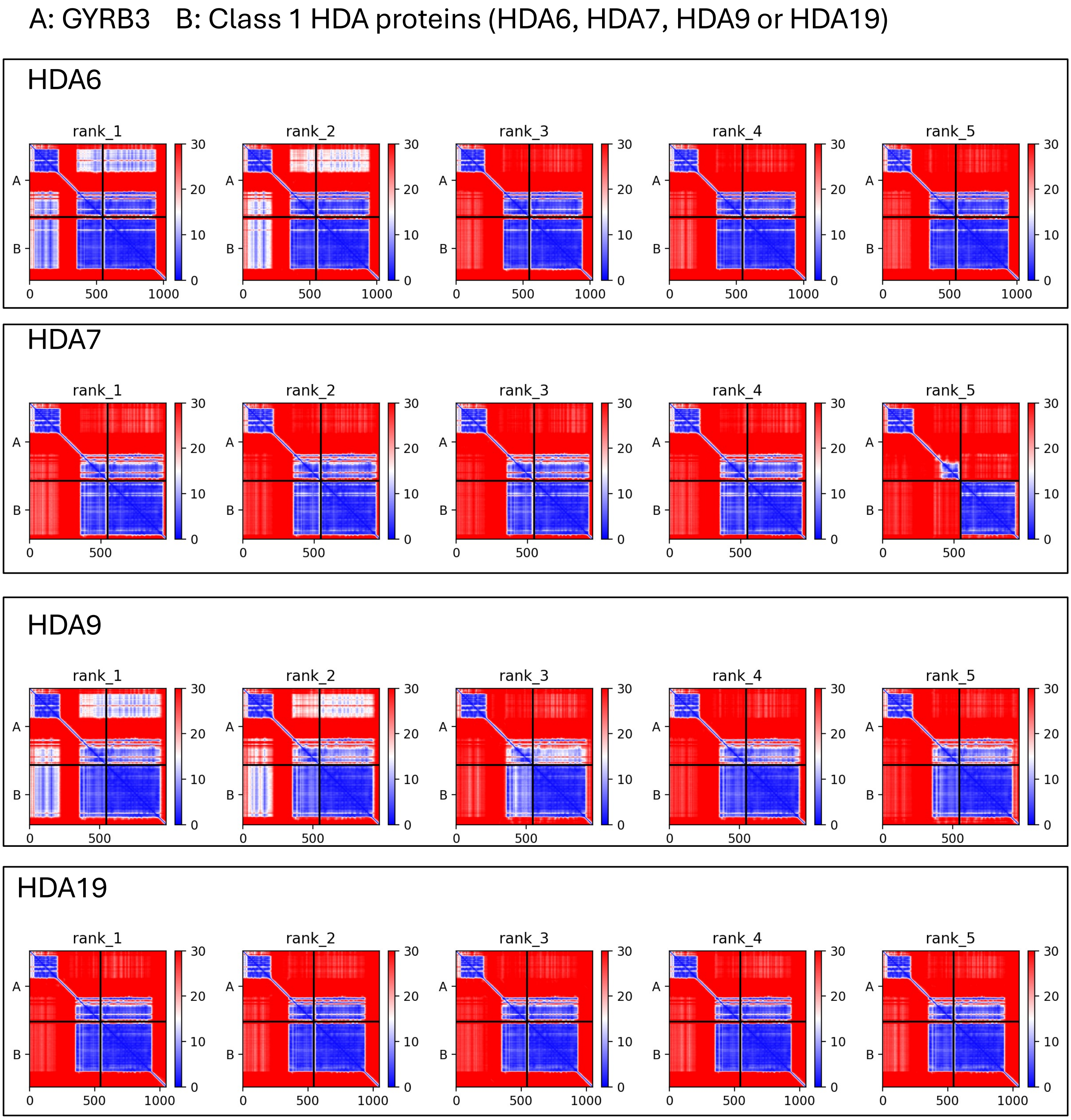


**Supplementary Fig. S8. 2D Plot of Predicted Aligned Error (PAE) for Protein Interactions with GyrB3**
Two-dimensional plot illustrating the Predicted Aligned Error (PAE) data from AFM outputs using HDA6, HDA7, HDA9, and HDA19 and GyrB3 as input proteins. The analysis predicts that all histone deacetylases are predicted to interact with GyrB3.

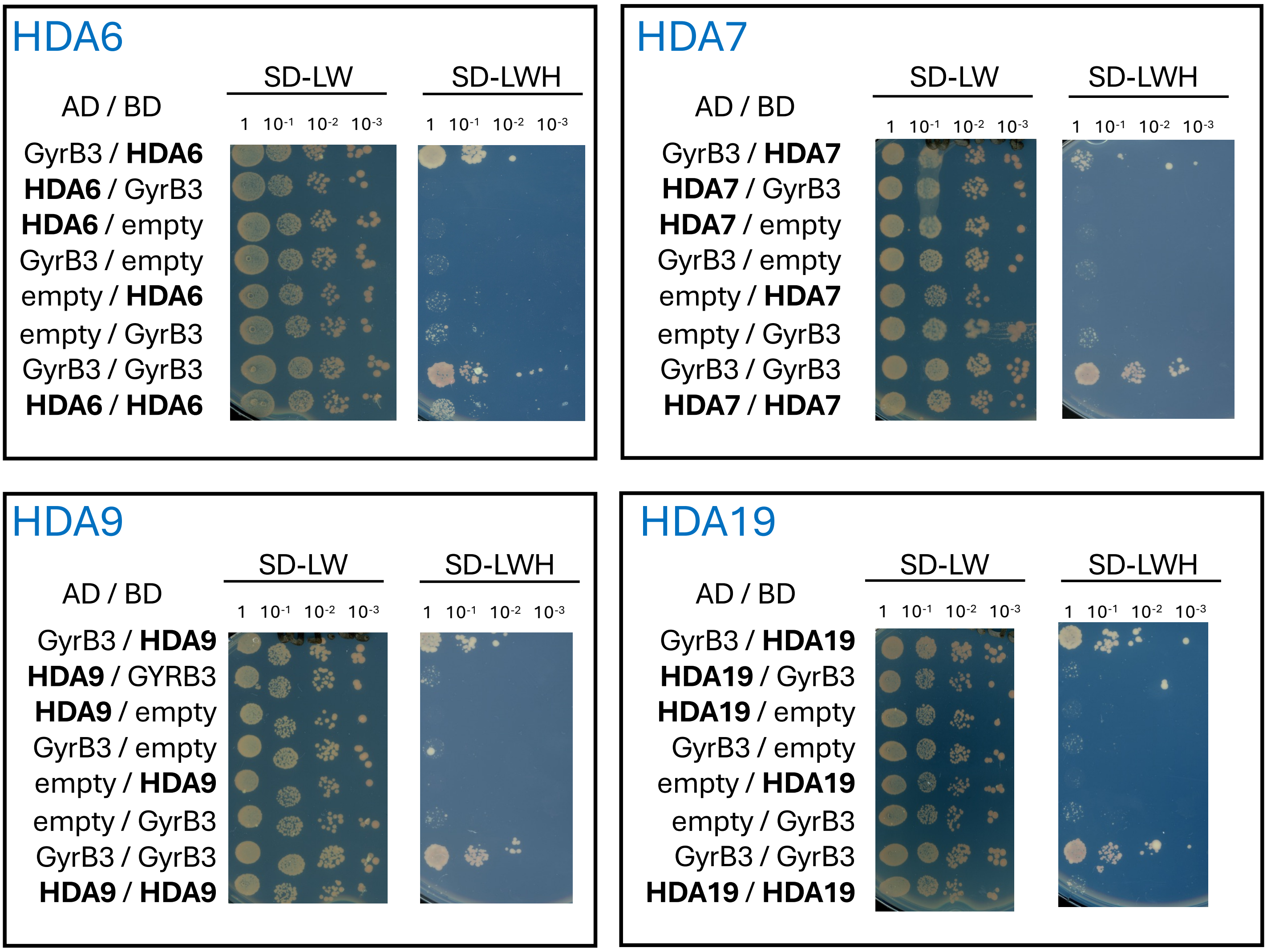


**Supplementary Fig. S9. Repeat of the yeast-two-hybrid assays testing interactions between HDA6, HDA7, HDA9, HDA19 and GyrB3 (Fig. 6D)**

Full length proteins were fused with Gal4 DNA binding domain (BD) and Gal4 activation domain (AD), respectively, and co-expressed in yeast cells. For each combination, serial dilutions of yeast cells were spotted on non-selective medium (-LW), or selective media (-LWH).

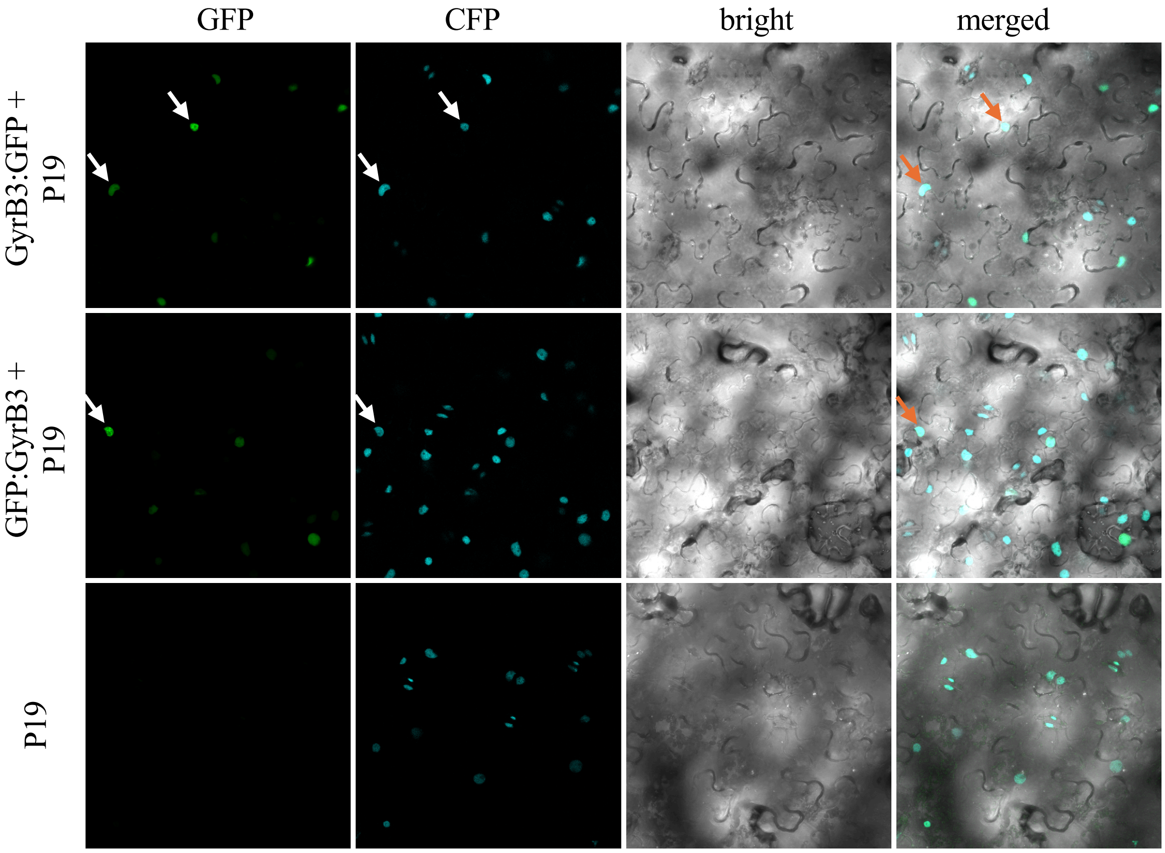


**Supplementary Fig. S10. Nuclear localization of GyrB3:GFP and GFP:GyrB3 fusion proteins in tobacco cells**

Subcellular localization patterns were examined in the epidermal cells of *Nicotiana benthamiana* leaves from transgenic plants stably expressing a nuclear CFP:Histone 2B fusion protein as a nuclear marker (45), following agroinfiltration with *a p35S::GyrB3:GFP* (GyrB3:GFP) construct or *p35S::GFP:GyrB3* (GFP:GyrB3) and a *p35S::P19* (P19) construct, which enhances transient expression (47). Fluorescence was analyzed 48 hours after infiltration. Arrows indicate representative nuclei.

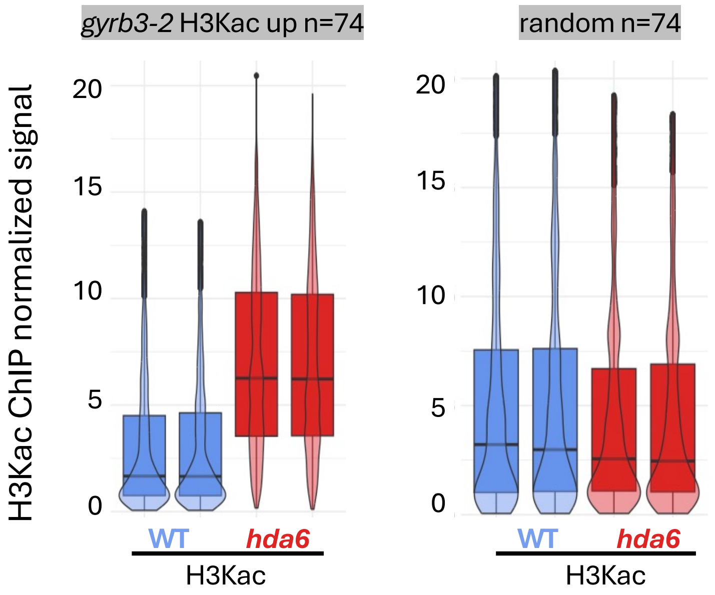


**Supplementary Fig. S11. Genomic regions exhibiting increased H3Kac levels in *gyrb3* also show elevated H3Kac accumulation in *hda6* mutant**

H3Kac levels in two wild-type (WT) and two *hda6* biological repeats in regions accumulating H3Kac in *gyrb3-2* (n=74). ChIP signals were quantified using normalized *BigWig* files scaled to 1 million mapped reads. The data (43) were retrieved from the GEO repository (accession number GSE132636). As a control, ChIP signals were measured in randomly selected regions of similar length.

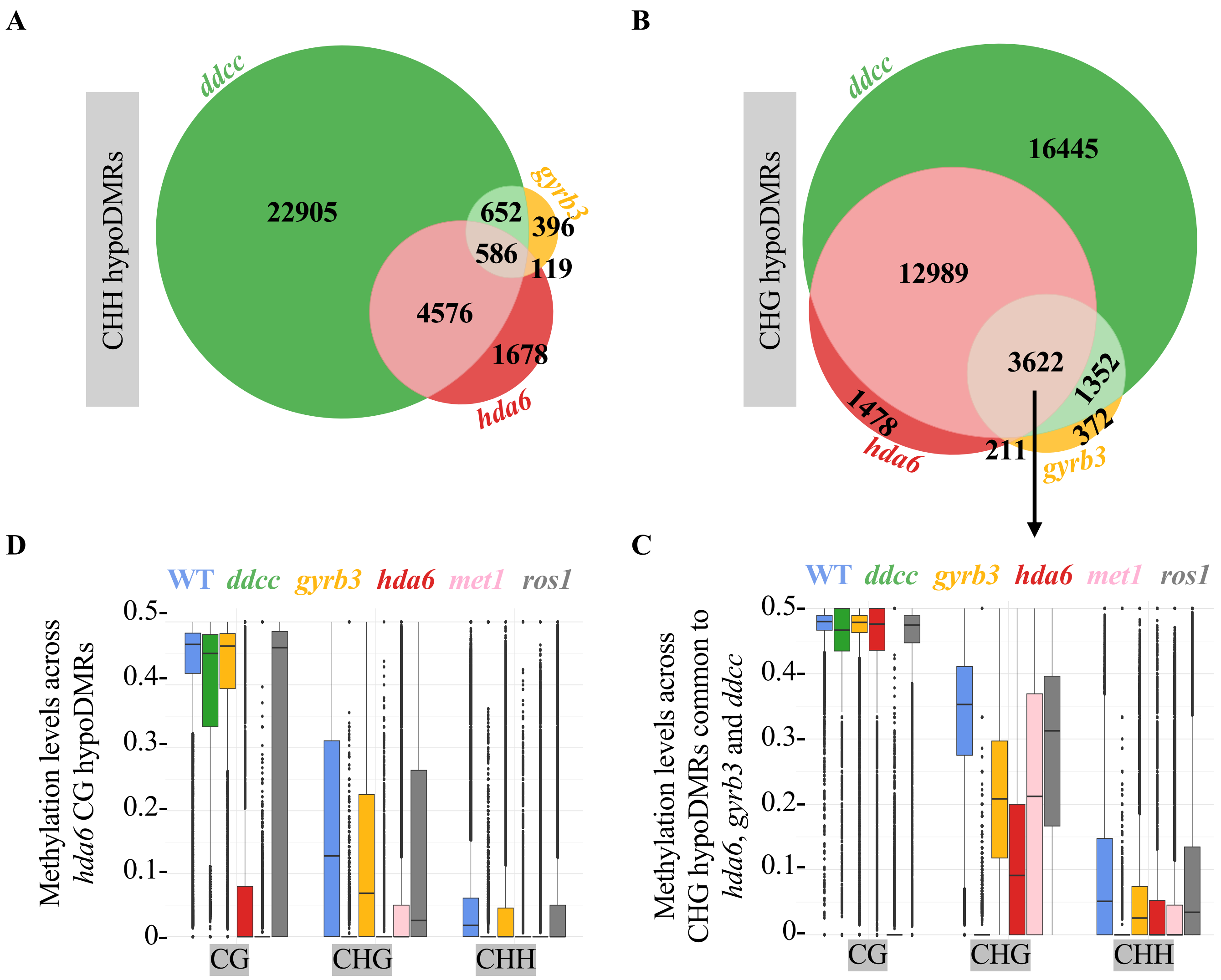


**Supplementary Fig. S12. *gyrb3* and *hda6* CHG hypoDMRs overlap and are controlled by common epigenetic pathways**

(A) Venn diagram showing overlaps between CHH hypoDMRs identified in *gyrb3-2* with those found in *hda6* (*axe1-5*) and the *drm1 drm2 cmt3 cmt2* quadruple mutant (*ddcc*) in which all non-CG methylation is abolished.

(B) Venn diagram showing overlaps between CHG hypoDMRs identified in *gyrb3-2* with those found in *hda6* (*axe1-5*) and *ddcc*. The CHG hypoDMRs used in (C) are indicated.

(C) DNA methylation levels across CHG hypoDMRs common to *gyrb3*, *hda6* and *ddcc* (n=3622) in the wild-type (WT), *ddcc*, *gyrb3-2*, *hda6* (*axe1-5*), *met1-9*, and *ros1-1*.

(D) DNA methylation levels across *hda6* CG hypoDMRs (n=5997) in the wild-type (WT), *ddcc*, *gyrb3-2*, *hda6* (*axe1-5*), *met1-9*, and *ros1-1*.

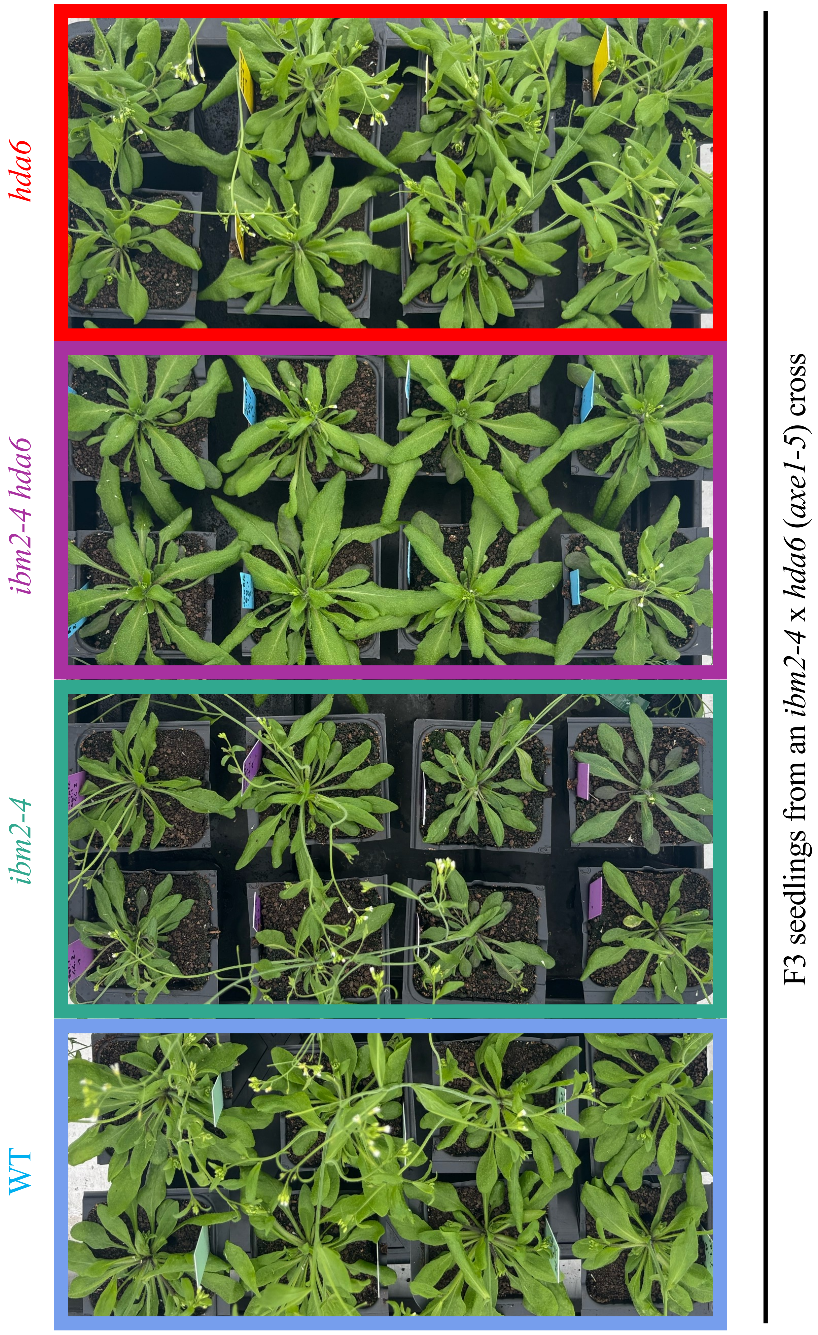


**Supplementary Fig. S13. A mutation in *HDA6* suppresses the developmental defect of *ibm2***

Pictures of the wild-type (WT), *ibm2-4*, *hda6* (*axe1-5*) and the double *ibm2-4 hda6* F3 plants taken 37 days after sowing. Lines were obtained by self-pollinating F2 plants from a cross between *ibm2-4* and *hda6*. All plants were individually genotyped for the mutations in both F2s and F3s.

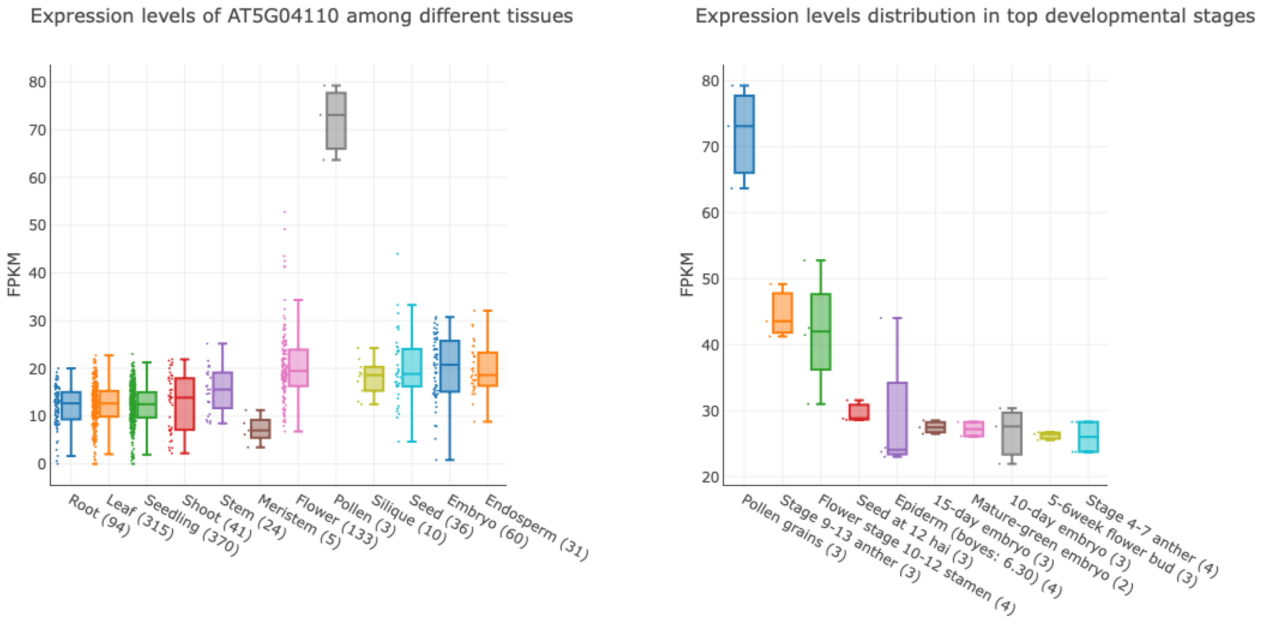


**Supplementary Fig. S14. *GYRB3* is expressed in pollen**

Mean expression levels (FPKM) of the *GYRB3* gene retrieved from the Arabidopsis RNA-seq Database (https://plantrnadb.com/athrdb/). The number in parentheses indicates the number of public Arabidopsis RNA-Seq libraries.
