## Supplementary Tables for "The Arabidopsis GyraseB3 contributes to transposon silencing by promoting histone deacetylation"

**Supplementary Table S1**

**Whole-genome bisulfite sequencing statistics**

| | ***ibm2-1* repeat#1** | ***ibm2-1* repeat#2** | ***ibm2-1 gyrb3-2* repeat#1** | ***ibm2-1 gyrb3-2* repeat#2** | **Col-0 repeat#1** | **Col-0 repeat#2** | ***gyrb3-2* repeat#1** | ***gyrb3-2* repeat#2** |
|---|---|---|---|---|---|---|---|---|
| Number of clean paired-end reads (after trimming | 36069112 | 36051044 | 36042275 | 36062779 | 36107066 | 36085347 | 36077206 | 36118721 |
| Number of mapped paired-end reads | 24955426 | 24555856 | 24432959 | 24690977 | 26280417 | 25991099 | 26156128 | 26350449 |
| Fold coverage | 36.6 | 35.9 | 35.9 | 36.3 | 40.2 | 39.9 | 40.1 | 40.3 |
| Number of Cs in CG context | 29321127 | 28751913 | 29990673 | 30534166 | 29736976 | 30849930 | 30722996 | 29113182 |
| Number of Cs in CHG context | 15159355 | 14979006 | 12673199 | 12956225 | 10693240 | 11094445 | 9206692 | 8668420 |
| Number of Cs in CHH context | 16362291 | 16377891 | 14150386 | 13995491 | 15865373 | 15687925 | 14165708 | 13212439 |
| Bisulfite conversion rates | 0.99 | 0.99 | 0.99 | 0.99 | 0.99 | 0.99 | 0.99 | 0.99 |

The number of paired-end reads obtained after trimming, the number of mapped paired-end reads, the coverage, and the numbers of cytosines in all contexts are indicated. The percentages of methylated cytosines is indicated in parentheses. Bisulfite conversion rates were determined by aligning the reads to the chloroplast sequence of Arabidopsis.

**Supplementary Table S2**

**Classification of the genomic regions according to their T**

RP, Repeat-poor regions; RR, Repeat-rich regions; INT, intermediate.

| chr | start | end | Region |
|---|---|---|---|
| 1 | 1 | 500000 | RP |
| 1 | 450001 | 950000 | RP |
| 1 | 900001 | 1400000 | RP |
| 1 | 1350001 | 1850000 | RP |
| 1 | 1800001 | 2300000 | RP |
| 1 | 2250001 | 2750000 | RP |
| 1 | 2700001 | 3200000 | RP |
| 1 | 3150001 | 3650000 | RP |
| 1 | 3600001 | 4100000 | RP |
| 1 | 4050001 | 4550000 | RP |
| 1 | 4500001 | 5000000 | RP |
| 1 | 4950001 | 5450000 | RP |
| 1 | 5400001 | 5900000 | RP |
| 1 | 5850001 | 6350000 | RP |
| 1 | 6300001 | 6800000 | RP |
| 1 | 6750001 | 7250000 | RP |
| 1 | 7200001 | 7700000 | RP |
| 1 | 7650001 | 8150000 | RP |
| 1 | 8100001 | 8600000 | RP |
| 1 | 8550001 | 9050000 | RP |
| 1 | 9000001 | 9500000 | RP |
| 1 | 9450001 | 9950000 | RP |
| 1 | 9900001 | 10400000 | RP |
| 1 | 10350001 | 10850000 | RP |
| 1 | 10800001 | 11300000 | INT |
| 1 | 11250001 | 11750000 | INT |
| 1 | 11700001 | 12200000 | INT |
| 1 | 12150001 | 12650000 | INT |
| 1 | 12600001 | 13100000 | RR |
| 1 | 13050001 | 13550000 | RR |
| 1 | 13500001 | 14000000 | RR |
| 1 | 13950001 | 14450000 | RR |
| 1 | 14400001 | 14900000 | RR |
| 1 | 14850001 | 15350000 | RR |
| 1 | 15300001 | 15800000 | RR |
| 1 | 15750001 | 16250000 | RR |
| 1 | 16200001 | 16700000 | RR |
| 1 | 16650001 | 17150000 | INT |
| 1 | 17100001 | 17600000 | INT |
| 1 | 17550001 | 18050000 | INT |
| 1 | 18000001 | 18500000 | RP |
| 1 | 18450001 | 18950000 | RP |

| | | | |
|---|---|---|---|
| 1 | 18900001 | 19400000 | INT |
| 1 | 19350001 | 19850000 | INT |
| 1 | 19800001 | 20300000 | RP |
| 1 | 20250001 | 20750000 | RP |
| 1 | 20700001 | 21200000 | RP |
| 1 | 21150001 | 21650000 | INT |
| 1 | 21600001 | 22100000 | INT |
| 1 | 22050001 | 22550000 | RP |
| 1 | 22500001 | 23000000 | RP |
| 1 | 22950001 | 23450000 | INT |
| 1 | 23400001 | 23900000 | RP |
| 1 | 23850001 | 24350000 | RP |
| 1 | 24300001 | 24800000 | INT |
| 1 | 24750001 | 25250000 | RP |
| 1 | 25200001 | 25700000 | RP |
| 1 | 25650001 | 26150000 | RP |
| 1 | 26100001 | 26600000 | RP |
| 1 | 26550001 | 27050000 | RP |
| 1 | 27000001 | 27500000 | RP |
| 1 | 27450001 | 27950000 | RP |
| 1 | 27900001 | 28400000 | RP |
| 1 | 28350001 | 28850000 | RP |
| 1 | 28800001 | 29300000 | RP |
| 1 | 29250001 | 29750000 | RP |
| 1 | 29700001 | 30200000 | RP |
| 1 | 30150001 | 30427671 | RP |
| 2 | 1 | 500000 | INT |
| 2 | 450001 | 950000 | INT |
| 2 | 900001 | 1400000 | INT |
| 2 | 1350001 | 1850000 | INT |
| 2 | 1800001 | 2300000 | RR |
| 2 | 2250001 | 2750000 | RR |
| 2 | 2700001 | 3200000 | RR |
| 2 | 3150001 | 3650000 | INT |
| 2 | 3600001 | 4100000 | RR |
| 2 | 4050001 | 4550000 | RR |
| 2 | 4500001 | 5000000 | RR |
| 2 | 4950001 | 5450000 | RR |
| 2 | 5400001 | 5900000 | RR |
| 2 | 5850001 | 6350000 | RR |
| 2 | 6300001 | 6800000 | RR |
| 2 | 6750001 | 7250000 | INT |
| 2 | 7200001 | 7700000 | INT |
| 2 | 7650001 | 8150000 | INT |
| 2 | 8100001 | 8600000 | RP |
| 2 | 8550001 | 9050000 | RP |
| 2 | 9000001 | 9500000 | RP |
| 2 | 9450001 | 9950000 | INT |

| | | | |
|---|---|---|---|
| 2 | 9900001 | 10400000 | INT |
| 2 | 10350001 | 10850000 | INT |
| 2 | 10800001 | 11300000 | RP |
| 2 | 11250001 | 11750000 | RP |
| 2 | 11700001 | 12200000 | RP |
| 2 | 12150001 | 12650000 | RP |
| 2 | 12600001 | 13100000 | RP |
| 2 | 13050001 | 13550000 | RP |
| 2 | 13500001 | 14000000 | RP |
| 2 | 13950001 | 14450000 | RP |
| 2 | 14400001 | 14900000 | RP |
| 2 | 14850001 | 15350000 | RP |
| 2 | 15300001 | 15800000 | RP |
| 2 | 15750001 | 16250000 | RP |
| 2 | 16200001 | 16700000 | RP |
| 2 | 16650001 | 17150000 | RP |
| 2 | 17100001 | 17600000 | RP |
| 2 | 17550001 | 18050000 | RP |
| 2 | 18000001 | 18500000 | RP |
| 2 | 18450001 | 18950000 | RP |
| 2 | 18900001 | 19400000 | RP |
| 2 | 19350001 | 19698289 | RP |
| 3 | 1 | 500000 | RP |
| 3 | 450001 | 950000 | RP |
| 3 | 900001 | 1400000 | RP |
| 3 | 1350001 | 1850000 | RP |
| 3 | 1800001 | 2300000 | RP |
| 3 | 2250001 | 2750000 | RP |
| 3 | 2700001 | 3200000 | RP |
| 3 | 3150001 | 3650000 | RP |
| 3 | 3600001 | 4100000 | RP |
| 3 | 4050001 | 4550000 | RP |
| 3 | 4500001 | 5000000 | RP |
| 3 | 4950001 | 5450000 | RP |
| 3 | 5400001 | 5900000 | RP |
| 3 | 5850001 | 6350000 | RP |
| 3 | 6300001 | 6800000 | RP |
| 3 | 6750001 | 7250000 | RP |
| 3 | 7200001 | 7700000 | RP |
| 3 | 7650001 | 8150000 | INT |
| 3 | 8100001 | 8600000 | INT |
| 3 | 8550001 | 9050000 | RP |
| 3 | 9000001 | 9500000 | INT |
| 3 | 9450001 | 9950000 | INT |
| 3 | 9900001 | 10400000 | INT |
| 3 | 10350001 | 10850000 | INT |
| 3 | 10800001 | 11300000 | INT |
| 3 | 11250001 | 11750000 | RR |

| | | | |
|---|---|---|---|
| 3 | 11700001 | 12200000 | RR |
| 3 | 12150001 | 12650000 | RR |
| 3 | 12600001 | 13100000 | RR |
| 3 | 13050001 | 13550000 | RR |
| 3 | 13500001 | 14000000 | RR |
| 3 | 13950001 | 14450000 | RR |
| 3 | 14400001 | 14900000 | RR |
| 3 | 14850001 | 15350000 | RR |
| 3 | 15300001 | 15800000 | RR |
| 3 | 15750001 | 16250000 | INT |
| 3 | 16200001 | 16700000 | INT |
| 3 | 16650001 | 17150000 | INT |
| 3 | 17100001 | 17600000 | INT |
| 3 | 17550001 | 18050000 | RP |
| 3 | 18000001 | 18500000 | RP |
| 3 | 18450001 | 18950000 | INT |
| 3 | 18900001 | 19400000 | RP |
| 3 | 19350001 | 19850000 | RP |
| 3 | 19800001 | 20300000 | RP |
| 3 | 20250001 | 20750000 | RP |
| 3 | 20700001 | 21200000 | RP |
| 3 | 21150001 | 21650000 | RP |
| 3 | 21600001 | 22100000 | RP |
| 3 | 22050001 | 22550000 | RP |
| 3 | 22500001 | 23000000 | RP |
| 3 | 22950001 | 23450000 | RP |
| 3 | 23400001 | 23459830 | RP |
| 4 | 1 | 500000 | RP |
| 4 | 450001 | 950000 | RP |
| 4 | 900001 | 1400000 | RP |
| 4 | 1350001 | 1850000 | INT |
| 4 | 1800001 | 2300000 | RR |
| 4 | 2250001 | 2750000 | INT |
| 4 | 2700001 | 3200000 | RR |
| 4 | 3150001 | 3650000 | RR |
| 4 | 3600001 | 4100000 | RR |
| 4 | 4050001 | 4550000 | RR |
| 4 | 4500001 | 5000000 | RR |
| 4 | 4950001 | 5450000 | RR |
| 4 | 5400001 | 5900000 | INT |
| 4 | 5850001 | 6350000 | INT |
| 4 | 6300001 | 6800000 | INT |
| 4 | 6750001 | 7250000 | INT |
| 4 | 7200001 | 7700000 | INT |
| 4 | 7650001 | 8150000 | INT |
| 4 | 8100001 | 8600000 | RP |
| 4 | 8550001 | 9050000 | RP |
| 4 | 9000001 | 9500000 | RP |

| | | | |
|---|---|---|---|
| 4 | 9450001 | 9950000 | RP |
| 4 | 9900001 | 10400000 | RP |
| 4 | 10350001 | 10850000 | RP |
| 4 | 10800001 | 11300000 | INT |
| 4 | 11250001 | 11750000 | RP |
| 4 | 11700001 | 12200000 | RP |
| 4 | 12150001 | 12650000 | RP |
| 4 | 12600001 | 13100000 | RP |
| 4 | 13050001 | 13550000 | RP |
| 4 | 13500001 | 14000000 | RP |
| 4 | 13950001 | 14450000 | RP |
| 4 | 14400001 | 14900000 | RP |
| 4 | 14850001 | 15350000 | RP |
| 4 | 15300001 | 15800000 | RP |
| 4 | 15750001 | 16250000 | RP |
| 4 | 16200001 | 16700000 | RP |
| 4 | 16650001 | 17150000 | RP |
| 4 | 17100001 | 17600000 | RP |
| 4 | 17550001 | 18050000 | RP |
| 4 | 18000001 | 18500000 | RP |
| 4 | 18450001 | 18585056 | RP |
| 5 | 1 | 500000 | RP |
| 5 | 450001 | 950000 | RP |
| 5 | 900001 | 1400000 | RP |
| 5 | 1350001 | 1850000 | RP |
| 5 | 1800001 | 2300000 | RP |
| 5 | 2250001 | 2750000 | RP |
| 5 | 2700001 | 3200000 | RP |
| 5 | 3150001 | 3650000 | RP |
| 5 | 3600001 | 4100000 | RP |
| 5 | 4050001 | 4550000 | RP |
| 5 | 4500001 | 5000000 | RP |
| 5 | 4950001 | 5450000 | RP |
| 5 | 5400001 | 5900000 | RP |
| 5 | 5850001 | 6350000 | RP |
| 5 | 6300001 | 6800000 | RP |
| 5 | 6750001 | 7250000 | RP |
| 5 | 7200001 | 7700000 | RP |
| 5 | 7650001 | 8150000 | RP |
| 5 | 8100001 | 8600000 | RP |
| 5 | 8550001 | 9050000 | INT |
| 5 | 9000001 | 9500000 | INT |
| 5 | 9450001 | 9950000 | INT |
| 5 | 9900001 | 10400000 | RR |
| 5 | 10350001 | 10850000 | RR |
| 5 | 10800001 | 11300000 | RR |
| 5 | 11250001 | 11750000 | RR |
| 5 | 11700001 | 12200000 | RR |

| | | | |
|---|---|---|---|
| 5 | 12150001 | 12650000 | RR |
| 5 | 12600001 | 13100000 | RR |
| 5 | 13050001 | 13550000 | RR |
| 5 | 13500001 | 14000000 | RR |
| 5 | 13950001 | 14450000 | RR |
| 5 | 14400001 | 14900000 | INT |
| 5 | 14850001 | 15350000 | INT |
| 5 | 15300001 | 15800000 | INT |
| 5 | 15750001 | 16250000 | RP |
| 5 | 16200001 | 16700000 | RP |
| 5 | 16650001 | 17150000 | INT |
| 5 | 17100001 | 17600000 | INT |
| 5 | 17550001 | 18050000 | RP |
| 5 | 18000001 | 18500000 | RP |
| 5 | 18450001 | 18950000 | RP |
| 5 | 18900001 | 19400000 | RP |
| 5 | 19350001 | 19850000 | RP |
| 5 | 19800001 | 20300000 | RP |
| 5 | 20250001 | 20750000 | RP |
| 5 | 20700001 | 21200000 | RP |
| 5 | 21150001 | 21650000 | RP |
| 5 | 21600001 | 22100000 | RP |
| 5 | 22050001 | 22550000 | RP |
| 5 | 22500001 | 23000000 | RP |
| 5 | 22950001 | 23450000 | RP |
| 5 | 23400001 | 23900000 | RP |
| 5 | 23850001 | 24350000 | RP |
| 5 | 24300001 | 24800000 | RP |
| 5 | 24750001 | 25250000 | RP |
| 5 | 25200001 | 25700000 | RP |
| 5 | 25650001 | 26150000 | RP |
| 5 | 26100001 | 26600000 | RP |
| 5 | 26550001 | 26975502 | RP |

**Supplementary Table S3**

***ibm2-4* suppressor mutations identified in the genetic screen**

Supp#11 (*fpa* ) was described previously (Deremetz et al. 2019).

| Candidate ID | Nucleotide change | Changes | Chromosome Position | Gene description |
|---|---|---|---|---|
| supp#1 SUP85 | G->A | G224R | Chr5: 4502904 | *KRYPTONITE* (*KYP*) |
| supp#2 SUP103 | G->A | R126K | Chr5: 4502168 | *KRYPTONITE* (*KYP*) |
| supp#3 SUP23 | G->A | splicing | Chr5: 4502582 | *KRYPTONITE* (*KYP*) |
| supp#4 SUP66 | G->A | L802F | Chr1: 26248612 | *CHROMETHYLASE3* (*CMT3*) |
| supp#5 SUP33 | G->A | P715L | Chr1: 26249218 | *CHROMETHYLASE3* (*CMT3*) |
| supp#6 SUP59 | C->T | G456D | Chr1: 26250521 | *CHROMETHYLASE3* (*CMT3*) |
| supp#7 SUP6 | G->A | P475L | Chr1: 26250464 | *CHROMETHYLASE3* (*CMT3*) |
| supp#8 SUP78 | G->A | STOP | Chr1: 26250258 | *CHROMETHYLASE3* (*CMT3*) |
| supp#9 SUP151 | G->A | P675L | Chr1: 26249523 | *CHROMETHYLASE3* (*CMT3*) |
| supp#10 SUP137 | G->A | L663F | Chr1: 26249560 | *CHROMETHYLASE3* (*CMT3*) |
| supp#11 | C->T | STOP | Position 586 | *FLOWERING TIME CONTROL PROTEIN* (*FPA*) |
| supp#12 SUP89 | C->T | G144E | Chr3: 4481079 | *LYSINE-SPECIFIC DEMETHYLASE1 LIKE2* (*LDL2*) |
| supp#13 SUP106 | C deletion | STOP | Chr5:1111465 | *GYRB3* |

### Supplementary Table S4

### List of TEs upregulated in *gyrb3* compared to Col-0

Results of DESeq2 differential analyis (RNA-seq). TEs were filtered for differentially abundant entries (log2FC < -1 or >1; p-value < 0.05)

| | | | | overlaping non-CG DMR *gyrb3* /Col | |
|---|---|---|---|---|---|
| **TE id** | **Fold change** | **p value** | **q value** | **CHG** | **CHH** |
| AT1TE22850 | 4.63 | 1.10E-10 | 1.40E-08 | yes | yes |
| AT1TE42875 | 10.85 | 0.00025977 | 0.0066154 | no | no |
| AT1TE58075 | 30.2 | 1.67E-05 | 0.00058013 | yes | yes |
| AT1TE76215 | 19.28 | 0.00010685 | 0.0028149 | yes | yes |
| AT2TE08225 | 18.85 | 3.58E-06 | 0.00016106 | yes | no |
| AT2TE11560 | 2.48 | 0.0036511 | 0.063738 | yes | no |
| AT2TE12115 | 4.69 | 0.0014527 | 0.03083 | yes | no |
| AT2TE19615 | 2.67 | 0.0031359 | 0.058435 | no | no |
| AT2TE27645 | 26.74 | 4.37E-05 | 0.0012826 | yes | yes |
| AT2TE28020 | 6.52 | 6.54E-08 | 4.54E-06 | no | no |
| AT2TE28025 | 8.22 | 3.10E-05 | 0.0010301 | no | no |
| AT2TE28280 | 8.39 | 7.26E-05 | 0.0019821 | yes | no |
| AT2TE29450 | 48.17 | 8.42E-07 | 4.81E-05 | yes | yes |
| AT2TE29460 | 81.99 | 1.06E-08 | 8.12E-07 | yes | yes |
| AT2TE37050 | 143.6 | 3.21E-10 | 3.51E-08 | yes | no |
| AT2TE77015 | 10.77 | 4.66E-06 | 0.00019794 | yes | no |
| AT2TE78210 | 17.46 | 1.81E-07 | 1.15E-05 | yes | yes |
| AT3TE20780 | 13.28 | 1.07E-05 | 0.00039057 | yes | no |
| AT3TE45385 | 100.86 | 4.26E-47 | 3.26E-44 | yes | yes |
| AT3TE51150 | 2.78 | 9.48E-06 | 0.00036208 | yes | no |
| AT3TE60310 | 13.96 | 1.07E-19 | 2.73E-17 | yes | no |
| AT3TE64435 | 3.7 | 0.0019321 | 0.038845 | yes | no |
| AT3TE68090 | 98.16 | 5.23E-09 | 4.44E-07 | yes | yes |
| AT3TE76225 | 34.33 | 2.06E-06 | 9.83E-05 | yes | yes |
| AT3TE90530 | 30.07 | 8.81E-07 | 4.81E-05 | yes | yes |
| AT3TE94195 | 2.09 | 4.24E-05 | 0.0012826 | no | no |
| AT4TE10600 | 23.15 | 6.31E-06 | 0.0002536 | yes | yes |
| AT5TE00480 | 6.6 | 1.13E-13 | 2.16E-11 | yes | yes |
| AT5TE08220 | 105.48 | 4.68E-10 | 4.47E-08 | yes | yes |
| AT5TE23185 | 20.68 | 7.71E-37 | 2.95E-34 | yes | no |
| AT5TE34980 | 150.55 | 2.84E-11 | 4.34E-09 | yes | yes |
| AT5TE37665 | 18.58 | 5.26E-05 | 0.0014884 | no | no |
| AT5TE47200 | 3.41 | 0.0020025 | 0.039229 | yes | no |
| AT5TE50260 | 24.47 | 3.91E-05 | 0.0012435 | no | no |
| AT5TE61735 | 8.71 | 0.00059104 | 0.014566 | no | no |
| AT5TE69650 | 44.95 | 1.11E-06 | 5.65E-05 | yes | yes |

**Supplementary Table S5**

**List of differently expressed genes**

Results of DESeq2 differential analyis (RNA-seq). Genes were filtered for differentially abundant entries (log2FC < -1 or >1; p-value < 0.05)

Col-0 versus *gyrb3-2*

| gene_id | log2FoldChange | pvalue | padj |
|---|---|---|---|
| AT1G02400 | 1.432916 | 5.39E-10 | 1.39E-07 |
| AT1G04310 | 1.048922 | 1.83E-10 | 5.22E-08 |
| AT1G06080 | 1.553642 | 3.01E-11 | 1.09E-08 |
| AT1G07050 | 1.457159 | 1.26E-06 | 9.39E-05 |
| AT1G08630 | 1.424257 | 1.61E-07 | 1.72E-05 |
| AT1G10060 | 1.218144 | 7.10E-17 | 1.04E-13 |
| AT1G10070 | 1.760104 | 4.93E-13 | 3.12E-10 |
| AT1G10370 | -1.567303 | 7.00E-16 | 7.00E-13 |
| AT1G11185 | 1.012734 | 3.94E-05 | 1.56E-03 |
| AT1G13650 | -2.249509 | 2.34E-13 | 1.66E-10 |
| AT1G14250 | -1.564449 | 3.06E-12 | 1.46E-09 |
| AT1G14930 | 1.379224 | 4.49E-07 | 4.13E-05 |
| AT1G17665 | 1.333929 | 8.00E-11 | 2.50E-08 |
| AT1G19050 | -1.129348 | 4.62E-07 | 4.20E-05 |
| AT1G19530 | 1.152451 | 2.89E-07 | 2.83E-05 |
| AT1G19960 | -1.255783 | 1.35E-07 | 1.48E-05 |
| AT1G23060 | 1.101959 | 1.71E-10 | 4.94E-08 |
| AT1G26761 | -1.306331 | 3.46E-13 | 2.28E-10 |
| AT1G29450 | -1.00005 | 2.69E-08 | 3.96E-06 |
| AT1G55390 | 1.359733 | 8.80E-06 | 4.70E-04 |
| AT1G64660 | 1.226913 | 1.54E-15 | 1.34E-12 |
| AT1G65490 | -1.418355 | 5.35E-07 | 4.62E-05 |
| AT1G67105 | 1.859402 | 3.05E-15 | 2.53E-12 |
| AT1G68050 | 1.265445 | 3.02E-17 | 4.81E-14 |
| AT1G69490 | 1.073042 | 2.13E-09 | 4.74E-07 |
| AT1G73600 | -1.401148 | 2.23E-09 | 4.90E-07 |
| AT1G73870 | -1.187172 | 1.76E-09 | 4.00E-07 |
| AT1G80160 | 1.049682 | 2.73E-05 | 1.18E-03 |
| AT2G04790 | -1.040704 | 1.50E-06 | 1.09E-04 |
| AT2G05914 | 3.215292 | 1.19E-06 | 8.93E-05 |
| AT2G18969 | 1.066371 | 4.94E-07 | 4.36E-05 |
| AT2G21650 | -1.524602 | 1.42E-07 | 1.54E-05 |
| AT2G21910 | 3.386289 | 1.20E-16 | 1.44E-13 |
| AT2G34620 | -1.116018 | 2.43E-12 | 1.19E-09 |
| AT2G34655 | 1.408131 | 4.11E-08 | 5.61E-06 |
| AT2G39250 | -1.19034 | 2.57E-08 | 3.84E-06 |
| AT2G40080 | 1.203072 | 4.05E-07 | 3.78E-05 |
| AT2G40340 | 2.092641 | 3.20E-23 | 8.74E-20 |
| AT2G40955 | 5.940354 | 2.05E-19 | 3.57E-16 |
| AT2G41230 | 2.038838 | 8.01E-13 | 4.25E-10 |
| AT2G41240 | -1.580441 | 1.01E-06 | 7.87E-05 |
| AT2G42690 | -1.021526 | 1.30E-11 | 5.09E-09 |
| AT2G42975 | -1.041583 | 5.80E-10 | 1.48E-07 |
| AT2G44910 | -1.599236 | 7.92E-09 | 1.40E-06 |
| AT2G46830 | -1.397107 | 4.40E-10 | 1.17E-07 |
| AT2G47000 | 1.266131 | 3.44E-08 | 4.83E-06 |
| AT3G01060 | -1.002148 | 8.22E-10 | 1.94E-07 |
| AT3G01970 | 1.197538 | 3.27E-08 | 4.63E-06 |
| AT3G04070 | 1.053648 | 3.63E-08 | 5.02E-06 |
| AT3G15440 | 1.224709 | 6.98E-08 | 8.67E-06 |
| AT3G21320 | 1.39983 | 2.84E-08 | 4.14E-06 |
| AT3G21670 | -1.009262 | 8.05E-10 | 1.93E-07 |
| AT3G22640 | 1.231356 | 4.25E-05 | 1.64E-03 |
| AT3G23150 | 1.128387 | 3.28E-12 | 1.53E-09 |
| AT3G23240 | 3.155869 | 6.47E-10 | 1.63E-07 |
| AT3G29644 | -1.772855 | 1.02E-16 | 1.29E-13 |
| AT3G47430 | -1.00733 | 1.78E-06 | 1.27E-04 |
| AT3G50120 | 1.101799 | 9.14E-06 | 4.85E-04 |
| AT3G51400 | 1.118629 | 2.68E-13 | 1.83E-10 |
| AT3G52720 | -1.051503 | 2.16E-07 | 2.20E-05 |
| AT3G53830 | -1.067026 | 1.17E-07 | 1.33E-05 |
| AT3G54730 | 2.876776 | 4.43E-07 | 4.09E-05 |
| AT3G56080 | -1.2153 | 2.43E-09 | 5.28E-07 |
| AT3G60390 | -1.04723 | 7.18E-08 | 8.85E-06 |
| AT3G62090 | 1.513731 | 2.59E-23 | 8.25E-20 |
| AT4G04223 | 1.635474 | 1.06E-20 | 2.26E-17 |
| AT4G04840 | -1.069425 | 5.19E-09 | 9.92E-07 |
| AT4G08770 | 1.031833 | 1.37E-05 | 6.72E-04 |
| AT4G08870 | -1.372343 | 6.79E-15 | 5.41E-12 |
| AT4G09350 | -1.008823 | 2.97E-06 | 1.89E-04 |
| AT4G13575 | -1.158764 | 1.26E-07 | 1.40E-05 |
| AT4G13770 | -1.02169 | 2.11E-09 | 4.74E-07 |
| AT4G14130 | 1.102261 | 4.89E-07 | 4.36E-05 |
| AT4G15550 | 1.027968 | 1.17E-11 | 4.65E-09 |
| AT4G16857 | -2.184827 | 8.06E-17 | 1.10E-13 |
| AT4G17470 | -1.34604 | 2.83E-09 | 6.07E-07 |
| AT4G19120 | 1.066256 | 4.34E-08 | 5.84E-06 |
| AT4G19430 | 1.182778 | 9.51E-06 | 4.99E-04 |
| AT4G22880 | -1.211809 | 3.59E-05 | 1.44E-03 |
| AT4G27970 | 1.039549 | 7.35E-06 | 4.05E-04 |
| AT4G30290 | 1.139105 | 6.41E-11 | 2.12E-08 |
| AT4G32810 | 1.156355 | 4.85E-08 | 6.34E-06 |
| AT4G33980 | 1.599811 | 1.01E-31 | 1.07E-27 |
| AT4G38410 | 1.277825 | 2.48E-07 | 2.47E-05 |
| AT4G39100 | -1.080704 | 1.43E-20 | 2.74E-17 |
| AT5G01015 | -1.156988 | 2.17E-07 | 2.20E-05 |
| AT5G01330 | 1.621394 | 9.00E-08 | 1.07E-05 |
| AT5G01790 | 1.100351 | 3.05E-09 | 6.47E-07 |
| AT5G02170 | 1.441024 | 1.21E-08 | 2.01E-06 |
| AT5G02540 | 1.074354 | 1.31E-10 | 3.92E-08 |
| AT5G02865 | -1.132867 | 5.11E-07 | 4.46E-05 |

Col-0 versus *ibm2-4 gyrb3-2*

| gene_id | log2FoldChange | pvalue | padj |
|---|---|---|---|
| AT1G01060 | -1.334971 | 2.14E-11 | 1.54E-09 |
| AT1G01600 | -1.033062 | 3.18E-07 | 8.77E-06 |
| AT1G01900 | -1.099201 | 1.04E-08 | 4.20E-07 |
| AT1G02400 | 1.884682 | 1.65E-14 | 2.04E-12 |
| AT1G02470 | 1.332175 | 2.09E-05 | 3.35E-04 |
| AT1G02820 | -1.228788 | 1.35E-05 | 2.28E-04 |
| AT1G02930 | 1.555438 | 3.61E-09 | 1.59E-07 |
| AT1G03055 | -1.225155 | 9.66E-05 | 1.25E-03 |
| AT1G03090 | 1.138732 | 1.64E-09 | 7.90E-08 |
| AT1G03630 | -1.315775 | 9.80E-14 | 1.04E-11 |
| AT1G04133 | 1.074095 | 4.71E-04 | 4.69E-03 |
| AT1G04180 | -1.04055 | 1.22E-04 | 1.52E-03 |
| AT1G04310 | 1.343528 | 4.72E-15 | 6.64E-13 |
| AT1G04350 | -1.030602 | 1.84E-18 | 4.52E-16 |
| AT1G04530 | -1.112413 | 1.41E-22 | 6.75E-20 |
| AT1G04640 | -1.472465 | 1.12E-22 | 5.73E-20 |
| AT1G05397 | 1.008518 | 4.61E-04 | 4.62E-03 |
| AT1G05680 | 1.111089 | 2.89E-05 | 4.42E-04 |
| AT1G06040 | -1.02898 | 9.04E-29 | 7.13E-26 |
| AT1G06080 | 1.3397 | 1.46E-08 | 5.66E-07 |
| AT1G06100 | -1.571528 | 1.14E-06 | 2.69E-05 |
| AT1G06360 | -1.259614 | 7.70E-15 | 1.06E-12 |
| AT1G07040 | 1.507511 | 5.06E-19 | 1.38E-16 |
| AT1G07050 | 2.225113 | 4.91E-10 | 2.73E-08 |
| AT1G08630 | 3.10176 | 5.25E-21 | 1.90E-18 |
| AT1G09080 | -1.540383 | 5.11E-06 | 9.93E-05 |
| AT1G09240 | 1.038585 | 7.68E-08 | 2.49E-06 |
| AT1G09420 | 1.50167 | 4.56E-15 | 6.50E-13 |
| AT1G10060 | 1.462726 | 1.27E-22 | 6.25E-20 |
| AT1G10070 | 3.3707 | 1.35E-36 | 2.48E-33 |
| AT1G10140 | 1.060606 | 1.03E-08 | 4.16E-07 |
| AT1G10370 | -1.779625 | 9.59E-19 | 2.52E-16 |
| AT1G10470 | -1.569184 | 1.29E-20 | 4.25E-18 |
| AT1G10657 | -1.024893 | 7.20E-06 | 1.33E-04 |
| AT1G10960 | -1.014212 | 2.07E-18 | 5.03E-16 |
| AT1G11185 | 1.16981 | 4.27E-05 | 6.17E-04 |
| AT1G11785 | 1.863769 | 5.25E-05 | 7.35E-04 |
| AT1G12080 | 1.481417 | 1.02E-10 | 6.50E-09 |
| AT1G13650 | -2.696088 | 1.26E-16 | 2.35E-14 |
| AT1G13740 | -1.013471 | 1.11E-06 | 2.62E-05 |
| AT1G14250 | -1.697477 | 2.67E-13 | 2.69E-11 |
| AT1G14345 | -1.490777 | 2.39E-17 | 4.84E-15 |
| AT1G14430 | -1.23084 | 3.59E-08 | 1.26E-06 |
| AT1G14600 | 1.135677 | 1.08E-08 | 4.32E-07 |
| AT1G14700 | -1.242046 | 7.25E-13 | 6.81E-11 |
| AT1G14930 | 2.677712 | 8.82E-17 | 1.66E-14 |
| AT1G15040 | 1.753859 | 7.51E-08 | 2.44E-06 |
| AT1G15050 | 1.424125 | 2.93E-05 | 4.49E-04 |
| AT1G15330 | 1.015956 | 1.70E-04 | 2.02E-03 |
| AT1G16060 | -1.835354 | 5.46E-15 | 7.58E-13 |
| AT1G16410 | -1.834951 | 3.33E-08 | 1.19E-06 |
| AT1G16489 | 2.269861 | 1.17E-18 | 3.03E-16 |
| AT1G17030 | -1.00299 | 2.67E-04 | 2.95E-03 |
| AT1G17050 | -1.133303 | 2.61E-12 | 2.27E-10 |
| AT1G17665 | 1.763159 | 2.75E-16 | 4.90E-14 |
| AT1G18010 | -5.134355 | 5.45E-08 | 1.83E-06 |
| AT1G18990 | -1.287407 | 8.70E-05 | 1.14E-03 |
| AT1G19050 | -2.045127 | 4.33E-15 | 6.29E-13 |
| AT1G19150 | -1.331923 | 3.59E-21 | 1.34E-18 |
| AT1G19530 | 1.612137 | 6.17E-11 | 4.06E-09 |
| AT1G19540 | 1.480083 | 3.04E-14 | 3.55E-12 |
| AT1G19640 | -1.138087 | 4.51E-04 | 4.52E-03 |
| AT1G19960 | -1.817574 | 1.84E-11 | 1.35E-09 |
| AT1G20515 | 1.050201 | 2.32E-04 | 2.63E-03 |
| AT1G20900 | 1.073299 | 2.67E-08 | 9.73E-07 |
| AT1G21400 | 1.653604 | 1.25E-18 | 3.18E-16 |
| AT1G21870 | -1.308848 | 1.11E-04 | 1.40E-03 |
| AT1G22590 | -1.019471 | 5.39E-06 | 1.04E-04 |
| AT1G22630 | -1.156705 | 2.31E-10 | 1.38E-08 |
| AT1G23965 | 1.139988 | 9.17E-05 | 1.20E-03 |
| AT1G24148 | -1.252497 | 1.56E-06 | 3.52E-05 |
| AT1G25141 | -4.065032 | 6.58E-07 | 1.67E-05 |
| AT1G26558 | 2.106323 | 2.01E-05 | 3.24E-04 |
| AT1G26560 | -1.319614 | 1.20E-08 | 4.75E-07 |
| AT1G26665 | 1.149429 | 2.16E-09 | 1.01E-07 |
| AT1G26761 | -2.374787 | 5.35E-31 | 6.57E-28 |
| AT1G27020 | 1.359025 | 1.64E-14 | 2.04E-12 |
| AT1G27670 | 1.149813 | 3.96E-05 | 5.77E-04 |
| AT1G28050 | 1.1085 | 2.15E-19 | 6.24E-17 |
| AT1G29430 | -1.731403 | 1.70E-15 | 2.64E-13 |
| AT1G29440 | -1.61952 | 1.49E-14 | 1.91E-12 |
| AT1G29450 | -1.622027 | 1.64E-16 | 3.02E-14 |
| AT1G29460 | -1.12193 | 1.74E-10 | 1.07E-08 |
| AT1G29500 | -1.107954 | 7.32E-09 | 3.02E-07 |
| AT1G29510 | -1.203435 | 4.35E-12 | 3.64E-10 |
| AT1G30080 | 1.135822 | 2.31E-07 | 6.65E-06 |
| AT1G30160 | 1.08198 | 8.29E-06 | 1.50E-04 |
| AT1G30290 | -3.862774 | 3.75E-37 | 7.52E-34 |
| AT1G30720 | 1.116034 | 1.88E-06 | 4.15E-05 |
| AT1G31580 | -1.571711 | 5.97E-14 | 6.66E-12 |
| AT1G31835 | 1.005541 | 5.85E-05 | 8.08E-04 |

Col-0 versus *ibm2-4*

| gene_id | log2FoldChange | pvalue | padj |
|---|---|---|---|
| AT1G02920 | 2.177048 | 4.17E-16 | 1.59E-13 |
| AT1G02930 | 2.269244 | 1.65E-15 | 5.56E-13 |
| AT1G03820 | 1.491296 | 1.53E-08 | 1.13E-06 |
| AT1G04090 | 1.118417 | 1.11E-04 | 2.09E-03 |
| AT1G04640 | -1.03047 | 9.07E-13 | 1.73E-10 |
| AT1G05300 | 1.034979 | 1.21E-06 | 5.01E-05 |
| AT1G05397 | 1.35153 | 2.74E-04 | 4.24E-03 |
| AT1G05675 | 1.267499 | 1.11E-04 | 2.08E-03 |
| AT1G05680 | 2.21802 | 1.07E-12 | 2.02E-10 |
| AT1G10070 | 1.797613 | 1.64E-12 | 2.85E-10 |
| AT1G10370 | -1.011488 | 3.11E-08 | 2.07E-06 |
| AT1G11210 | 1.070259 | 1.65E-17 | 7.92E-15 |
| AT1G11410 | -1.118214 | 6.06E-09 | 5.01E-07 |
| AT1G13470 | 1.024398 | 3.14E-05 | 7.59E-04 |
| AT1G13650 | -1.670443 | 1.71E-08 | 1.24E-06 |
| AT1G14780 | 1.076652 | 1.06E-05 | 3.09E-04 |
| AT1G14930 | 1.14014 | 2.64E-05 | 6.57E-04 |
| AT1G15125 | 1.305323 | 1.52E-17 | 7.62E-15 |
| AT1G16489 | 2.326261 | 6.14E-19 | 3.65E-16 |
| AT1G17180 | 1.255488 | 2.24E-05 | 5.73E-04 |
| AT1G17250 | 1.047015 | 1.00E-04 | 1.93E-03 |
| AT1G17960 | 1.302791 | 2.56E-06 | 9.44E-05 |
| AT1G18590 | 1.225552 | 4.09E-14 | 1.04E-11 |
| AT1G19180 | 1.017044 | 5.20E-09 | 4.39E-07 |
| AT1G19610 | 2.116321 | 1.24E-08 | 9.40E-07 |
| AT1G20515 | 1.491259 | 6.69E-06 | 2.09E-04 |
| AT1G21651 | -1.480784 | 1.96E-13 | 4.35E-11 |
| AT1G21870 | -1.1136 | 2.71E-04 | 4.20E-03 |
| AT1G23100 | 1.055835 | 3.34E-06 | 1.18E-04 |
| AT1G24095 | 1.185275 | 2.08E-06 | 7.99E-05 |
| AT1G24140 | 1.025425 | 1.94E-04 | 3.23E-03 |
| AT1G24260 | 1.105985 | 2.36E-09 | 2.19E-07 |
| AT1G25141 | -3.788045 | 2.78E-07 | 1.41E-05 |
| AT1G26380 | 2.399401 | 7.71E-11 | 1.02E-08 |
| AT1G26390 | 1.569882 | 1.84E-05 | 4.88E-04 |
| AT1G26410 | 1.552304 | 5.73E-05 | 1.23E-03 |
| AT1G26420 | 2.203933 | 7.20E-12 | 1.15E-09 |
| AT1G26761 | -1.166947 | 8.91E-11 | 1.15E-08 |
| AT1G27020 | 1.181994 | 1.31E-11 | 1.98E-09 |
| AT1G28370 | 1.040104 | 2.29E-04 | 3.66E-03 |
| AT1G30290 | -3.636843 | 3.66E-37 | 9.18E-34 |
| AT1G30757 | 1.035651 | 8.22E-06 | 2.48E-04 |
| AT1G30760 | 1.514674 | 1.20E-09 | 1.22E-07 |
| AT1G31835 | 1.273113 | 1.68E-06 | 6.64E-05 |
| AT1G32870 | 1.177437 | 4.31E-17 | 1.80E-14 |
| AT1G32960 | 1.196326 | 2.52E-04 | 3.97E-03 |
| AT1G35240 | -2.79196 | 2.26E-05 | 5.78E-04 |
| AT1G35320 | 1.41427 | 4.60E-14 | 1.15E-11 |
| AT1G35730 | 1.715283 | 8.70E-05 | 1.72E-03 |
| AT1G36070 | -1.623197 | 9.19E-25 | 1.48E-21 |
| AT1G37150 | 1.19485 | 1.51E-04 | 2.67E-03 |
| AT1G43770 | -1.423892 | 5.71E-12 | 9.21E-10 |
| AT1G49000 | 1.369615 | 1.74E-05 | 4.66E-04 |
| AT1G49570 | 1.233526 | 1.23E-04 | 2.25E-03 |
| AT1G49860 | 1.159487 | 5.00E-06 | 1.66E-04 |
| AT1G51270 | 1.621368 | 3.08E-11 | 4.41E-09 |
| AT1G51800 | 1.122632 | 3.08E-05 | 7.47E-04 |
| AT1G52070 | 1.159984 | 3.10E-05 | 7.49E-04 |
| AT1G54000 | 1.203835 | 1.34E-09 | 1.33E-07 |
| AT1G54020 | 1.319822 | 5.55E-05 | 1.20E-03 |
| AT1G54820 | -1.015154 | 1.15E-09 | 1.17E-07 |
| AT1G57980 | -1.448581 | 7.24E-11 | 9.67E-09 |
| AT1G58400 | 1.404159 | 4.82E-08 | 3.07E-06 |
| AT1G58602 | -2.307229 | 1.45E-21 | 1.21E-18 |
| AT1G59030 | -1.591185 | 7.33E-05 | 1.51E-03 |
| AT1G60095 | 1.692057 | 5.22E-05 | 1.15E-03 |
| AT1G60110 | 1.633708 | 2.07E-08 | 1.45E-06 |
| AT1G61120 | 1.425237 | 2.71E-04 | 4.21E-03 |
| AT1G61500 | -2.122319 | 1.18E-18 | 6.81E-16 |
| AT1G61800 | 1.792109 | 3.15E-06 | 1.12E-04 |
| AT1G62290 | 1.189164 | 5.18E-16 | 1.93E-13 |
| AT1G63240 | -1.30171 | 2.17E-23 | 2.17E-20 |
| AT1G64220 | 1.042241 | 3.47E-04 | 5.15E-03 |
| AT1G64590 | -1.2632 | 6.94E-05 | 1.45E-03 |
| AT1G65370 | 1.01834 | 2.11E-14 | 5.67E-12 |
| AT1G65500 | 2.385558 | 2.39E-10 | 2.79E-08 |
| AT1G66700 | 2.664743 | 4.95E-08 | 3.15E-06 |
| AT1G66920 | 2.124424 | 2.33E-20 | 1.64E-17 |
| AT1G67105 | 5.097802 | 3.83E-99 | 8.65E-95 |
| AT1G68290 | 1.450074 | 2.41E-04 | 3.83E-03 |
| AT1G69880 | 2.798048 | 8.13E-16 | 2.91E-13 |
| AT1G71380 | 3.998725 | 8.76E-08 | 5.26E-06 |
| AT1G71770 | 1.669658 | 1.67E-05 | 4.51E-04 |
| AT1G74640 | -1.063351 | 2.58E-24 | 3.43E-21 |
| AT1G75040 | 1.104787 | 1.04E-07 | 6.02E-06 |
| AT1G77120 | 1.138115 | 6.50E-11 | 8.79E-09 |
| AT1G78410 | 1.496107 | 3.06E-05 | 7.42E-04 |
| AT1G78950 | -1.489396 | 6.65E-06 | 2.08E-04 |
| AT1G79680 | 1.298196 | 7.04E-05 | 1.47E-03 |
| AT1G79890 | -1.304551 | 7.84E-05 | 1.59E-03 |
| AT1G80130 | 1.128685 | 1.21E-09 | 1.22E-07 |

| | | | |
|---|---|---|---|
| AT5G04110 | -2.860022 | 1.75E-29 | 1.11E-25 |
| AT5G10250 | -1.10935 | 2.54E-06 | 1.68E-04 |
| AT5G17860 | 1.360554 | 1.61E-08 | 2.49E-06 |
| AT5G18470 | 1.064257 | 1.64E-07 | 1.74E-05 |
| AT5G22390 | -1.289104 | 3.11E-08 | 4.43E-06 |
| AT5G22500 | 1.669074 | 7.32E-16 | 7.00E-13 |
| AT5G23240 | 1.610589 | 2.84E-16 | 3.20E-13 |
| AT5G24120 | -1.0444 | 5.87E-09 | 1.06E-06 |
| AT5G24240 | 1.375587 | 7.97E-06 | 4.32E-04 |
| AT5G26270 | 2.029586 | 4.41E-23 | 1.05E-19 |
| AT5G38005 | 1.504415 | 1.49E-08 | 2.38E-06 |
| AT5G38430 | -1.013225 | 1.00E-10 | 3.08E-08 |
| AT5G39610 | 1.006874 | 4.25E-12 | 1.93E-09 |
| AT5G42900 | 2.379247 | 1.12E-31 | 1.07E-27 |
| AT5G45650 | -1.164451 | 4.79E-12 | 2.13E-09 |
| AT5G47590 | 1.473216 | 2.15E-11 | 8.06E-09 |
| AT5G48250 | 1.055776 | 8.54E-16 | 7.77E-13 |
| AT5G48490 | -1.4651 | 1.80E-07 | 1.87E-05 |
| AT5G49690 | 1.319469 | 8.90E-09 | 1.52E-06 |
| AT5G50950 | -1.42732 | 1.43E-23 | 5.45E-20 |
| AT5G57640 | 1.034671 | 1.10E-05 | 5.66E-04 |
| AT5G59130 | -1.182791 | 2.65E-08 | 3.93E-06 |
| AT5G60100 | 1.360916 | 8.95E-08 | 1.07E-05 |
| AT5G62920 | -1.35776 | 1.42E-06 | 1.05E-04 |
| AT5G63650 | 1.95831 | 1.70E-24 | 8.13E-21 |

| | | | |
|---|---|---|---|
| AT1G32900 | -1.217009 | 1.41E-10 | 8.83E-09 |
| AT1G35320 | 1.178874 | 1.85E-10 | 1.12E-08 |
| AT1G35560 | -1.069562 | 6.53E-07 | 1.66E-05 |
| AT1G35730 | 1.527224 | 1.07E-04 | 1.35E-03 |
| AT1G36070 | -1.670005 | 1.79E-26 | 1.24E-23 |
| AT1G37130 | -1.185214 | 2.57E-06 | 5.48E-05 |
| AT1G37150 | 1.067915 | 3.78E-04 | 3.91E-03 |
| AT1G38065 | -14.64803 | 2.48E-07 | 7.03E-06 |
| AT1G44000 | -1.245667 | 5.32E-15 | 7.44E-13 |
| AT1G44446 | -1.597233 | 7.42E-17 | 1.44E-14 |
| AT1G45201 | -1.183868 | 2.23E-14 | 2.68E-12 |
| AT1G48260 | -1.323199 | 1.42E-06 | 3.26E-05 |
| AT1G50320 | -1.033035 | 2.48E-21 | 9.78E-19 |
| AT1G51270 | 1.51985 | 1.51E-10 | 9.39E-09 |
| AT1G52290 | -1.286814 | 1.34E-08 | 5.23E-07 |
| AT1G52410 | -1.018646 | 1.70E-09 | 8.11E-08 |
| AT1G52890 | 1.628593 | 2.96E-10 | 1.73E-08 |
| AT1G54820 | -1.328695 | 3.94E-14 | 4.50E-12 |
| AT1G55180 | -1.48052 | 1.31E-05 | 2.23E-04 |
| AT1G55390 | 2.074802 | 7.65E-08 | 2.48E-06 |
| AT1G55810 | 1.254939 | 3.20E-10 | 1.86E-08 |
| AT1G57980 | -1.117065 | 1.31E-07 | 3.98E-06 |
| AT1G58280 | -1.037005 | 1.17E-09 | 5.92E-08 |
| AT1G58400 | 1.416689 | 2.57E-08 | 9.39E-07 |
| AT1G58602 | -2.227952 | 7.10E-21 | 2.52E-18 |
| AT1G60110 | 1.772174 | 5.92E-10 | 3.19E-08 |
| AT1G61300 | -1.318549 | 4.51E-09 | 1.94E-07 |
| AT1G61500 | -1.311444 | 6.95E-10 | 3.71E-08 |
| AT1G61795 | -1.021048 | 5.65E-05 | 7.85E-04 |
| AT1G62085 | -1.540146 | 3.19E-16 | 5.59E-14 |
| AT1G62250 | -1.191833 | 1.22E-18 | 3.12E-16 |
| AT1G62480 | 1.193548 | 1.51E-08 | 5.84E-07 |
| AT1G62560 | -1.802965 | 1.55E-07 | 4.64E-06 |
| AT1G63240 | -1.289634 | 5.77E-23 | 3.03E-20 |
| AT1G63710 | -2.477008 | 4.96E-10 | 2.73E-08 |
| AT1G64160 | 4.040069 | 5.44E-07 | 1.41E-05 |
| AT1G64220 | 1.419876 | 3.84E-05 | 5.61E-04 |
| AT1G64500 | -1.12789 | 4.00E-10 | 2.27E-08 |
| AT1G64660 | 1.521713 | 6.86E-22 | 2.91E-19 |
| AT1G64770 | -1.187227 | 8.24E-25 | 5.06E-22 |
| AT1G64860 | -1.207095 | 2.58E-16 | 4.62E-14 |
| AT1G65390 | 1.189269 | 3.51E-05 | 5.23E-04 |
| AT1G65450 | -1.017891 | 8.30E-08 | 2.67E-06 |
| AT1G65490 | -1.5357 | 4.57E-07 | 1.21E-05 |
| AT1G65860 | -1.530063 | 7.75E-07 | 1.93E-05 |
| AT1G66040 | 1.320232 | 2.71E-07 | 7.59E-06 |
| AT1G66130 | -1.151171 | 6.70E-14 | 7.33E-12 |
| AT1G66920 | 1.287894 | 3.93E-09 | 1.72E-07 |
| AT1G67105 | 3.624647 | 4.25E-50 | 3.13E-46 |
| AT1G68050 | 1.502062 | 1.21E-22 | 6.08E-20 |
| AT1G68725 | -1.361558 | 1.39E-05 | 2.35E-04 |
| AT1G69160 | -1.165958 | 3.27E-21 | 1.25E-18 |
| AT1G69490 | 1.207772 | 1.06E-10 | 6.75E-09 |
| AT1G69523 | -1.161665 | 2.51E-09 | 1.15E-07 |
| AT1G69570 | -1.083881 | 8.39E-14 | 8.99E-12 |
| AT1G69730 | -1.262369 | 5.20E-07 | 1.35E-05 |
| AT1G69870 | 1.12442 | 3.22E-11 | 2.23E-09 |
| AT1G72330 | 1.046396 | 1.51E-14 | 1.93E-12 |
| AT1G72416 | -1.501239 | 9.48E-07 | 2.31E-05 |
| AT1G73550 | -1.110967 | 1.10E-04 | 1.39E-03 |
| AT1G73600 | -1.43442 | 4.07E-09 | 1.78E-07 |
| AT1G73870 | -1.20224 | 4.14E-09 | 1.80E-07 |
| AT1G74070 | -1.145227 | 2.71E-10 | 1.60E-08 |
| AT1G74110 | 1.262804 | 2.83E-04 | 3.10E-03 |
| AT1G74640 | -1.359781 | 1.49E-36 | 2.53E-33 |
| AT1G75100 | -1.238684 | 3.00E-20 | 8.94E-18 |
| AT1G75280 | -1.202061 | 3.43E-16 | 5.97E-14 |
| AT1G76100 | -1.022155 | 1.32E-14 | 1.73E-12 |
| AT1G76110 | -1.215567 | 3.10E-22 | 1.46E-19 |
| AT1G76410 | 1.048933 | 6.11E-09 | 2.55E-07 |
| AT1G77380 | 1.721073 | 2.96E-12 | 2.53E-10 |
| AT1G78370 | -1.948063 | 4.10E-10 | 2.32E-08 |
| AT1G78440 | -1.033721 | 3.94E-04 | 4.05E-03 |
| AT1G78950 | -1.086708 | 1.28E-04 | 1.59E-03 |
| AT1G79360 | 1.299803 | 5.74E-12 | 4.70E-10 |
| AT1G79680 | 1.011087 | 4.81E-04 | 4.77E-03 |
| AT1G80160 | 2.377779 | 1.94E-12 | 1.71E-10 |
| AT1G80730 | -1.162521 | 1.84E-04 | 2.16E-03 |
| AT2G01080 | 1.045486 | 2.35E-14 | 2.79E-12 |
| AT2G01200 | 1.27337 | 1.42E-08 | 5.52E-07 |
| AT2G01422 | 1.302209 | 2.93E-04 | 3.18E-03 |
| AT2G01590 | -1.197195 | 2.65E-15 | 3.98E-13 |
| AT2G02930 | 1.364127 | 2.58E-07 | 7.30E-06 |
| AT2G03130 | 2.898879 | 8.98E-06 | 1.61E-04 |
| AT2G03750 | -1.007804 | 1.41E-09 | 6.96E-08 |
| AT2G04032 | -1.57463 | 4.57E-10 | 2.56E-08 |
| AT2G04039 | -1.028684 | 1.44E-13 | 1.48E-11 |
| AT2G04050 | 2.95384 | 2.38E-11 | 1.70E-09 |
| AT2G05070 | -1.580655 | 9.07E-15 | 1.22E-12 |
| AT2G05100 | -1.534729 | 4.42E-18 | 9.97E-16 |
| AT2G05540 | 1.055982 | 3.37E-06 | 6.88E-05 |
| AT2G05635 | 1.496031 | 1.22E-04 | 1.52E-03 |
| AT2G05825 | -2.652432 | 1.33E-05 | 2.25E-04 |
| AT2G05914 | 3.21366 | 1.02E-06 | 2.45E-05 |
| AT2G06002 | 1.506393 | 9.62E-06 | 1.71E-04 |
| AT2G14170 | 1.274882 | 2.03E-14 | 2.45E-12 |
| AT2G15555 | 3.632999 | 2.04E-06 | 4.46E-05 |
| AT2G15830 | 1.030728 | 2.35E-07 | 6.74E-06 |
| AT2G15880 | 1.264995 | 3.64E-07 | 9.90E-06 |

| | | | |
|---|---|---|---|
| AT1G80240 | 1.408009 | 5.48E-06 | 1.78E-04 |
| AT1G80520 | 1.200298 | 7.42E-05 | 1.52E-03 |
| AT1G80570 | -1.711212 | 9.22E-16 | 3.25E-13 |
| AT2G01200 | 1.132342 | 2.18E-07 | 1.14E-05 |
| AT2G01422 | 1.701346 | 1.38E-04 | 2.48E-03 |
| AT2G02930 | 1.363107 | 3.29E-07 | 1.64E-05 |
| AT2G03130 | 4.695134 | 2.58E-07 | 1.32E-05 |
| AT2G04032 | -1.534559 | 1.77E-09 | 1.71E-07 |
| AT2G04040 | 1.653437 | 1.14E-06 | 4.72E-05 |
| AT2G04050 | 3.584207 | 5.19E-15 | 1.60E-12 |
| AT2G05825 | -3.215943 | 1.09E-05 | 3.15E-04 |
| AT2G05914 | 4.464361 | 2.54E-09 | 2.30E-07 |
| AT2G07708 | 1.222183 | 3.98E-04 | 5.75E-03 |
| AT2G11810 | -1.011274 | 5.63E-05 | 1.22E-03 |
| AT2G12550 | -1.964795 | 6.81E-23 | 6.41E-20 |
| AT2G15390 | 1.106568 | 8.43E-07 | 3.65E-05 |
| AT2G15400 | 1.462465 | 1.45E-09 | 1.43E-07 |
| AT2G15490 | 1.025034 | 2.19E-06 | 8.36E-05 |
| AT2G16595 | -1.093918 | 2.50E-07 | 1.29E-05 |
| AT2G16660 | -1.045854 | 5.30E-07 | 2.45E-05 |
| AT2G17690 | 3.654318 | 2.55E-12 | 4.30E-10 |
| AT2G18193 | 3.026797 | 1.47E-32 | 2.55E-29 |
| AT2G18660 | 3.264473 | 6.82E-21 | 5.13E-18 |
| AT2G18690 | 1.106438 | 1.29E-05 | 3.62E-04 |
| AT2G18890 | -1.353502 | 5.23E-16 | 1.93E-13 |
| AT2G18969 | 1.159533 | 2.01E-07 | 1.07E-05 |
| AT2G19210 | 3.495558 | 6.49E-06 | 2.04E-04 |
| AT2G19240 | -1.342386 | 1.88E-04 | 3.14E-03 |
| AT2G20750 | -1.165534 | 5.54E-08 | 3.48E-06 |
| AT2G20800 | 4.386812 | 1.40E-10 | 1.74E-08 |
| AT2G21260 | 1.042432 | 3.21E-04 | 4.83E-03 |
| AT2G21640 | 3.256997 | 1.23E-38 | 5.54E-35 |
| AT2G21910 | 3.317141 | 2.80E-16 | 1.09E-13 |
| AT2G22880 | 1.856316 | 2.95E-06 | 1.06E-04 |
| AT2G25470 | 2.60675 | 3.21E-06 | 1.14E-04 |
| AT2G26010 | 2.296546 | 2.95E-07 | 1.48E-05 |
| AT2G26440 | 1.067234 | 1.63E-06 | 6.50E-05 |
| AT2G26560 | 1.019639 | 1.02E-06 | 4.28E-05 |
| AT2G28250 | -1.021763 | 6.81E-08 | 4.18E-06 |
| AT2G28815 | 2.127618 | 2.18E-09 | 2.04E-07 |
| AT2G28880 | -1.127947 | 3.39E-09 | 3.00E-07 |
| AT2G29460 | 1.590309 | 5.80E-07 | 2.64E-05 |
| AT2G30540 | -1.85493 | 5.94E-09 | 4.95E-07 |
| AT2G30750 | 2.381726 | 1.72E-08 | 1.25E-06 |
| AT2G30766 | -1.440663 | 1.06E-08 | 8.20E-07 |
| AT2G30770 | 2.157953 | 1.88E-07 | 1.01E-05 |
| AT2G31230 | 1.048993 | 1.55E-05 | 4.24E-04 |
| AT2G31830 | -1.206977 | 6.18E-05 | 1.32E-03 |
| AT2G31902 | -2.088115 | 3.38E-08 | 2.25E-06 |
| AT2G32020 | 1.021098 | 3.32E-04 | 4.97E-03 |
| AT2G32179 | -2.776146 | 6.34E-06 | 2.00E-04 |
| AT2G32180 | -1.663408 | 6.12E-15 | 1.82E-12 |
| AT2G32280 | 1.095048 | 1.71E-09 | 1.66E-07 |
| AT2G32640 | -1.47764 | 1.56E-09 | 1.53E-07 |
| AT2G32650 | -1.117135 | 1.56E-10 | 1.90E-08 |
| AT2G32810 | -1.58909 | 1.28E-18 | 7.25E-16 |
| AT2G33175 | 1.853891 | 7.03E-07 | 3.11E-05 |
| AT2G33240 | -2.241839 | 8.34E-06 | 2.51E-04 |
| AT2G33710 | 1.598704 | 5.66E-07 | 2.59E-05 |
| AT2G34010 | -1.104034 | 6.50E-05 | 1.37E-03 |
| AT2G34655 | 2.171793 | 5.87E-14 | 1.43E-11 |
| AT2G34790 | 1.263226 | 2.71E-08 | 1.86E-06 |
| AT2G35720 | -1.151922 | 1.09E-12 | 2.03E-10 |
| AT2G35820 | -1.368177 | 4.16E-17 | 1.77E-14 |
| AT2G38530 | 2.185035 | 2.21E-23 | 2.17E-20 |
| AT2G38860 | 1.656473 | 1.65E-17 | 7.92E-15 |
| AT2G39210 | 1.130365 | 2.62E-06 | 9.68E-05 |
| AT2G39250 | -1.329125 | 4.99E-09 | 4.25E-07 |
| AT2G39330 | 1.853092 | 1.31E-06 | 5.37E-05 |
| AT2G39350 | 1.223151 | 1.52E-11 | 2.28E-09 |
| AT2G39370 | 1.676949 | 3.40E-06 | 1.20E-04 |
| AT2G39510 | 1.261383 | 5.70E-06 | 1.83E-04 |
| AT2G40030 | -1.117348 | 3.06E-08 | 2.05E-06 |
| AT2G40340 | 1.242396 | 1.68E-09 | 1.64E-07 |
| AT2G41070 | 1.167916 | 9.00E-11 | 1.15E-08 |
| AT2G41230 | 1.454639 | 2.11E-07 | 1.11E-05 |
| AT2G41240 | -1.599209 | 5.47E-06 | 1.78E-04 |
| AT2G41480 | 1.080362 | 1.78E-04 | 3.02E-03 |
| AT2G41730 | 1.542275 | 2.67E-10 | 3.04E-08 |
| AT2G43140 | 1.328836 | 7.76E-06 | 2.36E-04 |
| AT2G43510 | 1.060899 | 1.44E-05 | 3.96E-04 |
| AT2G43570 | 1.740323 | 1.86E-09 | 1.79E-07 |
| AT2G43620 | 3.915835 | 2.95E-56 | 2.22E-52 |
| AT2G46650 | 1.120632 | 7.23E-12 | 1.15E-09 |
| AT2G46830 | -1.070305 | 8.92E-07 | 3.84E-05 |
| AT2G47520 | 2.818638 | 2.16E-05 | 5.58E-04 |
| AT2G47780 | 1.598468 | 1.26E-08 | 9.52E-07 |
| AT3G01600 | 3.88261 | 2.82E-38 | 1.06E-34 |
| AT3G02840 | 1.315873 | 9.82E-05 | 1.90E-03 |
| AT3G03660 | 2.858004 | 1.54E-10 | 1.89E-08 |
| AT3G04350 | -1.210055 | 3.51E-33 | 6.60E-30 |
| AT3G05660 | 1.237655 | 5.49E-06 | 1.78E-04 |
| AT3G06160 | -1.000128 | 1.03E-05 | 3.00E-04 |
| AT3G06170 | -1.327029 | 5.77E-50 | 3.26E-46 |
| AT3G06435 | 2.053783 | 7.43E-17 | 2.94E-14 |
| AT3G09405 | 1.1075 | 4.74E-06 | 1.60E-04 |
| AT3G11050 | 1.306404 | 1.29E-05 | 3.62E-04 |
| AT3G11110 | -1.333263 | 4.79E-07 | 2.25E-05 |
| AT3G11410 | -1.007325 | 3.48E-08 | 2.29E-06 |

| | | | |
|---|---|---|---|
| AT2G17690 | 2.195279 | 8.06E-07 | 2.00E-05 |
| AT2G17900 | 1.063746 | 5.54E-10 | 3.01E-08 |
| AT2G18050 | 1.231382 | 1.32E-09 | 6.57E-08 |
| AT2G18193 | 1.426034 | 2.24E-09 | 1.04E-07 |
| AT2G18690 | 1.218177 | 4.26E-06 | 8.47E-05 |
| AT2G18969 | 1.047364 | 2.33E-06 | 5.02E-05 |
| AT2G19210 | 2.02479 | 4.33E-05 | 6.25E-04 |
| AT2G19800 | 1.093748 | 1.22E-05 | 2.09E-04 |
| AT2G20750 | -1.150251 | 1.04E-07 | 3.25E-06 |
| AT2G20800 | 2.619936 | 3.09E-06 | 6.40E-05 |
| AT2G21640 | 2.586224 | 1.54E-25 | 1.00E-22 |
| AT2G21650 | -1.466011 | 7.91E-07 | 1.97E-05 |
| AT2G21660 | 1.266082 | 5.89E-12 | 4.78E-10 |
| AT2G21910 | 4.247504 | 9.70E-26 | 6.49E-23 |
| AT2G22190 | -1.357566 | 1.75E-08 | 6.65E-07 |
| AT2G22860 | 1.293206 | 7.17E-06 | 1.32E-04 |
| AT2G22880 | 1.188895 | 1.78E-04 | 2.10E-03 |
| AT2G23000 | -1.466748 | 3.95E-11 | 2.68E-09 |
| AT2G24150 | -1.092659 | 1.16E-15 | 1.85E-13 |
| AT2G24850 | -1.464564 | 8.83E-05 | 1.16E-03 |
| AT2G28815 | 2.151732 | 3.85E-10 | 2.20E-08 |
| AT2G29650 | -1.174808 | 8.42E-16 | 1.36E-13 |
| AT2G30520 | -1.022265 | 1.70E-14 | 2.07E-12 |
| AT2G30766 | -2.635234 | 3.83E-18 | 8.91E-16 |
| AT2G31085 | 1.862472 | 9.41E-10 | 4.84E-08 |
| AT2G31380 | -1.133473 | 4.80E-10 | 2.68E-08 |
| AT2G32180 | -1.553597 | 7.59E-14 | 8.21E-12 |
| AT2G32640 | -1.317138 | 2.87E-08 | 1.04E-06 |
| AT2G32650 | -1.280898 | 8.24E-13 | 7.61E-11 |
| AT2G32810 | -1.665847 | 1.26E-20 | 4.23E-18 |
| AT2G33710 | 1.25296 | 1.98E-05 | 3.20E-04 |
| AT2G34010 | -1.417303 | 5.75E-06 | 1.10E-04 |
| AT2G34140 | 1.242244 | 7.43E-07 | 1.86E-05 |
| AT2G34510 | -1.448252 | 9.22E-12 | 7.22E-10 |
| AT2G34620 | -1.596165 | 1.17E-21 | 4.76E-19 |
| AT2G34655 | 1.370121 | 3.25E-07 | 8.93E-06 |
| AT2G34790 | 1.382707 | 1.73E-09 | 8.27E-08 |
| AT2G35150 | -1.245265 | 7.03E-05 | 9.51E-04 |
| AT2G35260 | -1.194362 | 5.03E-17 | 9.83E-15 |
| AT2G35720 | -1.072512 | 3.33E-11 | 2.30E-09 |
| AT2G35820 | -1.482385 | 2.25E-19 | 6.45E-17 |
| AT2G35830 | -1.085962 | 1.53E-10 | 9.52E-09 |
| AT2G36270 | 1.208581 | 3.45E-07 | 9.44E-06 |
| AT2G37460 | -1.019161 | 1.26E-08 | 4.98E-07 |
| AT2G37870 | -1.076139 | 6.03E-04 | 5.76E-03 |
| AT2G37970 | -1.026473 | 2.99E-08 | 1.07E-06 |
| AT2G38400 | 1.240899 | 1.47E-09 | 7.19E-08 |
| AT2G38465 | 1.042851 | 4.83E-06 | 9.50E-05 |
| AT2G38740 | -1.010083 | 1.15E-14 | 1.52E-12 |
| AT2G38995 | -1.132276 | 8.08E-06 | 1.47E-04 |
| AT2G39250 | -1.107983 | 4.09E-07 | 1.10E-05 |
| AT2G39980 | 1.004954 | 1.34E-14 | 1.74E-12 |
| AT2G40080 | 1.798968 | 1.09E-11 | 8.48E-10 |
| AT2G40340 | 2.780599 | 3.25E-38 | 7.85E-35 |
| AT2G40350 | 1.499722 | 1.24E-04 | 1.54E-03 |
| AT2G40460 | -1.247643 | 6.32E-14 | 6.98E-12 |
| AT2G40955 | 4.145275 | 9.78E-12 | 7.63E-10 |
| AT2G41230 | 2.32105 | 2.49E-15 | 3.77E-13 |
| AT2G41240 | -2.913211 | 3.74E-11 | 2.55E-09 |
| AT2G41650 | 1.531093 | 1.85E-25 | 1.17E-22 |
| AT2G42170 | 1.06869 | 9.52E-14 | 1.01E-11 |
| AT2G42380 | -1.375338 | 1.59E-12 | 1.42E-10 |
| AT2G42690 | -1.265521 | 7.21E-16 | 1.19E-13 |
| AT2G42975 | -1.328624 | 1.43E-13 | 1.47E-11 |
| AT2G43100 | -1.195648 | 2.11E-07 | 6.14E-06 |
| AT2G43140 | 1.357912 | 4.88E-06 | 9.57E-05 |
| AT2G43400 | 1.191947 | 8.07E-11 | 5.25E-09 |
| AT2G43620 | 2.450531 | 2.18E-23 | 1.20E-20 |
| AT2G44230 | -1.024842 | 2.61E-09 | 1.19E-07 |
| AT2G44910 | -1.85599 | 3.27E-10 | 1.89E-08 |
| AT2G45210 | 1.301496 | 5.59E-08 | 1.87E-06 |
| AT2G46420 | -1.100502 | 9.33E-10 | 4.81E-08 |
| AT2G46650 | -1.176794 | 1.47E-09 | 7.18E-08 |
| AT2G46830 | -1.77366 | 1.42E-13 | 1.47E-11 |
| AT2G47000 | 1.526007 | 4.96E-10 | 2.73E-08 |
| AT2G47550 | 1.166258 | 3.40E-05 | 5.09E-04 |
| AT2G47750 | -1.041366 | 2.27E-05 | 3.60E-04 |
| AT3G01060 | -1.348062 | 7.14E-15 | 9.85E-13 |
| AT3G01345 | 3.579689 | 3.44E-08 | 1.22E-06 |
| AT3G01440 | -1.208195 | 1.86E-06 | 4.11E-05 |
| AT3G01550 | -1.209829 | 3.04E-11 | 2.12E-09 |
| AT3G01600 | 3.379979 | 2.82E-29 | 2.49E-26 |
| AT3G01970 | 1.430012 | 5.29E-10 | 2.89E-08 |
| AT3G02020 | -1.342816 | 1.59E-06 | 3.58E-05 |
| AT3G02380 | -1.288497 | 4.59E-12 | 3.81E-10 |
| AT3G02550 | 1.135901 | 7.53E-10 | 3.97E-08 |
| AT3G02870 | -1.013076 | 5.65E-11 | 3.74E-09 |
| AT3G03470 | 1.043898 | 6.92E-12 | 5.58E-10 |
| AT3G03780 | -1.198303 | 4.72E-17 | 9.38E-15 |
| AT3G03820 | -1.153017 | 8.09E-07 | 2.00E-05 |
| AT3G03830 | -1.058052 | 2.16E-08 | 8.05E-07 |
| AT3G03840 | -1.068221 | 8.67E-07 | 2.14E-05 |
| AT3G03850 | -1.150672 | 1.33E-07 | 4.02E-06 |
| AT3G04070 | 1.2554 | 4.96E-10 | 2.73E-08 |
| AT3G04210 | -1.016098 | 9.74E-07 | 2.36E-05 |
| AT3G04290 | -1.004991 | 3.01E-11 | 2.11E-09 |
| AT3G06170 | -1.168438 | 1.04E-39 | 2.87E-36 |
| AT3G06435 | 2.272767 | 1.08E-20 | 3.67E-18 |
| AT3G06850 | 1.15337 | 6.91E-11 | 4.54E-09 |
| AT3G13090 | 1.103321 | 6.90E-05 | 1.44E-03 |
| AT3G13784 | -1.839928 | 4.98E-07 | 2.32E-05 |
| AT3G14450 | -1.000015 | 1.78E-06 | 7.01E-05 |
| AT3G14810 | -1.459134 | 1.37E-12 | 2.47E-10 |
| AT3G15440 | 1.245059 | 1.65E-07 | 9.00E-06 |
| AT3G15518 | 1.754368 | 5.74E-06 | 1.84E-04 |
| AT3G17690 | 1.030523 | 5.42E-04 | 7.33E-03 |
| AT3G20420 | -1.041767 | 2.39E-08 | 1.66E-06 |
| AT3G20470 | 2.019976 | 1.50E-17 | 7.62E-15 |
| AT3G21780 | 1.079429 | 5.37E-08 | 3.39E-06 |
| AT3G21781 | 1.647154 | 5.62E-06 | 1.81E-04 |
| AT3G22160 | 1.595253 | 7.10E-14 | 1.67E-11 |
| AT3G22640 | 3.257686 | 3.14E-07 | 1.57E-05 |
| AT3G23120 | 2.448715 | 1.52E-08 | 1.13E-06 |
| AT3G23240 | 2.359979 | 7.88E-07 | 3.45E-05 |
| AT3G23250 | 2.069177 | 5.99E-06 | 1.91E-04 |
| AT3G24360 | -1.168804 | 1.34E-04 | 2.41E-03 |
| AT3G25250 | 1.992632 | 1.01E-06 | 4.25E-05 |
| AT3G25510 | 1.157023 | 4.91E-05 | 1.10E-03 |
| AT3G25670 | -1.102618 | 1.75E-10 | 2.10E-08 |
| AT3G25730 | 1.430736 | 2.91E-13 | 6.08E-11 |
| AT3G25780 | 1.722407 | 3.04E-08 | 2.05E-06 |
| AT3G26500 | 1.132665 | 9.30E-09 | 7.31E-07 |
| AT3G26830 | 2.602226 | 3.70E-13 | 7.66E-11 |
| AT3G27400 | 1.469068 | 1.01E-08 | 7.78E-07 |
| AT3G28500 | 1.799586 | 4.92E-09 | 4.20E-07 |
| AT3G28540 | 1.016267 | 6.36E-06 | 2.01E-04 |
| AT3G28580 | 2.032464 | 7.69E-08 | 4.67E-06 |
| AT3G28930 | 1.442887 | 1.18E-19 | 7.63E-17 |
| AT3G29631 | 3.098037 | 7.75E-11 | 1.02E-08 |
| AT3G29639 | 4.013048 | 2.53E-13 | 5.40E-11 |
| AT3G32030 | 1.095691 | 1.12E-04 | 2.09E-03 |
| AT3G32040 | 1.699416 | 2.62E-05 | 6.54E-04 |
| AT3G32920 | 2.030293 | 8.58E-05 | 1.71E-03 |
| AT3G44690 | -3.44335 | 2.31E-12 | 3.92E-10 |
| AT3G45090 | -1.759682 | 1.92E-37 | 6.20E-34 |
| AT3G46280 | 1.025132 | 1.13E-04 | 2.11E-03 |
| AT3G46490 | -1.079351 | 1.99E-11 | 2.94E-09 |
| AT3G46920 | -1.193529 | 1.65E-10 | 2.00E-08 |
| AT3G48390 | -2.824576 | 2.71E-56 | 2.22E-52 |
| AT3G49970 | 1.642883 | 6.53E-06 | 2.05E-04 |
| AT3G50660 | -1.010373 | 1.93E-08 | 1.37E-06 |
| AT3G50760 | 1.064982 | 2.01E-05 | 5.27E-04 |
| AT3G50900 | 1.060198 | 1.75E-08 | 1.26E-06 |
| AT3G53960 | -1.145288 | 2.06E-07 | 1.09E-05 |
| AT3G55710 | -1.321725 | 2.02E-11 | 2.96E-09 |
| AT3G55920 | -1.135839 | 4.75E-14 | 1.18E-11 |
| AT3G57240 | 3.373956 | 7.68E-20 | 5.10E-17 |
| AT3G57460 | 1.52769 | 4.38E-05 | 9.91E-04 |
| AT3G58150 | 2.228854 | 1.85E-06 | 7.23E-05 |
| AT3G58270 | 1.249396 | 1.02E-09 | 1.05E-07 |
| AT3G58670 | -1.026168 | 7.08E-14 | 1.67E-11 |
| AT3G59220 | 1.297235 | 2.03E-06 | 7.83E-05 |
| AT3G59330 | -1.26463 | 1.46E-05 | 4.02E-04 |
| AT3G60070 | -1.687337 | 9.71E-16 | 3.37E-13 |
| AT3G60290 | -1.23809 | 4.74E-09 | 4.08E-07 |
| AT3G60710 | 1.498983 | 1.96E-04 | 3.26E-03 |
| AT3G62090 | 1.066856 | 1.49E-12 | 2.62E-10 |
| AT3G62460 | 1.932945 | 9.95E-25 | 1.50E-21 |
| AT4G00250 | 1.846308 | 8.26E-06 | 2.49E-04 |
| AT4G01700 | 1.003409 | 7.71E-06 | 2.35E-04 |
| AT4G01920 | 1.393038 | 5.49E-07 | 2.53E-05 |
| AT4G02330 | 2.130663 | 3.78E-13 | 7.76E-11 |
| AT4G02520 | 1.110253 | 6.65E-08 | 4.09E-06 |
| AT4G02660 | -2.208577 | 8.05E-11 | 1.05E-08 |
| AT4G03110 | -1.030689 | 7.53E-13 | 1.47E-10 |
| AT4G04223 | -1.717998 | 2.54E-14 | 6.59E-12 |
| AT4G04610 | 1.388323 | 1.53E-20 | 1.11E-17 |
| AT4G04800 | -1.16376 | 1.77E-13 | 3.99E-11 |
| AT4G05380 | 1.355059 | 2.72E-04 | 4.21E-03 |
| AT4G07995 | 2.155524 | 1.61E-06 | 6.42E-05 |
| AT4G08770 | 1.995675 | 2.32E-11 | 3.36E-09 |
| AT4G11130 | -1.03193 | 2.12E-09 | 2.00E-07 |
| AT4G11350 | 1.369606 | 1.09E-04 | 2.05E-03 |
| AT4G11470 | 1.024187 | 7.83E-04 | 9.71E-03 |
| AT4G11521 | 1.485179 | 9.50E-08 | 5.61E-06 |
| AT4G12470 | 1.581793 | 1.96E-13 | 4.35E-11 |
| AT4G12480 | 1.554309 | 8.10E-10 | 8.54E-08 |
| AT4G12490 | 1.502377 | 5.61E-08 | 3.51E-06 |
| AT4G12735 | 1.654102 | 1.22E-04 | 2.24E-03 |
| AT4G12830 | -1.347708 | 3.93E-24 | 4.93E-21 |
| AT4G14400 | 1.185585 | 7.22E-06 | 2.23E-04 |
| AT4G14640 | 1.227616 | 2.67E-04 | 4.16E-03 |
| AT4G15670 | -1.186692 | 2.67E-04 | 4.15E-03 |
| AT4G15680 | -1.428005 | 7.04E-05 | 1.47E-03 |
| AT4G15890 | 1.752645 | 1.06E-14 | 3.02E-12 |
| AT4G15975 | 1.167614 | 3.82E-05 | 8.87E-04 |
| AT4G16110 | -1.013656 | 1.31E-12 | 2.40E-10 |
| AT4G16250 | -2.768816 | 3.16E-36 | 6.48E-33 |
| AT4G16260 | 1.111253 | 6.90E-10 | 7.34E-08 |
| AT4G16857 | -3.125632 | 4.92E-24 | 5.85E-21 |
| AT4G17860 | -1.182216 | 6.37E-05 | 1.35E-03 |
| AT4G17980 | 1.30141 | 1.70E-04 | 2.93E-03 |
| AT4G18250 | 2.383436 | 4.81E-10 | 5.20E-08 |
| AT4G18253 | 2.060182 | 6.94E-15 | 2.03E-12 |
| AT4G18900 | 4.754665 | 2.68E-07 | 1.37E-05 |
| AT4G19430 | 1.043017 | 1.04E-04 | 1.99E-03 |
| AT4G20000 | 3.223473 | 1.03E-07 | 5.96E-06 |
| AT4G22390 | 2.932524 | 7.59E-20 | 5.10E-17 |

| | | | |
|---|---|---|---|
| AT3G06880 | -1.073258 | 4.06E-06 | 8.10E-05 |
| AT3G07105 | 1.495967 | 2.52E-05 | 3.95E-04 |
| AT3G07650 | 1.55203 | 2.48E-27 | 1.82E-24 |
| AT3G09390 | 1.002903 | 9.11E-07 | 2.24E-05 |
| AT3G09745 | 1.077987 | 2.52E-06 | 5.38E-05 |
| AT3G10570 | -1.025217 | 3.84E-08 | 1.34E-06 |
| AT3G11110 | -1.376782 | 2.14E-07 | 6.21E-06 |
| AT3G11773 | 1.033809 | 4.86E-04 | 4.81E-03 |
| AT3G12320 | -1.381109 | 1.66E-10 | 1.02E-08 |
| AT3G13450 | 1.200078 | 8.08E-17 | 1.54E-14 |
| AT3G13760 | 1.115745 | 4.53E-05 | 6.48E-04 |
| AT3G13784 | -1.596153 | 2.08E-06 | 4.54E-05 |
| AT3G14810 | -1.128249 | 1.22E-08 | 4.85E-07 |
| AT3G15354 | -1.03619 | 1.96E-09 | 9.19E-08 |
| AT3G15370 | 1.296961 | 2.99E-04 | 3.23E-03 |
| AT3G15440 | 2.197186 | 3.94E-18 | 9.07E-16 |
| AT3G15620 | 1.020637 | 2.11E-11 | 1.53E-09 |
| AT3G16175 | -1.35625 | 1.80E-04 | 2.12E-03 |
| AT3G16650 | -2.031898 | 4.62E-12 | 3.82E-10 |
| AT3G19030 | -2.153635 | 5.61E-16 | 9.52E-14 |
| AT3G19390 | 1.90547 | 1.57E-15 | 2.48E-13 |
| AT3G19450 | -1.133403 | 1.76E-15 | 2.72E-13 |
| AT3G19710 | -1.784994 | 2.40E-20 | 7.57E-18 |
| AT3G20810 | 1.906415 | 1.05E-21 | 4.39E-19 |
| AT3G21320 | 1.332094 | 3.45E-07 | 9.43E-06 |
| AT3G21670 | -1.523376 | 1.19E-17 | 2.58E-15 |
| AT3G21781 | 1.46589 | 1.41E-05 | 2.37E-04 |
| AT3G22160 | 1.188967 | 8.47E-09 | 3.46E-07 |
| AT3G22231 | -1.890832 | 4.80E-09 | 2.05E-07 |
| AT3G22235 | -1.677527 | 6.50E-09 | 2.70E-07 |
| AT3G22640 | 2.979604 | 5.41E-07 | 1.40E-05 |
| AT3G22740 | -1.833423 | 3.91E-10 | 2.23E-08 |
| AT3G23150 | 1.434072 | 1.27E-17 | 2.72E-15 |
| AT3G23240 | 3.238196 | 1.84E-10 | 1.12E-08 |
| AT3G23800 | 1.57284 | 2.71E-07 | 7.60E-06 |
| AT3G25900 | 1.401005 | 3.82E-16 | 6.54E-14 |
| AT3G26450 | -1.148002 | 1.63E-15 | 2.55E-13 |
| AT3G26570 | -1.27491 | 1.94E-13 | 1.98E-11 |
| AT3G27690 | -1.455076 | 7.45E-14 | 8.11E-12 |
| AT3G28930 | 1.033117 | 3.59E-11 | 2.46E-09 |
| AT3G29631 | 1.267876 | 1.45E-04 | 1.76E-03 |
| AT3G29639 | 2.741942 | 4.41E-08 | 1.51E-06 |
| AT3G30720 | 3.301799 | 5.35E-43 | 1.69E-39 |
| AT3G32030 | 1.070825 | 1.86E-04 | 2.17E-03 |
| AT3G43850 | 1.178209 | 1.00E-07 | 3.16E-06 |
| AT3G44450 | -1.259792 | 8.88E-05 | 1.16E-03 |
| AT3G44970 | -1.335568 | 4.05E-11 | 2.72E-09 |
| AT3G45090 | -1.475012 | 6.28E-29 | 5.14E-26 |
| AT3G45140 | -1.364977 | 1.68E-09 | 8.07E-08 |
| AT3G45300 | 1.213754 | 4.60E-13 | 4.44E-11 |
| AT3G46490 | -2.223288 | 3.17E-30 | 3.18E-27 |
| AT3G46880 | -1.445308 | 1.17E-04 | 1.47E-03 |
| AT3G47430 | -1.113833 | 9.38E-07 | 2.29E-05 |
| AT3G48100 | -1.385971 | 1.89E-11 | 1.38E-09 |
| AT3G48390 | -2.06785 | 5.52E-35 | 8.70E-32 |
| AT3G48460 | -1.376043 | 8.22E-16 | 1.33E-13 |
| AT3G48610 | -1.037273 | 5.51E-10 | 3.00E-08 |
| AT3G49420 | 1.118997 | 6.80E-08 | 2.23E-06 |
| AT3G49620 | 1.012255 | 2.85E-05 | 4.38E-04 |
| AT3G50685 | -1.070941 | 2.56E-09 | 1.17E-07 |
| AT3G51220 | -1.574821 | 1.19E-07 | 3.65E-06 |
| AT3G52072 | 1.013066 | 1.24E-08 | 4.89E-07 |
| AT3G52720 | -2.037679 | 1.68E-17 | 3.44E-15 |
| AT3G53830 | -1.412481 | 2.23E-10 | 1.34E-08 |
| AT3G54390 | -1.007403 | 4.35E-08 | 1.50E-06 |
| AT3G54500 | -1.129755 | 1.14E-09 | 5.79E-08 |
| AT3G54600 | -1.469382 | 1.64E-14 | 2.04E-12 |
| AT3G54730 | 3.731895 | 3.02E-09 | 1.35E-07 |
| AT3G55630 | -1.057104 | 2.52E-09 | 1.16E-07 |
| AT3G55920 | -1.30437 | 3.78E-17 | 7.59E-15 |
| AT3G56080 | -1.519745 | 7.30E-12 | 5.86E-10 |
| AT3G56290 | -1.172503 | 1.47E-07 | 4.41E-06 |
| AT3G57040 | -1.241563 | 3.25E-13 | 3.22E-11 |
| AT3G57260 | -1.200129 | 1.13E-05 | 1.96E-04 |
| AT3G58150 | 1.527897 | 4.65E-05 | 6.63E-04 |
| AT3G58270 | 1.219727 | 2.57E-09 | 1.17E-07 |
| AT3G58540 | 1.126768 | 4.29E-04 | 4.34E-03 |
| AT3G58990 | -1.161712 | 4.31E-08 | 1.49E-06 |
| AT3G59320 | -1.035057 | 3.67E-06 | 7.42E-05 |
| AT3G59330 | -1.352514 | 6.50E-06 | 1.22E-04 |
| AT3G59400 | -1.341461 | 3.91E-30 | 3.75E-27 |
| AT3G59930 | 1.244865 | 6.65E-05 | 9.03E-04 |
| AT3G60710 | 2.191163 | 1.32E-05 | 2.23E-04 |
| AT3G61510 | -1.716992 | 7.41E-05 | 9.95E-04 |
| AT3G62070 | -1.064635 | 8.88E-06 | 1.59E-04 |
| AT3G62090 | 1.853637 | 5.00E-33 | 6.91E-30 |
| AT3G62410 | -1.138412 | 8.45E-10 | 4.39E-08 |
| AT3G62460 | 1.46619 | 2.32E-15 | 3.53E-13 |
| AT4G00250 | 2.493418 | 2.10E-08 | 7.87E-07 |
| AT4G00430 | -1.00932 | 1.68E-13 | 1.72E-11 |
| AT4G01920 | 1.173915 | 9.67E-06 | 1.71E-04 |
| AT4G02330 | 1.157556 | 7.40E-06 | 1.36E-04 |
| AT4G02660 | -1.246841 | 6.56E-06 | 1.22E-04 |
| AT4G02850 | -1.314094 | 3.22E-10 | 1.87E-08 |
| AT4G03110 | -1.266674 | 9.13E-18 | 2.00E-15 |
| AT4G04180 | 1.128447 | 7.65E-10 | 4.02E-08 |
| AT4G04223 | 2.067695 | 5.98E-31 | 6.94E-28 |
| AT4G04800 | -1.457425 | 4.05E-19 | 1.13E-16 |
| AT4G04810 | -1.113878 | 8.86E-06 | 1.59E-04 |
| AT4G22470 | 2.268686 | 1.69E-10 | 2.04E-08 |
| AT4G22960 | 2.159554 | 6.29E-07 | 2.82E-05 |
| AT4G23140 | 1.888701 | 1.61E-07 | 8.85E-06 |
| AT4G23170 | 1.46015 | 2.14E-10 | 2.53E-08 |
| AT4G23210 | 1.015655 | 1.10E-05 | 3.17E-04 |
| AT4G23810 | 2.424305 | 3.20E-21 | 2.49E-18 |
| AT4G25120 | -2.227734 | 3.07E-17 | 1.42E-14 |
| AT4G25200 | 1.760507 | 1.08E-04 | 2.03E-03 |
| AT4G26120 | 1.41389 | 1.25E-08 | 9.47E-07 |
| AT4G27030 | -1.498341 | 4.23E-09 | 3.67E-07 |
| AT4G28040 | 1.362796 | 2.05E-06 | 7.91E-05 |
| AT4G28790 | 1.009683 | 4.29E-05 | 9.75E-04 |
| AT4G31500 | 1.592133 | 4.43E-22 | 3.85E-19 |
| AT4G31970 | 1.37823 | 2.14E-04 | 3.50E-03 |
| AT4G33070 | 1.241304 | 2.50E-06 | 9.32E-05 |
| AT4G34419 | 1.095144 | 6.30E-04 | 8.24E-03 |
| AT4G34950 | -1.183564 | 4.65E-08 | 2.98E-06 |
| AT4G35720 | 1.367443 | 3.88E-05 | 8.98E-04 |
| AT4G35985 | -1.01794 | 6.34E-08 | 3.93E-06 |
| AT4G37080 | -1.0026 | 2.57E-13 | 5.42E-11 |
| AT4G37370 | 1.278012 | 1.70E-07 | 9.21E-06 |
| AT4G38410 | 1.272453 | 1.08E-06 | 4.52E-05 |
| AT4G39800 | -1.005541 | 2.42E-24 | 3.42E-21 |
| AT4G39950 | 1.34503 | 4.82E-09 | 4.13E-07 |
| AT5G01540 | 1.536958 | 1.52E-09 | 1.49E-07 |
| AT5G02580 | 1.353038 | 1.83E-12 | 3.13E-10 |
| AT5G03520 | -1.593224 | 1.07E-14 | 3.02E-12 |
| AT5G03890 | 1.250868 | 2.53E-07 | 1.30E-05 |
| AT5G05130 | 1.485278 | 3.51E-18 | 1.84E-15 |
| AT5G05600 | 1.143599 | 6.03E-06 | 1.93E-04 |
| AT5G05890 | -1.001792 | 1.68E-12 | 2.90E-10 |
| AT5G06730 | 1.311025 | 4.34E-06 | 1.47E-04 |
| AT5G07010 | 1.036997 | 2.24E-14 | 5.89E-12 |
| AT5G08710 | -1.373139 | 1.27E-12 | 2.35E-10 |
| AT5G09570 | 4.620033 | 1.49E-23 | 1.60E-20 |
| AT5G10140 | 1.987359 | 6.27E-16 | 2.28E-13 |
| AT5G10250 | -2.469812 | 5.89E-15 | 1.80E-12 |
| AT5G11410 | 1.340186 | 7.52E-06 | 2.30E-04 |
| AT5G12330 | 1.052088 | 1.14E-08 | 8.73E-07 |
| AT5G13330 | 1.262694 | 3.70E-06 | 1.29E-04 |
| AT5G14650 | -1.110246 | 4.47E-07 | 2.13E-05 |
| AT5G15120 | 1.293109 | 3.18E-08 | 2.12E-06 |
| AT5G15380 | 1.281305 | 3.77E-04 | 5.51E-03 |
| AT5G16160 | -1.3099 | 1.33E-05 | 3.72E-04 |
| AT5G16570 | -1.115341 | 2.33E-10 | 2.74E-08 |
| AT5G17760 | 1.213231 | 2.19E-14 | 5.83E-12 |
| AT5G18470 | 1.809507 | 1.88E-15 | 6.16E-13 |
| AT5G22530 | 1.183233 | 5.14E-04 | 7.03E-03 |
| AT5G22570 | 1.094347 | 5.77E-04 | 7.70E-03 |
| AT5G24110 | 2.11875 | 4.49E-05 | 1.01E-03 |
| AT5G24206 | 1.089973 | 4.93E-04 | 6.79E-03 |
| AT5G24240 | 2.238365 | 5.11E-08 | 3.24E-06 |
| AT5G24280 | 1.743242 | 3.64E-17 | 1.61E-14 |
| AT5G24640 | 2.913846 | 3.30E-10 | 3.67E-08 |
| AT5G25130 | -1.062944 | 8.61E-09 | 6.87E-07 |
| AT5G25470 | 1.197753 | 8.73E-05 | 1.72E-03 |
| AT5G26270 | 1.353301 | 5.77E-11 | 7.90E-09 |
| AT5G27220 | -1.955675 | 7.75E-08 | 4.69E-06 |
| AT5G27230 | -1.023275 | 3.78E-04 | 5.52E-03 |
| AT5G27240 | -1.315347 | 2.16E-05 | 5.59E-04 |
| AT5G35480 | 2.125603 | 2.89E-15 | 9.33E-13 |
| AT5G35490 | 1.781369 | 8.04E-08 | 4.86E-06 |
| AT5G35740 | -1.726548 | 2.31E-13 | 5.01E-11 |
| AT5G35970 | -1.104028 | 6.08E-10 | 6.51E-08 |
| AT5G37530 | -1.683242 | 2.92E-10 | 3.28E-08 |
| AT5G38140 | -1.110574 | 8.67E-10 | 9.02E-08 |
| AT5G38240 | 1.481549 | 8.15E-07 | 3.56E-05 |
| AT5G38550 | 1.060641 | 3.90E-05 | 9.00E-04 |
| AT5G38900 | 1.096359 | 9.95E-05 | 1.92E-03 |
| AT5G39120 | 1.020253 | 8.05E-04 | 9.92E-03 |
| AT5G39850 | 1.139907 | 7.66E-13 | 1.48E-10 |
| AT5G39890 | 1.230184 | 1.01E-07 | 5.88E-06 |
| AT5G40270 | -1.315995 | 3.52E-17 | 1.59E-14 |
| AT5G40990 | 1.820408 | 1.01E-04 | 1.93E-03 |
| AT5G41761 | 1.951428 | 2.87E-18 | 1.54E-15 |
| AT5G42900 | 1.149247 | 2.13E-09 | 2.00E-07 |
| AT5G44210 | 1.07523 | 9.59E-09 | 7.46E-07 |
| AT5G44420 | 1.298627 | 2.34E-05 | 5.94E-04 |
| AT5G44585 | 2.135402 | 1.35E-08 | 1.01E-06 |
| AT5G44620 | -1.565336 | 1.43E-05 | 3.94E-04 |
| AT5G44680 | -1.077032 | 1.21E-23 | 1.37E-20 |
| AT5G45990 | 1.312372 | 4.59E-05 | 1.03E-03 |
| AT5G46050 | 2.155101 | 1.70E-14 | 4.68E-12 |
| AT5G47220 | 1.677181 | 2.06E-14 | 5.60E-12 |
| AT5G47960 | 1.290314 | 5.02E-07 | 2.34E-05 |
| AT5G48070 | 1.604396 | 5.24E-09 | 4.41E-07 |
| AT5G48490 | -1.073496 | 4.60E-05 | 1.04E-03 |
| AT5G48657 | 1.063153 | 5.39E-06 | 1.77E-04 |
| AT5G49520 | 1.129897 | 4.20E-07 | 2.00E-05 |
| AT5G50335 | 1.193638 | 5.13E-15 | 1.60E-12 |
| AT5G50915 | -1.106118 | 1.20E-05 | 3.40E-04 |
| AT5G50950 | -1.211747 | 2.33E-17 | 1.10E-14 |
| AT5G51440 | 1.299422 | 5.06E-05 | 1.12E-03 |
| AT5G51920 | 1.208309 | 1.18E-04 | 2.18E-03 |
| AT5G52070 | 1.848997 | 2.71E-36 | 6.12E-33 |
| AT5G52390 | 1.507786 | 9.74E-06 | 2.88E-04 |
| AT5G52930 | 1.05054 | 8.68E-06 | 2.60E-04 |
| AT5G52940 | 2.747104 | 4.88E-19 | 2.98E-16 |
| AT5G53230 | 3.842175 | 2.85E-06 | 1.04E-04 |

| | | | |
|---|---|---|---|
| AT4G04840 | -1.739146 | 2.10E-18 | 5.04E-16 |
| AT4G05110 | -1.626098 | 6.03E-06 | 1.14E-04 |
| AT4G06746 | 1.269802 | 8.01E-09 | 3.29E-07 |
| AT4G07820 | 1.01722 | 9.76E-05 | 1.26E-03 |
| AT4G07995 | 1.225111 | 2.13E-04 | 2.43E-03 |
| AT4G08770 | 1.181242 | 8.88E-06 | 1.59E-04 |
| AT4G08870 | -1.457743 | 8.10E-16 | 1.32E-13 |
| AT4G09350 | -2.193911 | 1.26E-14 | 1.66E-12 |
| AT4G09650 | -1.069382 | 1.03E-09 | 5.26E-08 |
| AT4G09750 | -1.030778 | 5.34E-09 | 2.26E-07 |
| AT4G11521 | 1.04572 | 3.69E-05 | 5.44E-04 |
| AT4G11910 | 1.24692 | 5.70E-07 | 1.47E-05 |
| AT4G12030 | -1.101791 | 3.18E-07 | 8.77E-06 |
| AT4G12470 | 1.839584 | 8.09E-18 | 1.80E-15 |
| AT4G12480 | 1.129364 | 1.81E-06 | 4.00E-05 |
| AT4G12830 | -1.464271 | 1.91E-27 | 1.45E-24 |
| AT4G13575 | -1.238211 | 1.07E-07 | 3.33E-06 |
| AT4G13770 | -1.579869 | 3.34E-18 | 7.85E-16 |
| AT4G14130 | 1.426367 | 2.47E-09 | 1.14E-07 |
| AT4G14548 | 1.062978 | 2.11E-05 | 3.38E-04 |
| AT4G15630 | -1.009509 | 1.05E-05 | 1.85E-04 |
| AT4G15890 | 2.000325 | 5.00E-19 | 1.38E-16 |
| AT4G16250 | -2.060127 | 1.40E-23 | 7.93E-21 |
| AT4G16857 | -2.792148 | 1.40E-21 | 5.60E-19 |
| AT4G16880 | -1.481262 | 2.24E-07 | 6.47E-06 |
| AT4G16980 | -1.016175 | 4.70E-31 | 6.10E-28 |
| AT4G17860 | -2.064675 | 9.15E-08 | 2.90E-06 |
| AT4G18150 | 2.20507 | 2.38E-05 | 3.76E-04 |
| AT4G18900 | 1.6108 | 9.88E-05 | 1.27E-03 |
| AT4G19120 | 1.199946 | 5.17E-09 | 2.19E-07 |
| AT4G19420 | -1.080874 | 2.11E-10 | 1.27E-08 |
| AT4G19430 | 1.27171 | 1.76E-05 | 2.90E-04 |
| AT4G19820 | -1.082877 | 9.61E-06 | 1.71E-04 |
| AT4G19950 | 1.157582 | 1.12E-06 | 2.65E-05 |
| AT4G21500 | 1.022206 | 1.50E-05 | 2.51E-04 |
| AT4G21760 | -1.473619 | 1.81E-05 | 2.97E-04 |
| AT4G21870 | -1.236695 | 1.77E-06 | 3.93E-05 |
| AT4G22390 | 2.310365 | 8.54E-14 | 9.11E-12 |
| AT4G22470 | 1.885518 | 1.05E-08 | 4.24E-07 |
| AT4G22570 | -1.103022 | 4.40E-11 | 2.95E-09 |
| AT4G23260 | -1.013693 | 1.23E-05 | 2.11E-04 |
| AT4G23670 | -1.061579 | 5.60E-07 | 1.44E-05 |
| AT4G23810 | 1.546116 | 1.50E-10 | 9.39E-09 |
| AT4G24040 | 1.120435 | 2.60E-12 | 2.27E-10 |
| AT4G24350 | -1.461515 | 9.27E-19 | 2.47E-16 |
| AT4G24972 | -1.04722 | 1.62E-05 | 2.68E-04 |
| AT4G25050 | -1.106086 | 7.76E-17 | 1.49E-14 |
| AT4G25120 | -1.3912 | 1.18E-09 | 5.96E-08 |
| AT4G25420 | -1.116151 | 2.72E-05 | 4.21E-04 |
| AT4G25580 | 3.469073 | 3.45E-22 | 1.59E-19 |
| AT4G27030 | -2.475211 | 1.71E-18 | 4.25E-16 |
| AT4G27290 | 1.339915 | 3.70E-05 | 5.45E-04 |
| AT4G27700 | -1.195481 | 4.13E-13 | 4.03E-11 |
| AT4G27970 | 1.117184 | 9.78E-06 | 1.73E-04 |
| AT4G28040 | 2.025758 | 3.96E-11 | 2.68E-09 |
| AT4G28680 | -1.079416 | 8.70E-05 | 1.14E-03 |
| AT4G30110 | -1.069449 | 3.30E-07 | 9.04E-06 |
| AT4G30290 | 1.728493 | 3.25E-21 | 1.25E-18 |
| AT4G33150 | 1.083046 | 1.70E-12 | 1.52E-10 |
| AT4G33550 | -1.299195 | 2.40E-05 | 3.78E-04 |
| AT4G33980 | 2.026893 | 1.48E-48 | 8.15E-45 |
| AT4G34400 | -1.052351 | 5.06E-04 | 4.98E-03 |
| AT4G34580 | 1.006282 | 1.00E-07 | 3.15E-06 |
| AT4G34610 | -1.44483 | 5.27E-14 | 5.91E-12 |
| AT4G35770 | 1.474106 | 1.67E-14 | 2.04E-12 |
| AT4G36410 | 1.068212 | 2.95E-07 | 8.19E-06 |
| AT4G36670 | 1.235884 | 2.45E-14 | 2.89E-12 |
| AT4G36770 | -1.082003 | 5.96E-04 | 5.70E-03 |
| AT4G37390 | 1.447895 | 8.16E-19 | 2.20E-16 |
| AT4G37925 | -1.484814 | 3.48E-14 | 4.00E-12 |
| AT4G38410 | 1.640347 | 1.12E-09 | 5.69E-08 |
| AT4G38580 | 1.004556 | 2.38E-12 | 2.10E-10 |
| AT4G38840 | -1.371637 | 4.85E-17 | 9.55E-15 |
| AT4G38860 | -1.343086 | 3.75E-22 | 1.69E-19 |
| AT4G39070 | 1.323191 | 1.23E-08 | 4.88E-07 |
| AT4G39800 | -1.142946 | 2.79E-30 | 2.94E-27 |
| AT5G01015 | -1.572054 | 1.64E-10 | 1.01E-08 |
| AT5G01210 | 1.130942 | 3.03E-13 | 3.02E-11 |
| AT5G01215 | 1.152403 | 1.38E-11 | 1.05E-09 |
| AT5G01540 | 1.338256 | 5.47E-08 | 1.83E-06 |
| AT5G01790 | 1.187026 | 8.26E-10 | 4.32E-08 |
| AT5G02580 | 1.149015 | 1.44E-09 | 7.07E-08 |
| AT5G02750 | 1.144496 | 8.37E-10 | 4.37E-08 |
| AT5G02865 | -1.388626 | 2.32E-08 | 8.58E-07 |
| AT5G03260 | -1.339198 | 7.44E-10 | 3.96E-08 |
| AT5G03350 | -1.043246 | 3.20E-05 | 4.85E-04 |
| AT5G03520 | -1.456397 | 4.93E-13 | 4.73E-11 |
| AT5G03730 | 1.002777 | 6.88E-07 | 1.73E-05 |
| AT5G03890 | 1.036883 | 1.04E-05 | 1.82E-04 |
| AT5G04110 | -2.14493 | 1.39E-20 | 4.52E-18 |
| AT5G04310 | 1.506767 | 1.06E-10 | 6.74E-09 |
| AT5G05060 | 1.078852 | 3.15E-19 | 8.91E-17 |
| AT5G05130 | 1.632925 | 5.93E-22 | 2.57E-19 |
| AT5G05270 | -1.595706 | 7.92E-11 | 5.17E-09 |
| AT5G05580 | -1.5598 | 2.06E-16 | 3.75E-14 |
| AT5G05880 | 1.023836 | 2.86E-06 | 6.00E-05 |
| AT5G05965 | 1.198512 | 1.59E-06 | 3.58E-05 |
| AT5G06570 | 1.434384 | 5.77E-09 | 2.41E-07 |
| AT5G06620 | 1.043016 | 9.37E-07 | 2.29E-05 |
| AT5G53240 | 2.160077 | 5.77E-05 | 1.24E-03 |
| AT5G53750 | 1.274321 | 6.68E-07 | 2.98E-05 |
| AT5G53820 | 1.42581 | 1.41E-05 | 3.89E-04 |
| AT5G54190 | -1.419331 | 2.53E-06 | 9.39E-05 |
| AT5G54550 | 4.654554 | 2.79E-07 | 1.42E-05 |
| AT5G54560 | 4.439655 | 9.32E-08 | 5.55E-06 |
| AT5G55450 | 1.055659 | 4.28E-06 | 1.46E-04 |
| AT5G55490 | 1.296846 | 3.36E-04 | 5.02E-03 |
| AT5G55570 | -2.147561 | 1.67E-07 | 9.07E-06 |
| AT5G56370 | 2.334333 | 3.40E-37 | 9.18E-34 |
| AT5G56810 | 2.650849 | 3.82E-17 | 1.66E-14 |
| AT5G58610 | 1.759177 | 1.65E-08 | 1.21E-06 |
| AT5G58660 | 1.194667 | 1.66E-08 | 1.21E-06 |
| AT5G60250 | 1.809206 | 3.00E-12 | 5.02E-10 |
| AT5G60530 | 1.559119 | 3.20E-06 | 1.14E-04 |
| AT5G60610 | 2.782641 | 2.72E-21 | 2.19E-18 |
| AT5G61520 | 1.057296 | 1.57E-22 | 1.42E-19 |
| AT5G61610 | 1.112953 | 1.91E-05 | 5.03E-04 |
| AT5G61730 | 1.14113 | 5.76E-04 | 7.70E-03 |
| AT5G62100 | -1.279698 | 6.05E-09 | 5.01E-07 |
| AT5G63650 | 1.42443 | 8.95E-14 | 2.08E-11 |
| AT5G64060 | 1.667621 | 1.18E-07 | 6.69E-06 |
| AT5G64120 | 1.093673 | 2.92E-07 | 1.47E-05 |
| AT5G64850 | -1.038222 | 5.58E-06 | 1.80E-04 |
| AT5G65080 | 1.082062 | 2.22E-04 | 3.58E-03 |
| AT5G65600 | 1.089486 | 1.13E-04 | 2.11E-03 |
| AT5G66630 | 1.142201 | 2.00E-06 | 7.73E-05 |
| AT5G67060 | 1.03995 | 1.53E-04 | 2.69E-03 |
| ATCG01180 | -1.050905 | 7.92E-09 | 6.43E-07 |

| | | | |
|---|---|---|---|
| AT5G07010 | 1.053572 | 1.75E-14 | 2.13E-12 |
| AT5G07440 | 1.131159 | 1.90E-12 | 1.69E-10 |
| AT5G07460 | -1.295774 | 4.98E-14 | 5.61E-12 |
| AT5G07560 | -1.186578 | 2.62E-04 | 2.91E-03 |
| AT5G07690 | -1.508071 | 1.75E-07 | 5.17E-06 |
| AT5G08030 | -1.815899 | 6.34E-05 | 8.67E-04 |
| AT5G08130 | -1.03777 | 3.30E-06 | 6.76E-05 |
| AT5G08710 | -1.338716 | 3.47E-12 | 2.94E-10 |
| AT5G09300 | -1.196167 | 1.93E-09 | 9.07E-08 |
| AT5G09570 | 3.955044 | 8.66E-18 | 1.91E-15 |
| AT5G09995 | -1.453683 | 1.91E-20 | 6.11E-18 |
| AT5G10140 | 1.425602 | 1.10E-09 | 5.59E-08 |
| AT5G10250 | -1.980915 | 8.12E-12 | 6.45E-10 |
| AT5G10760 | -1.166354 | 1.12E-05 | 1.94E-04 |
| AT5G13220 | 1.045171 | 1.17E-08 | 4.66E-07 |
| AT5G13330 | 1.051621 | 5.10E-05 | 7.17E-04 |
| AT5G13770 | -1.24754 | 1.25E-16 | 2.35E-14 |
| AT5G14200 | -1.493568 | 2.74E-13 | 2.75E-11 |
| AT5G14470 | 1.095993 | 1.53E-06 | 3.48E-05 |
| AT5G14920 | 1.278693 | 1.63E-14 | 2.04E-12 |
| AT5G15120 | 1.736708 | 4.01E-13 | 3.94E-11 |
| AT5G15850 | -1.349445 | 2.90E-09 | 1.31E-07 |
| AT5G16160 | -1.397964 | 5.61E-06 | 1.08E-04 |
| AT5G16190 | -1.119955 | 3.38E-05 | 5.07E-04 |
| AT5G16570 | -1.309685 | 6.60E-13 | 6.22E-11 |
| AT5G17230 | -1.161315 | 2.64E-20 | 7.98E-18 |
| AT5G17300 | -1.044406 | 3.87E-09 | 1.69E-07 |
| AT5G17860 | 1.512539 | 2.86E-09 | 1.29E-07 |
| AT5G18020 | -1.096572 | 1.65E-07 | 4.90E-06 |
| AT5G18030 | -1.351065 | 8.39E-10 | 4.37E-08 |
| AT5G18080 | -1.073657 | 6.54E-09 | 2.71E-07 |
| AT5G18470 | 1.590321 | 5.73E-13 | 5.45E-11 |
| AT5G18660 | -1.448629 | 1.57E-12 | 1.41E-10 |
| AT5G19230 | 1.310952 | 3.59E-09 | 1.58E-07 |
| AT5G19500 | -1.381595 | 6.60E-16 | 1.10E-13 |
| AT5G20935 | -1.000817 | 1.29E-06 | 2.98E-05 |
| AT5G21100 | -1.441012 | 1.36E-11 | 1.03E-09 |
| AT5G22390 | -1.85182 | 7.88E-13 | 7.34E-11 |
| AT5G22500 | 1.626513 | 8.57E-15 | 1.17E-12 |
| AT5G23010 | -1.42316 | 3.02E-11 | 2.11E-09 |
| AT5G23050 | 1.172906 | 8.84E-12 | 6.97E-10 |
| AT5G23060 | -1.10382 | 5.96E-16 | 9.98E-14 |
| AT5G23240 | 2.068937 | 3.39E-24 | 1.97E-21 |
| AT5G23700 | 1.325356 | 3.46E-08 | 1.23E-06 |
| AT5G24105 | -1.372957 | 1.53E-05 | 2.54E-04 |
| AT5G24120 | -1.332356 | 2.62E-12 | 2.27E-10 |
| AT5G24150 | -1.129665 | 8.55E-05 | 1.12E-03 |
| AT5G24280 | 1.060873 | 7.24E-08 | 2.36E-06 |
| AT5G24420 | -1.36907 | 1.73E-12 | 1.54E-10 |
| AT5G24640 | 2.652154 | 2.15E-09 | 1.00E-07 |
| AT5G25130 | -1.831106 | 1.51E-18 | 3.79E-16 |
| AT5G25350 | 1.017074 | 1.58E-08 | 6.08E-07 |
| AT5G26200 | -1.265018 | 9.09E-07 | 2.23E-05 |
| AT5G26270 | 4.23459 | 3.28E-95 | 7.25E-91 |
| AT5G27360 | -1.153318 | 5.46E-10 | 2.98E-08 |
| AT5G30495 | 1.012413 | 5.54E-09 | 2.33E-07 |
| AT5G35740 | -2.142131 | 1.54E-17 | 3.18E-15 |
| AT5G37530 | -1.151708 | 9.89E-07 | 2.39E-05 |
| AT5G38005 | 2.197219 | 4.49E-14 | 5.08E-12 |
| AT5G38430 | -1.859441 | 4.04E-29 | 3.43E-26 |
| AT5G39160 | 1.28286 | 4.09E-06 | 8.15E-05 |
| AT5G39190 | 1.224777 | 2.72E-09 | 1.23E-07 |
| AT5G39410 | 1.473888 | 2.51E-20 | 7.70E-18 |
| AT5G39520 | 1.726909 | 1.27E-10 | 8.02E-09 |
| AT5G39860 | -2.216276 | 7.65E-14 | 8.24E-12 |
| AT5G39890 | 1.858237 | 9.45E-15 | 1.26E-12 |
| AT5G40540 | 1.048863 | 1.07E-09 | 5.47E-08 |
| AT5G41761 | 1.207071 | 2.12E-08 | 7.91E-07 |
| AT5G42530 | -1.269731 | 7.51E-08 | 2.44E-06 |
| AT5G42900 | 3.441232 | 4.87E-62 | 5.38E-58 |
| AT5G43380 | 1.056634 | 1.66E-06 | 3.72E-05 |
| AT5G44190 | -1.088014 | 9.01E-11 | 5.80E-09 |
| AT5G44440 | 1.926008 | 1.04E-06 | 2.49E-05 |
| AT5G44585 | 1.450201 | 7.87E-06 | 1.43E-04 |
| AT5G44620 | -1.901376 | 1.11E-06 | 2.62E-05 |
| AT5G44680 | -1.514102 | 2.35E-43 | 8.66E-40 |
| AT5G44910 | 1.008341 | 2.11E-05 | 3.37E-04 |
| AT5G45650 | -1.1122 | 9.62E-11 | 6.18E-09 |
| AT5G45820 | -1.39196 | 3.55E-38 | 7.85E-35 |
| AT5G45950 | -1.226147 | 1.02E-11 | 7.90E-10 |
| AT5G46050 | 1.293236 | 4.97E-07 | 1.30E-05 |
| AT5G46490 | -1.057078 | 1.02E-05 | 1.79E-04 |
| AT5G47590 | 1.666604 | 3.06E-13 | 3.04E-11 |
| AT5G48180 | 1.67803 | 1.36E-17 | 2.90E-15 |
| AT5G48250 | 1.371262 | 1.55E-24 | 9.25E-22 |
| AT5G48490 | -2.159055 | 1.67E-11 | 1.24E-09 |
| AT5G49520 | 1.044877 | 2.80E-06 | 5.92E-05 |
| AT5G49690 | 1.596063 | 4.84E-11 | 3.24E-09 |
| AT5G49740 | -1.145634 | 6.38E-07 | 1.63E-05 |
| AT5G50335 | 1.255068 | 2.89E-16 | 5.10E-14 |
| AT5G50740 | -1.686201 | 4.41E-23 | 2.37E-20 |
| AT5G50800 | -1.877381 | 1.96E-05 | 3.19E-04 |
| AT5G50915 | -1.764458 | 1.12E-09 | 5.69E-08 |
| AT5G50950 | -1.762307 | 1.56E-33 | 2.29E-30 |
| AT5G51480 | 2.186385 | 1.54E-11 | 1.15E-09 |
| AT5G51850 | -1.182485 | 7.73E-07 | 1.93E-05 |
| AT5G52070 | 2.048539 | 6.65E-45 | 2.94E-41 |
| AT5G52170 | 1.109428 | 2.14E-07 | 6.20E-06 |
| AT5G52910 | 1.487392 | 5.74E-12 | 4.70E-10 |

| | | | |
|---|---|---|---|
| AT5G52940 | 2.457779 | 3.78E-16 | 6.52E-14 |
| AT5G53750 | 1.366227 | 1.38E-07 | 4.16E-06 |
| AT5G54190 | -2.053629 | 2.87E-10 | 1.68E-08 |
| AT5G54470 | -1.153192 | 5.81E-05 | 8.04E-04 |
| AT5G54550 | 3.972006 | 9.32E-07 | 2.28E-05 |
| AT5G54560 | 3.52333 | 1.37E-06 | 3.16E-05 |
| AT5G55570 | -1.558521 | 4.66E-06 | 9.18E-05 |
| AT5G55720 | -1.733219 | 1.04E-06 | 2.49E-05 |
| AT5G56370 | 1.702235 | 7.19E-21 | 2.52E-18 |
| AT5G56810 | 1.530811 | 1.07E-07 | 3.33E-06 |
| AT5G56850 | -1.649146 | 1.14E-29 | 1.05E-26 |
| AT5G56870 | 1.133998 | 1.17E-11 | 9.03E-10 |
| AT5G57110 | 1.002785 | 3.64E-08 | 1.28E-06 |
| AT5G57190 | -1.213567 | 8.48E-08 | 2.71E-06 |
| AT5G57240 | 1.104055 | 4.55E-13 | 4.41E-11 |
| AT5G57345 | -1.139455 | 3.50E-08 | 1.24E-06 |
| AT5G57640 | 1.711107 | 6.53E-10 | 3.50E-08 |
| AT5G58610 | 1.326339 | 3.79E-06 | 7.61E-05 |
| AT5G58660 | 1.643201 | 3.24E-14 | 3.74E-12 |
| AT5G59130 | -1.137138 | 2.18E-07 | 6.30E-06 |
| AT5G59570 | 1.822796 | 4.42E-15 | 6.38E-13 |
| AT5G59750 | -1.063158 | 1.26E-11 | 9.61E-10 |
| AT5G60100 | 2.015761 | 1.24E-12 | 1.13E-10 |
| AT5G60250 | 1.456159 | 4.70E-09 | 2.02E-07 |
| AT5G60610 | 1.394976 | 2.30E-07 | 6.63E-06 |
| AT5G61160 | 1.027597 | 1.14E-05 | 1.96E-04 |
| AT5G61270 | -1.046435 | 2.24E-10 | 1.34E-08 |
| AT5G61380 | 1.539109 | 1.65E-26 | 1.17E-23 |
| AT5G61610 | 1.056329 | 5.12E-05 | 7.19E-04 |
| AT5G62040 | -1.963261 | 1.07E-05 | 1.87E-04 |
| AT5G62100 | -1.021702 | 1.07E-06 | 2.56E-05 |
| AT5G62140 | -1.026119 | 1.12E-05 | 1.94E-04 |
| AT5G62430 | -1.68046 | 1.98E-11 | 1.44E-09 |
| AT5G62920 | -2.443977 | 1.78E-11 | 1.32E-09 |
| AT5G63350 | 1.505794 | 1.94E-09 | 9.09E-08 |
| AT5G63650 | 2.227414 | 1.89E-30 | 2.09E-27 |
| AT5G63810 | 1.10429 | 2.25E-14 | 2.69E-12 |
| AT5G63980 | -1.576061 | 2.51E-16 | 4.54E-14 |
| AT5G64850 | -2.065014 | 8.80E-15 | 1.19E-12 |
| AT5G65010 | -1.11585 | 3.51E-10 | 2.01E-08 |
| AT5G65080 | 1.375133 | 3.43E-05 | 5.12E-04 |
| AT5G65140 | -1.075387 | 3.13E-06 | 6.47E-05 |
| AT5G65850 | 1.072241 | 7.55E-05 | 1.01E-03 |
| AT5G65890 | -1.075595 | 9.19E-12 | 7.22E-10 |
| AT5G66580 | -1.134247 | 1.20E-05 | 2.06E-04 |
| AT5G66630 | 1.665139 | 8.32E-11 | 5.40E-09 |
| AT5G67060 | 1.016686 | 2.68E-04 | 2.97E-03 |
| AT5G67370 | -1.128102 | 1.35E-11 | 1.03E-09 |
| ATMG00020 | 1.06988 | 2.53E-08 | 9.29E-07 |
